# Brain injury environment critically influences the connectivity of transplanted neurons

**DOI:** 10.1101/2021.12.13.472270

**Authors:** S. Grade, J. Thomas, Y. Zarb, M. Thorwirth, Karl-Klaus Conzelmann, Stefanie M. Hauck, Magdalena Götz

**Affiliations:** Biomedical Center, Ludwig-Maximilians University Munich, 82152 Planegg-Martinsried, Germany; Institute of Stem Cell Research, Helmholtz Center Munich, German Center for Environmental Health, 82152 Planegg-Martinsried, Germany; Graduate School of Systemic Neuroscience, Ludwig-Maximilians University Munich, 82152 Planegg-Martinsried, Germany; Max von Pettenkofer Institute Virology, Medical faculty and Gene Center, Ludwig-Maximilians University Munich, 81377 Munich, Germany; Research Unit Protein Science and Metabolomics and Proteomics Core, Helmholtz Center Munich, German Center for Environmental Health, Neuherberg, 85764, Germany; SYNERGY, Excellence Cluster of Systems Neurology, Ludwig-Maximilians University Munich, 81377 Munich, Germany

## Abstract

Cell transplantation is a promising approach for the reconstruction of neuronal circuits after brain damage. Transplanted neurons integrate with remarkable specificity into circuitries of the mouse cerebral cortex affected by neuronal ablation. However, it remains unclear how neurons perform in a local environment undergoing reactive gliosis, inflammation, macrophage infiltration and scar formation, as in traumatic brain injury (TBI). To elucidate this, we transplanted cells from the embryonic mouse cerebral cortex into TBI-injured, inflamed-only, or intact cortex of adult mice. Brain-wide quantitative connectomics unraveled graft inputs from correct regions across the brain in all conditions, with pronounced quantitative differences: scarce in intact and inflamed brain, versus exuberant after trauma. In the latter, excessive synapse pruning follows the initial overshoot of connectivity resulting in only a few connections left. Proteomic profiling identifies candidate molecules involved in the synaptic yield, a pivotal parameter to tailor for functional restoration of neuronal circuits.

**Teaser:** Neuronal grafts in a brain area affected by trauma receive excessive yet mostly vulnerable inputs from host circuits.

## Introduction

The adult mammalian brain poorly regenerates in the aftermath of an injury and thus strategies of neuronal transplantation have been pursued, to rebuild circuits and restore behavioral function (1–4). Quantitative connectomics has enabled a fine comparison of the connections developed by transplants to their endogenous counterparts (5–7). Previous analysis in the mouse cortex has shown that new neurons mature and connect with the host brain in a remarkably correct manner, with high specificity both in their synaptic integration and in the response to external stimuli (5). The injury, in this case, consisted on the selective ablation of a cohort of neurons in the cortex by inducing apoptosis in the targeted neurons, and implied little inflammatory reaction, no glial scar or brain-blood barrier breakdown (8).

While these findings are encouraging, it remains open whether accurate connectivity is achieved in more clinically-relevant injuries like a TBI, which provide a distinctive cellular and molecular milieu. While several transplantation studies described a bystander effect via neuroprotection and immunomodulation in TBI (9–12) still little is known about neuronal replacement and brain-wide connectivity in this injury. This has been probed only using grafts with low amounts of mature neurons and lacking full and quantitative connectivity analysis (13). Herein, we analyzed graft connectivity in a mouse model of penetrating TBI. TBI results in severe reactive gliosis and scar formation, with infiltration of phagocytic macrophages and accumulation of high content of inflammatory molecules (14–16). Molecular and cellular components of an injured brain parenchyma have been implicated in synapse pruning (17,18). One may hence predict deficits in the initial synaptic integration, as synapse formation may concur with uncontrollable pruning. Microglial cells, in particular, are critical for brain wiring and synapse pruning (19) and are the first responders to brain injury, swiftly adopting an activated phenotype and initiating neuroinflammation (20,21). To examine their contribution, we utilized a systemic injection of lipopolysaccharide (LPS), a well-documented inflammatory stimulus as an additional condition complementing the TBI. LPS activates TLR4 receptor which is specifically expressed by microglial cells in the rodent brain (22). To achieve the least damaged environment we also transplanted into the intact cerebral cortex, only inflicted by the thin transplantation needle, as a 3^rd^ paradigm.

Importantly, we also set out to study the maintenance of initial connectivity. A phase of active synapse remodeling with a net gain was observed by live imaging in the neurites of transplanted neurons during the first month (5). This synapse turnover then gradually reached the basal turnover rate observed in the cortex of adult mice (23) at about 2-3 months after transplantation. However, the balance may be tilted towards an excessive elimination of new synapses in an environment characterized by a reactive and inflammatory state with large amount of pruning cells. The stability of the initial connectivity in TBI conditions remains unknown. Long-term connectivity is crucial for the successful outcome of a neuronal replacement therapy and needs to be ensured prior to clinical trials. Here, we investigated the role of the host microenvironment in initial and long-term connectivity of neuronal transplants by using rabies virus (RABV) mediated tracing and brain-wide quantitative connectomics, and leveraged proteomics to identify candidate molecules involved in the connectivity differences.

## Results

### Transplanted neurons develop complex morphologies and synaptic protrusions in cortical stab injury

To produce a brain injury in the cerebral cortex of the adult mouse brain we inflicted a stab injury or so-called stab wound (SW) within the primary visual cortex. The choice of the injury model rests on its high reproducibility and extensive knowledge about the temporal dynamics of wound healing, reactive gliosis, scar formation, as well as transcriptome and proteome (15,16,24– 26). Donor cells from mouse embryonic cortex expressing GFP or RFP were transplanted into the center of the incision a week later and analyzed 5 days and 5 weeks post-transplantation (dpt/wpt; Fig. 1A, B). Early, at 5dpt, transplanted cells express the immature neuronal marker doublecortin (Dcx; Fig. 1C). By 5wpt, the graft remained confined to the site of transplantation and injury, with no or little cell dispersion, and cells had developed complex arborizations of their neurites (Fig. 1D-F). Cux1 immunolabeling demonstrated that a majority of transplanted neurons displayed cortical upper layer identity (Fig. 1E and S3E; with 70.27% of the transplanted cells expressing Cux1). These observations are in line with those in the neuronal ablation model with low inflammation (5). Note that Cux1 is also expressed in reelin+ interneurons at early postnatal stages (27). Nevertheless both the morphology observed herein (also by live imaging *in vivo* in (5)) together with Cux1 and additionally Satb2 staining suggests that most transplanted neurons are pyramidal neurons, and at least some of callosal projection neuron identity (Satb2+; Fig. S3F). Inspection at high magnification shows that a complex network of axons and dendrites has outgrown from single transplanted neurons with appreciable density of axonal boutons and dendritic spines throughout their length (Fig. 1F, insets; Movie S1). Spontaneous cell-to-cell fusion events can occur between transplanted cells and central nervous system neurons although these are extremely rare (28,29). We tested for the occurrence of cell fusion by using Emx1-Cre/GFP donor cells (30,31) and tdTomato reporter mice as hosts (32) (Fig. S1A). Cell fusion upon transplantation would result in Cre-mediated recombination and expression of tdTomato. We observed no tdTomato neurons and no GFP+/TdTomato+ neurons (Fig. S1B). Thus, cell fusion can be excluded in our experimental paradigm.

**Fig. 1.**
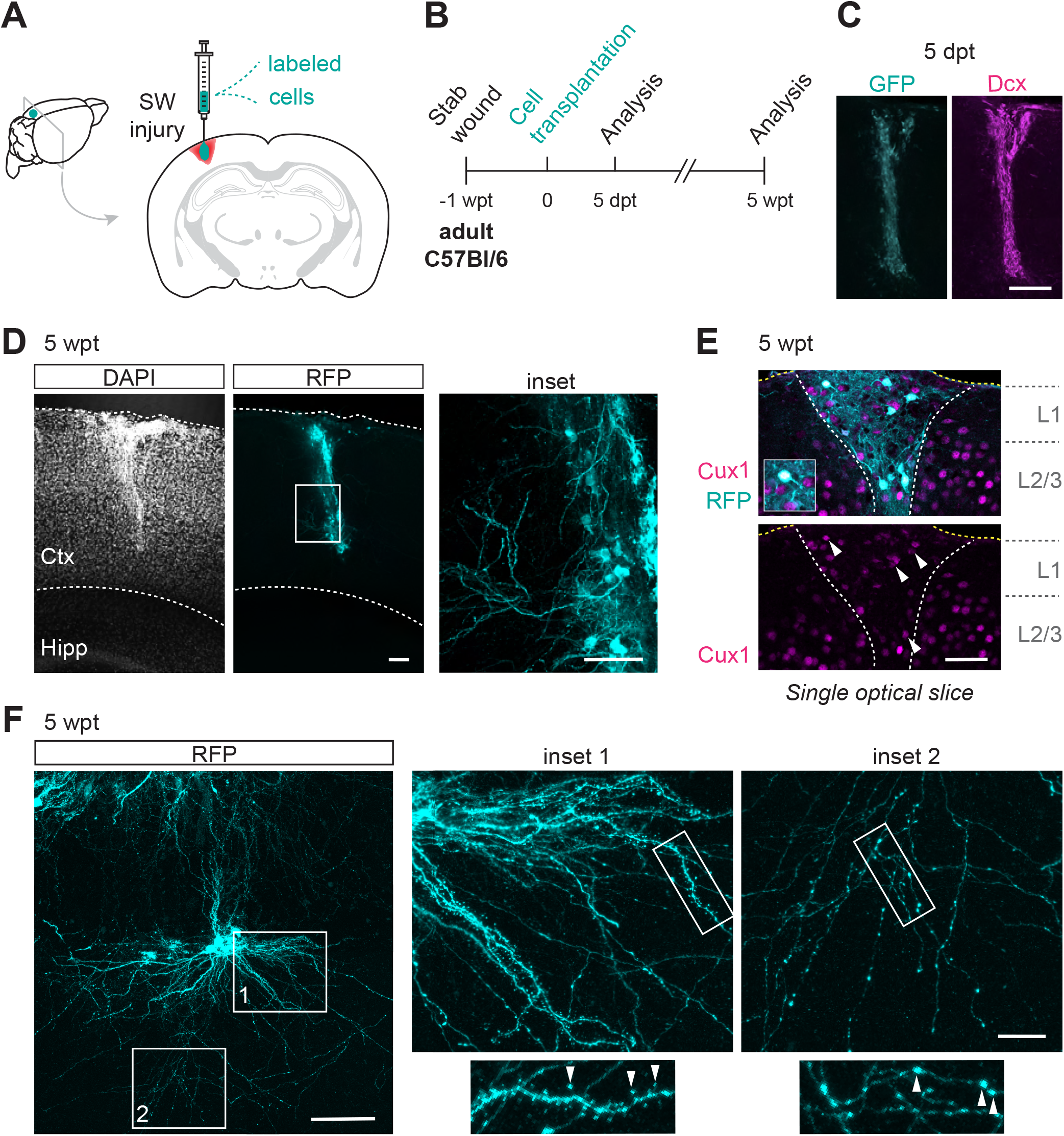
Transplanted neurons develop mature morphologies and synaptic structures within a cortical stab injury. (**A-B**) Schematic and timeline of the experimental procedure. (**C**) Representative confocal image of a GFP graft, 5 days post-transplantation, showing an almost complete overlap with the immature neuronal marker Dcx (n=4). (**D**) Confocal images of a representative transplantation site in the mouse cortex inflicted by a SW injury, 5 wpt (see all nuclei stained with DAPI; n=5). Boxed area highlights dendritic branches of RFP transplanted neurons. (**E**) Example image of a single optical section in the transplant shows colocalization of RFP with Cux1, a marker of upper layer cortical identity (n=2). (**F**) Z-stack projection of an example of grafted neurons and respective high magnification insets shows the extension of apical dendrites as well as profuse basal dendrites (1) and axonal arborizations (2) from the cell bodies (n=5). Notice the appearance of spines and boutons in dendrites and axons, respectively (arrowheads in the highlighted neurite). Scale bars: (C) 100 μm, (D) left, 100 μm, right, 50 μm (E) 50 μm, (F) left, 50 μm, right, 10 μm. Ctx, cortex; Hipp, hippocampus.

Together, these observations demonstrate that cells from the embryonic cortex transplanted into adult mouse cortex subjected to an invasive injury survive and develop morphological traits of mature cortical neurons, such as complex dendritic arbors and synaptic specializations.

### Host environment dictates neuronal integration: injury promotes initial integration of neuronal transplants

To uncover the synaptic network established between transplanted neurons and the host brain we used a modified RABV that allows monosynaptic tracing of inputs to targeted cells (33). In this deletion mutant rabies virus, ΔG RABV, the glycoprotein gene (G), necessary for the transsynaptic spread, is deleted and replaced by a fluorescent reporter. Additionally, the virus envelope is EnvA-pseudotyped to infect only cells expressing TVA. As a result, the primary infection is targeted to cells co-expressing TVA receptors and the rabies G allowing transcomplementation and assembly of new infectious particles. We therefore engineered donor cells derived from embryonic day (E) 14 cerebral cortex cultured *in vitro* by retroviral infection to express RFP/G/TVA prior to transplantation, or alternatively, transplanted acutely dissociated cells from E18 Emx1-Cre/G-TVA/GFP embryos (30,31,34). Complementary GFP or mCherry-expressing ΔG RABV, injected 1 month after transplantation thus propagates retrogradely and across one synapse to the direct pre-synaptic partners. As they lack G protein, infection is halted at these cells allowing unambiguous identification and mapping at fine scale the pre-synaptic partners of transplanted neurons brain-wide (Fig. 2A). Most importantly, by combining the comprehensive anatomical registration of single cells across the entire brain with computation of the connectivity index (number of neurons in a given anatomical region per primarily infected neuron also called “starter” neuron e.g. Fig. 2B) we can run comparative analysis between connecting areas or experimental groups (5). Note that every batch of RABV was tested in naïve C57BL/6J wild-type mice beforehand and rare batches resulting in transduction of few cells in these experiments were not used in this study. Besides, we have ran *in vitro* controls that exclude changes in G protein levels in the transduced neurons by inflammatory stimuli, such as interferon gamma, lipopolysaccharide (LPS) and interleukin 1b *in vitro* (Thomas et al., submitted back-to-back). This is an important control since variable levels of G across conditions could result in different ability of the virus to spread and thus quantitative differences in connectivity.

**Fig. 2.**
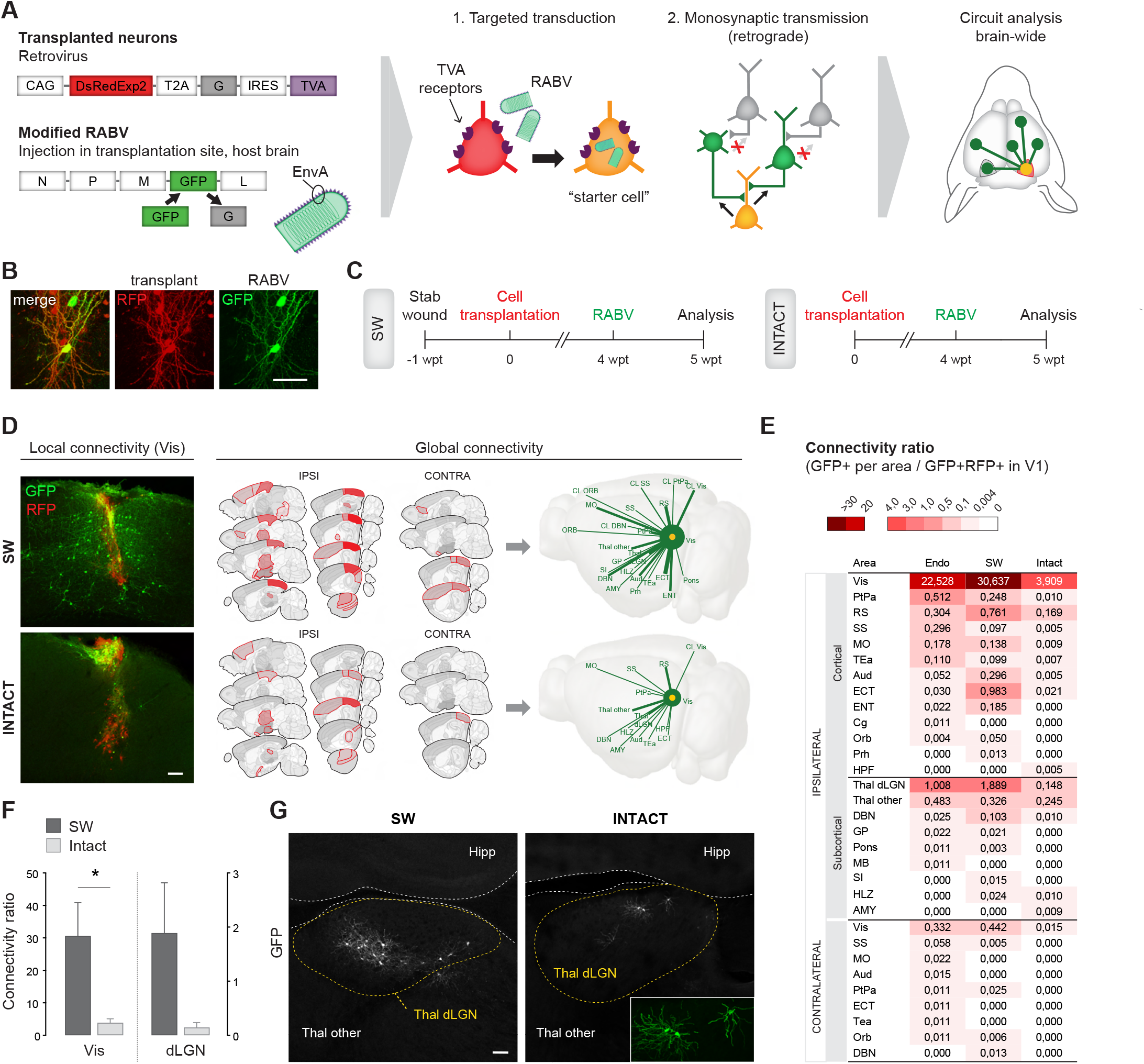
Input connectivity of neuronal transplants in SW and intact cortex. (**A**) Molecular tools and rationale of brain-wide monosynaptic tracing. (**B**) Example of a ‘starter’ neuron (RFP+/GFP+): a neuron within the RFP transplant (RFP+) that has been infected by the GFP rabies virus. (**C**) Experimental timeline. (**D-E**) Local and brain-wide inputs (GFP-only) to transplants, traced at 4 wpt (n=4/5). (**D**) Schematics depict brain regions that innervate the transplant. Red grading in sagittal sections and thickness of the lines in 3D connectograms reflect the connectivity ratio for a given connection. (**E**) Color-coded connectivity ratio for transplants in SW or intact cortex (n=4/5 respectively), as well as for endogenous neurons. The data shown for endogenous neurons (Endo) have been published before (5) and are used here solely for comparison. Note the excessive connectivity in SW and scarce in intact, as compared to the native. (**F**) Quantification of Vis-Vis and dLGN-Vis connectivity (n=4/5, *p < 0.05 using Mann-Whitney test). (**G**) Pre-synaptic neurons (GFP+) in the dLGN of the thalamus. Scale bars: (B) 50 μm, (D,G) 100 μm. See Table S1 for abbreviations. Contra, contralateral; Ipsi, ipsilateral.

To investigate whether neurons can connect properly within a gliotic microenvironment and the influence of a preceding injury, we transplanted reporter/G/TVA-expressing cells into either the SW-inflicted or intact primary visual cortex of adult mice, injected ΔG RABV (expressing a different reporter protein) 1 month later in the transplantation site and examined the brains 1 week afterwards (Fig. 2A, C). Mapping and quantification of pre-synaptic input neurons showed the highest connectivity index for the visual cortex (primary and high-order areas) among all innervating areas in both experimental conditions (Fig. 2D; see Data S1 for details). However, these short-distance connections within the visual cortex were abundant for transplants in the SW, while they were extremely scarce for neurons in the intact cortex with ∼8x lower connectivity index (Fig. 2D-F). Also, the global input landscape across the brain was strikingly different between these conditions, with more synaptic connections per region for neurons in the SW-injured cortex and with overall more connecting regions, compared to neurons in the intact cortex (23 versus 15 afferent regions for transplants in SW and intact cortex, respectively) (Fig. 2D, E, Fig. S2; see Table S1 for abbreviations). In both groups, neurons receive input from various cortical and subcortical regions of the ipsilateral hemisphere including sensory cortices, associative areas and thalamic nuclei, but less regions are represented in the connectome of transplants in the intact cortex. A major source of inputs to the mouse primary visual cortex is the dorsal lateral geniculate nucleus (dLGN), in the thalamus, which relays visual information from the retina to the primary visual cortex (35). While transplants in the SW visual cortex receive a considerable input from the dLGN, those in the intact group hardly do so (Fig. 2F, G). Likewise, in the latter, input from the contralateral hemisphere is reduced to a minor innervation from the visual cortex, whereas a few other cortical and subcortical regions of the contralateral hemisphere contribute to graft connectivity in the SW injury condition (Fig. 2E). Thus, the injury condition has a profound impact on the input connectome of neuronal transplants.

To determine how close these connectomes were to the connectome of endogenous neurons, we used our previously published data obtained by electroporation of upper layer visual cortex neurons with the RFP/G/TVA plasmid during mouse cortical development and subsequent RABV-mediated tracing during adulthood (5). Notably, the RABV strain and helper constructs, experimenter and data analysis pipeline were the same in the present study and previous study. The overall number of afferent regions was found to be very similar with 25 regions for endogenous neurons and 23 regions for regenerated circuits after SW. However, the quantitative comparison revealed significantly higher connectivity ratios for neurons transplanted in the SW cortex (Fig. 2E). Here, transplants receive input from more neurons in their surrounding visual areas. We could also appreciate a seemingly stronger innervation from the neighboring retrosplenial and ectorhinal cortices, and from the distant thalamic dLGN, a key component of the visual pathway as aforementioned. In stark contrast, the connectivity of transplanted neurons in the intact brain lags significantly behind the endogenous rates, with a local connectivity ∼6x lower, and lower connectivity index for most of the afferents throughout the brain (Fig. 2E). Importantly, for both groups all the identified regions are known to project to the visual cortex (36,37; Allen Connectivity dataset).

Thus, neuronal integration into pre-existing circuits of the mouse cerebral cortex requires an altered local environment, but a highly inflammatory and gliotic parenchyma is overly permissive to synaptogenesis resulting in supernumerary graft-host connections.

### Neuronal survival and differentiation in the intact cortex resemble those in injured cortex

One possibility explaining the poor input connectivity to neurons derived from transplants into the intact cortex, could be poor differentiation and/or survival. To explore this, we analyzed grafts at early time points after transplantation (Fig. S3A). Five days after transplantation into the intact cortex, the graft was largely immunopositive for Dcx similarly to that observed in the SW grafts (Fig. S3B and 1C). This finding suggests that the transplanted cell population goes through a similar trajectory with no major imbalance or lag in their differentiation. Moreover, neurons had already projected many neurites through the host parenchyma (Fig. S3B, C). These neurites display enlarged terminal structures at their tips which were reminiscent of growth cones, suggesting ongoing pathfinding. A few had reached the corpus callosum at about 300-400 μm from the transplant, indicating no major obstacle to their navigation through the cortical tissue. By 2 wpt, spine-bearing dendrites were identifiable (Fig. S4C) and neurons had acquired expression of the mature neuronal marker NeuN (Fig. S4B). Many co-expressed the upper layer neuron marker Cux1 as observed for transplants in a SW injury (Fig. S4B and 1D; 63.47% and 70.27% respectively) or in a neuronal ablation injury (5). Finally, graft volume was considered as a proxy for graft survival and no significant difference on graft volume was detectable between intact or SW cortex despite a trend of the latter towards larger volumes (Fig. S5).

Altogether, these data indicate that also the environment of the intact cerebral cortex can nurture early neuronal development, but fails to allow synaptic integration of new neurons.

### LPS-elicited inflammation is insufficient to promote graft-host connectivity

One obvious difference between intact or SW-injured cortex is the activation of glial cells and the inflammatory stimuli elicited by this invasive injury (15,16,24,26,38). To determine if such an inflammatory component of an injury may be sufficient to elicit the circuit plasticity required for the adequate integration of new neurons, we injected bacterial LPS intraperitoneally (i.p.), transplanted cells 1 week later and then analyzed their connectivity. First, we monitored the reactive gliosis elicited by different LPS concentrations at 1 week post-injection in comparison to the reaction elicited by the SW injury (Fig. 3A). Chosen concentrations were previously used as an inflammatory challenge (39–41) without neuronal cell death (42,43). Reactive astrocytes and microglia were stained by GFAP and Iba1 respectively and showed evident reactivity in the SW and with the higher LPS dose even though not the very same extent as in the penetrating stab injury (Fig. 3A,B; note that GFAP in LPS is not significantly higher than in the intact group in spite of a trend). Hypertrophic astrocytes and de-ramified microglia were noticeable in these two groups as compared to the resting-like counterparts characterized by lack of GFAP and thin/multiple processes respectively, observed in the naïve cortex and contralateral to SW. In contrast, immunoreactivity and glial cell morphology with the lower LPS dose was comparable to the contralateral visual cortex from SW brains or naïve mice (Fig. 3A).

**Fig. 3.**
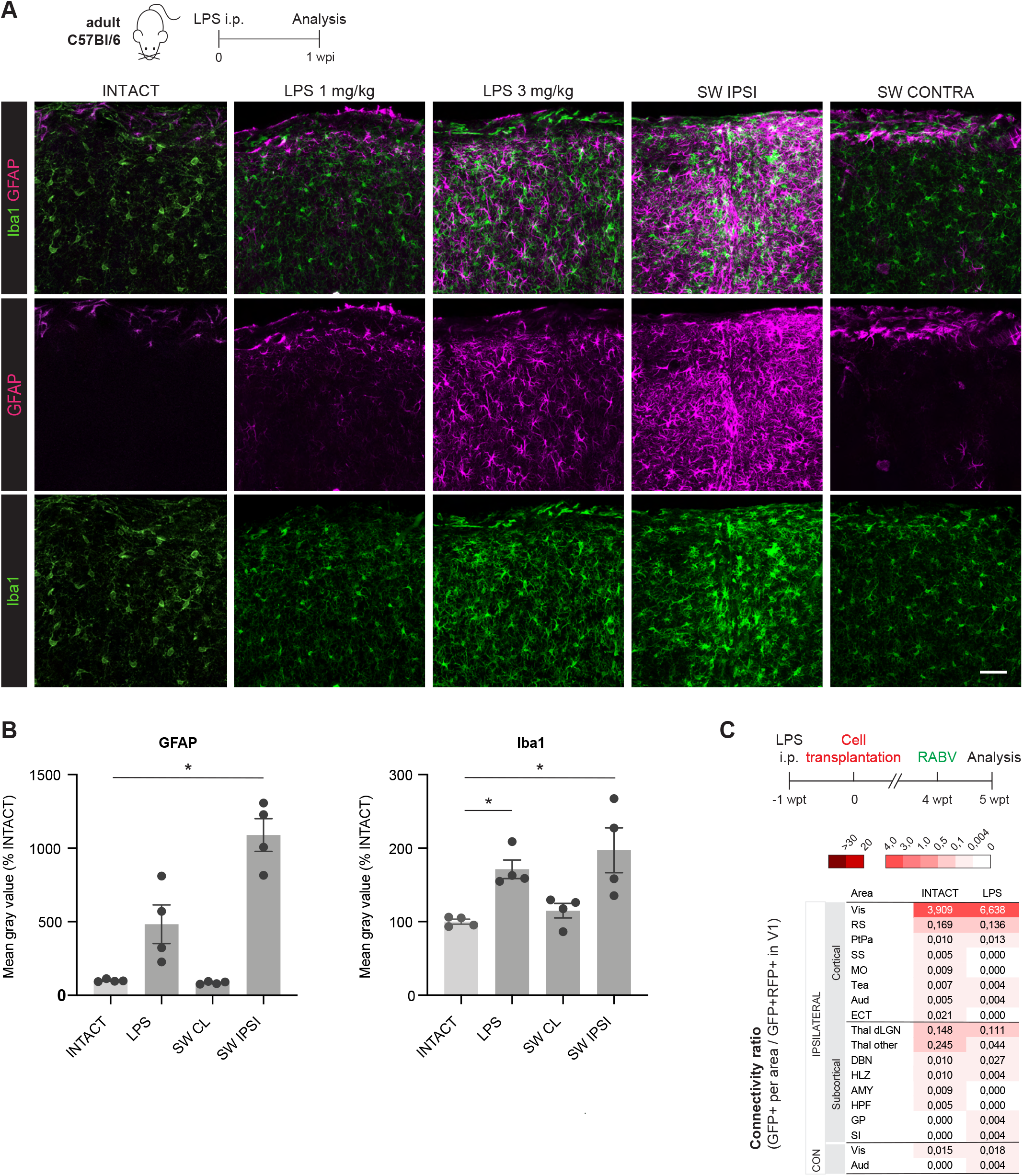
Gliosis in the various experimental groups and input connectivity of neuronal transplants in LPS-induced cortex. (**A**) (**C**) Timeline (top) and confocal images of the visual cortex of mice treated with 1 or 3 mg/kg LPS, or inflicted with a SW injury, and controls (intact (naïve mice) and contralateral SW, as an additional control to SW). Sections were immunostained for microglia and reactive astrocyte markers (Iba1 and GFAP respectively; bottom) (n=4 for all conditions except LPS 1 mg/kg where a n=2 mice was analyzed). Note the cellular response with the highest concentration of LPS is much closer to that observed in the SW-inflicted cortex. (**B**) Mean gray value of all pixels, calculated using Z-stack projections of similar thickness, for intact, LPS 3 mg/kg and SW groups (n=4; note that cortical layer 1 was excluded from the selected area). (**C**) Timeline and analysis of the brain-wide monosynaptic input to grafts in the brain of 3mg/kg LPS treated mice as compared to those in the brain of intact mice (n=5/6 respectively). Scale bar: (A) 50 μm. See Table S1 for abbreviations. Con/Contra, contralateral; Ipsi, ipsilateral.

We therefore used the higher LPS dose eliciting stronger reactive gliosis to test the influence of inflammation in graft connectivity (Fig. 3C). Connectivity of neuronal transplants in the cortex with LPS-induced reactive gliosis was rather similar to those in the intact cortex, despite a small trend for a higher local connectivity (15 versus 13 afferent regions; 6.64 versus 3.91 connectivity ratio for visual-visual connections, for LPS and intact, respectively). These experiments revealed a minor or neglectable role for inflammation and the ensuing reactive gliosis in graft-host synaptic connectivity.

### Increased levels of complement proteins and reduced levels of synapse proteins in the SW cortex

As LPS-induced inflammation showed little effect boosting the integration of the neurons differentiating in the transplants, we pursued an unbiased proteomics approach to identify mechanisms fostering integration of new neurons in the host environment. Using a biopsy punch, we collected the host visual cortex at the time when cells would normally be transplanted, i.e. 1 week after inflicting the SW injury or after LPS injection, and collected the same region from age- and sex-matched control visual cortex (“intact”; Fig. 4A). We collected the ipsilateral visual cortices of SW mice (n = 10 hemispheres, from 10 mice), both visual cortices of LPS-injected mice (n = 10 hemispheres, from 5 mice) and both visual cortices of control intact mice (n = 20 hemispheres, from 10 mice). By shotgun proteomics using liquid chromatography-coupled tandem mass spectrometry (LC-MS/MS) we reproducibly detected a total of 5225 proteins and quantitatively compared the samples testing for significant differences (t-test). First, we compared each of the conditions (SW/LPS) with the control samples (intact brain).

**Fig. 4.**
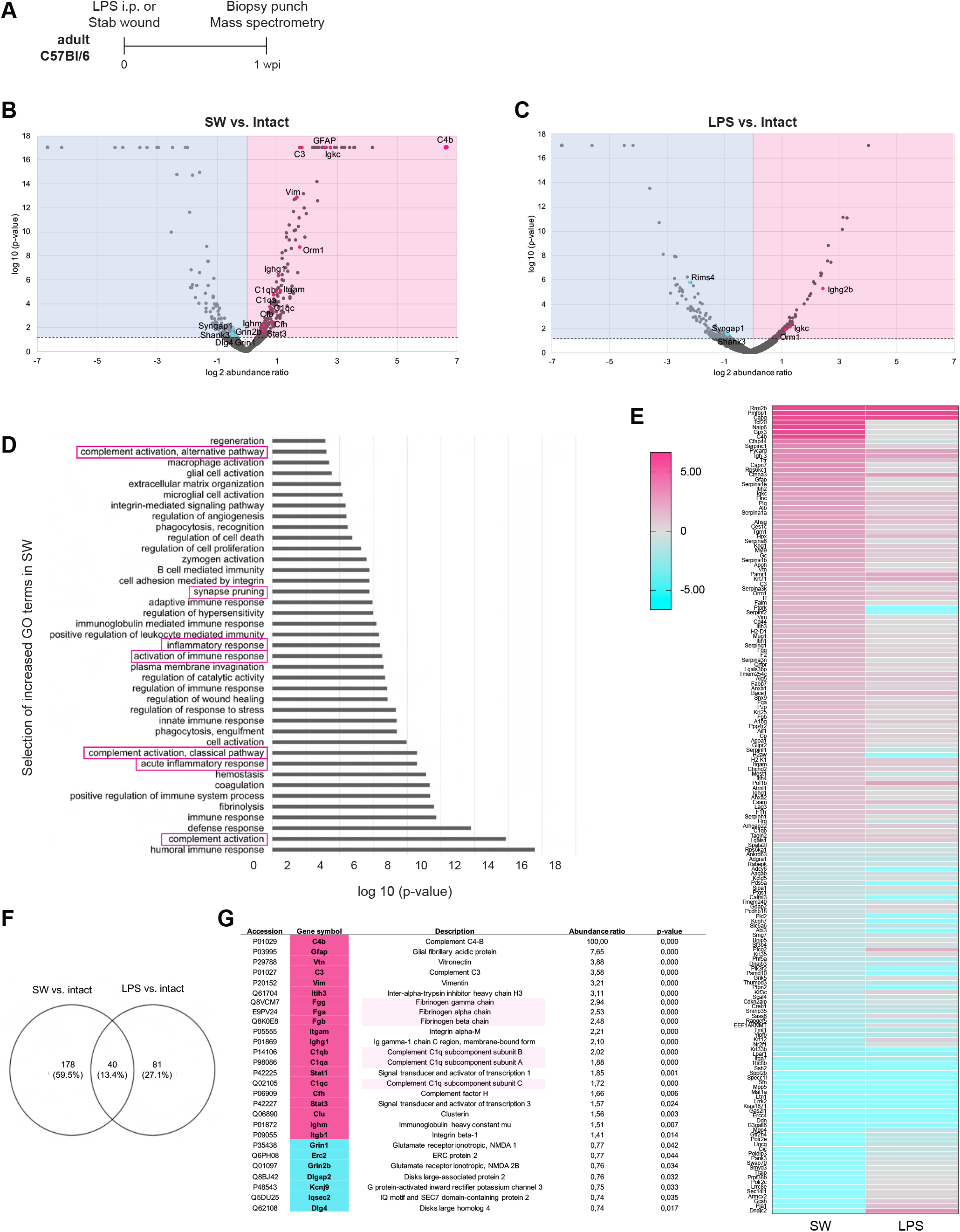
Comprehensive proteome analysis of SW-injured and LPS-inflamed visual cortex. (**A**) Timeline for tissue punch collection. (**B-C**) Volcano plots showing log2 mean abundance ratio and corresponding log10 p-value comparing SW-injured (n=10, B) and LPS-treated (n=10, C) with intact control (n=20) cortical tissue (derived from 10, 5 and 10 mice respectively, as we can consider both hemispheres from LPS and intact brains). Upregulated proteins in pink area and downregulated proteins in blue area of the plots. (**D**) Selection of enriched GO terms (biological process) of significantly enriched proteins in SW vs. intact cortex. For the complete set of GO terms see Data S2. (**E**) Heatmap shows the significantly regulated proteins in the SW cortex, along with their regulation in the LPS cortex. (**F**) Venn diagram depicts differentially regulated proteins that overlap or are exclusive for each condition. (**G**) Selection of proteins which are exclusively up- or downregulated in the SW-injured cortex. Highlighted are proteins further tested by immunostaining (see Fig. S5). See Data S2 for protein and GO analysis. wpi, weeks post-injury (SW)/injection (LPS).

After SW injury, 221 proteins differed significantly in their abundance compared to the control (Fig. 4B, Data S2a). A total of 168 proteins were significantly enriched in the injury condition including some expected proteins, such as GFAP (7.65 fold) and vimentin (3.21 fold; Vim) which are upregulated in reactive astrocytes (see Fig. 3A for GFAP) (15,38,44). Likewise, the typical injury associated extracellular matrix (ECM) proteins transglutaminase 1 and 2 and inter trypsin alpha inhibitors (16) were significantly enriched. Consistent with an ongoing inflammatory response, gene ontology (GO) term enrichment analysis for biological processes showed significantly enriched categories such as “acute inflammatory response”/”inflammatory response” and “activation of immune response” including e.g. the complement factor C3, complement C1q subcomponent subunits A, B and C (C1qa, C1qb, C1qc), immunoglobulin heavy constant gamma 1 (Ighg1), immunoglobulin kappa constant (Igkc), alpha-1-acid glycoprotein 1 (Orm1), immunoglobulin heavy constant (Ighm), signal transducer and activator of transcription 3 (Stat3) and Ig gamma-2B chain C region (Ighg2b) (highlighted GO terms in Fig. 4D, Data S2b). Indeed, we noted several complement system-related proteins, besides C3 and the various C1q, like complement C4 B (C4b), complement factor b (Cfb) and complement factor h (Cfh). Altogether, these are involved in the classical and alternative pathways of the complement system that propagate neuroinflammation (45–47) and have been implicated in synapse and dendrite remodeling in different conditions (46,48,49). Accordingly, GO term enrichment analysis revealed the significantly enriched categories “complement activation” and “synapse pruning” that both include C4b, C3, C1qa, C1qb, C1qc, Cfh, Cfb, Igkc, Ighg1, integrin alpha-M (Itgam), amongst other proteins. Consistent with the damage to the blood brain barrier (BBB) in this injury model (50), each of the three polypeptide chains forming the blood protein fibrinogen (fibrinogen alpha, beta and gamma chains, i.e. Fga, Fgb and Fgg) was significantly more abundant. Interestingly, fibrinogen induces spine elimination via microglia activation in a model of Alzheimer’s disease with vascular dysfunction (51). Fibrinogen binds the same receptor as complement protein (CR3 or CD11b) (52,53) and presumably both ligands thus contribute to microglia-mediated synapse pruning in conditions involving a breach of the cerebral vasculature. The significant increase of both fibrinogen and C1q in the SW cortex was further verified by immunostaining (Fig. S6A,B).

We also detected 53 proteins with significantly decreased abundance after SW. These included proteins known to perform a variety of central roles in maintaining cell function. Amongst the most downregulated proteins are 14-3-3 sigma (Sfn), a protein that is highly conserved and multifunctional, regulating essential cellular processes from cell metabolism to transcription and signal transduction (54) and known to be downregulated in brain disease (55). Also proteins with central roles in transcriptional regulation and part of transcription machinery, such as the histone-lysine N-methyltransferase (Smyd3) and the DNA-directed RNA polymerase II subunit (Polr2c or Rpb3) were found to be significantly decreased (56,57). Interestingly, COUP transcription factor 1 (CoupTF1 or Nr2f1), a transcription regulator that controls neurogenesis/gliogenesis balance, was found to be down-regulated in our data, and in previous work upon neuroinflammation (58) (Data S2a). Other proteins involved in neurogenesis are downregulated such as the chromatin remodeling factors: transcription activator BRG1 (Smarca4), chromodomain-helicase-DNA-binding protein 5 (Chd5) and transcriptional repressor p66-beta Gatad2b. Accordingly, GO terms include “histone modification”, “regulation of neurotransmitter receptor activity”, “regulation of neuronal synaptic plasticity” (Data S2b). The latter includes disk large homolog 4 (Dlg4), SH3 and multiple ankyrin repeat domains protein 3 (Shank3), glutamate receptor ionotropic NMDA 1 and NMDA 2b (Grin1 and Grin2b), Ras/Rap GTPase activating protein (Syngap1) suggesting a decrease of synaptic structures after brain injury (Fig. 4B, Data S2a,b). These data are consistent with reduced neuronal excitability and firing in the acute phase after TBI (59).

Comparing the LPS-induced cortex biopsies to controls, we found 123 proteins of significantly different abundance (Fig. 4C, Data S1c). Amongst the 60 proteins more abundant upon LPS injection, we observed Ighg2b, Igkc and Orm1, similar to the increased proteins in the brain injury. We also found the increased GO term “acute inflammatory response” to be enriched in LPS condition (Data S2d). Amongst the 63 proteins with lower abundance, e.g. Sfn is similarly on the 3 most downregulated, and also synapse related proteins like Shank3, Syngap1 and regulating synaptic membrane exocytosis protein 4 (Rims4), suggesting some, albeit small (Fig. 4E), overlap and similarity between SW and LPS conditions.

We next focused on the proteins uniquely regulated upon SW and not upon LPS as they may contribute to fostering the integration of new neurons (Fig. 4E-G, Data S2e). Indeed, the activation of the complement system, synapse pruning, and immune system proteins like C4b, C3, C1qa, b, c, Itgam, Ighg1 were specific in this environment, as was the enrichment of GFAP, Vim, Fga, Fgb and Fgg. Also, the downregulated proteins Dlg4, Grin1, Grin2b, and others that are involved in synaptic plasticity were injury-specific.

Taken together, proteome comparison of microenvironments eliciting low and high graft connectivity reveals fibrinogen and complement activation as possible mechanisms that may promote integration of new neurons into synaptic networks of the host brain. At the same time, these findings led us to ask if their integration is stable or synapse pruning lingers for a longer period and eventually affects also newly formed host-graft connections.

### Excessive graft-host connectivity in cortical stab injury is transient

Given the above observation that fibrinogen and complement factors are particularly abundant in the SW condition, and their documented role in synaptic pruning, we set out to determine the persistence of new synapses after the initial synaptic integration in this condition. In the neuronal ablation model we had observed a net increase in spine number in the first month, followed by 1 more month of active spine turnover with no overall change in input connections and a stable connectivity ratio between 1 and 3 mpt (5). We therefore traced input connectivity at 3 mpt in SW cortex and compared to the data obtained at 1 mpt (Fig. 5A). Surprisingly, by 3 months, connectivity with the host brain was severely diminished, with about 3 fold fewer local connections (Fig. 5B, C) and a total of 13 afferent areas as opposed to the earlier 23 (Fig. 4C, D). Importantly, connectivity had dropped to metrics below those of normal circuits (endogenous connectivity) with only 5 as compared to 12 afferents with connectivity ratio ≥ 0.05 respectively. Most of the weaker connections (connectivity ratio ≤ 0.03) observed at 1 mpt had been lost, with e.g. only one contralateral innervation persisting, from the contralateral visual cortex. Even though the calculated connectivity ratio accounts for the number of starter neurons, it is important to note that grafts at later time points were overall smaller (Fig. S5B) suggesting that not only loss of synapses but also cell death has occurred between 1 and 3 mpt. This is a notable difference from other conditions, such as in the brains with targeted neuronal ablation (Falkner et al., 2016). These results imply that early formed synapses with transplants in SW are eliminated in the course of the following months and thus call for caution and the need to explore specific injury conditions prior to use in neuronal replacement therapy.

**Fig. 5.**
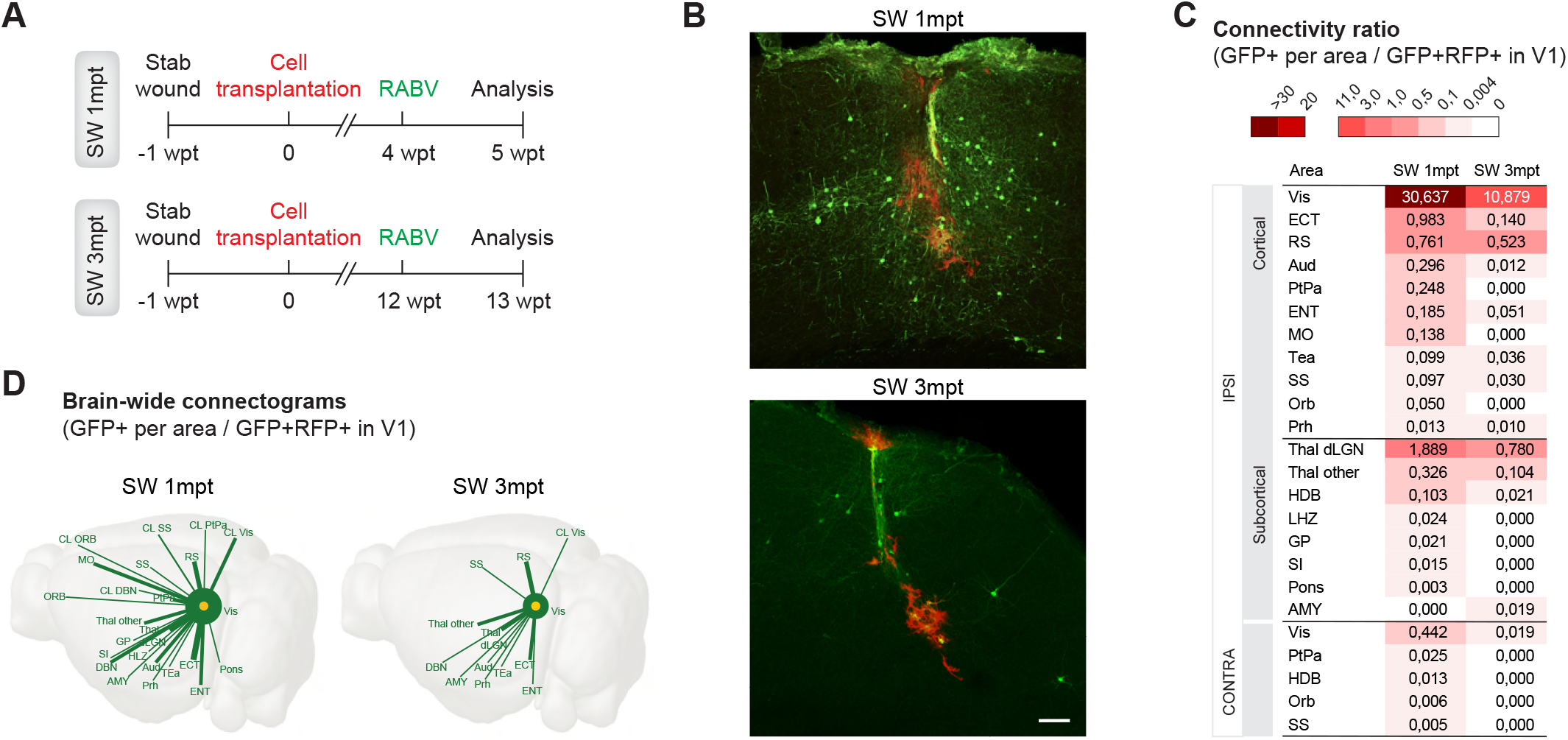
Comparison of early and late connectivity shows transience in cortical stab injuries. (**A**) Experimental timelines. (**B**) Local inputs (GFP-only) to transplants, traced at 4 and 12 wpt. (**C**) Color-coded brain-wide connectivity at 1 or 3 mpt in SW (n=4/6 respectively). Decreased connectivity at 3 mpt shows that many of the early synaptic connections have been pruned. (**D**) Distribution and strength of single host-graft connections evidenced by thickness of the lines between the graft (yellow) and each area (green). Scale bar: (B) 100 μm. See Table S1 for abbreviations.

## Discussion

Despite the broad experimental use of neuronal transplantation for cell replacement in cortical brain injury, an outstanding open question is still whether the injured environment, variable across pathologies, plays a role in the integration of transplanted neurons into pre-existing circuits. Our data show that the input connectome of transplanted neurons is highly influenced by the local environment where they develop. While the transplant survival and gross morphological maturation share features across different hosting environments, its degree of input connectivity with the host circuits and stability of these new connections differs substantially.

By transplanting the same donor cell type as previously in a neuronal ablation model (5) now in a penetrating TBI we demonstrated pronounced differences in initial and long-term input connectivity, with initial excessive inputs from the host networks that were eventually pruned to sub-normal levels. In contrast, neurons grafted in intact or LPS-induced networks were scarcely innervated by the host brain.

Neurons derived from mouse embryonic stem cells fail to project long-distance axons after transplantation if no injury is previously inflicted, as opposed to an injured brain, despite good graft survival in both experimental groups (60). Transplantation into the intact dentate gyrus or striatum of adult immunodeficient Rag2-/- mice yielded widespread inputs to large transplants, but their quantitative nature is unknown (61). Interestingly, cells from the spinal cord of rat embryos transplanted into the adult rat spinal cord fail to contact nearby corticospinal axons if no spinal cord injury precedes (62). Also, transplantations of embryonic neurons in the postnatal mouse cortex have suggested that a preceding apoptotic injury promotes morphological maturation and synapse formation between the host and graft (63). Clearly, there are obstacles to the development of graft outputs and inputs in the intact central nervous system. Our findings add knowledge by showing a priming effect of an injury opening brain networks to accommodate new neurons and form new synapses in the adult cerebral cortex. On the flipside, it became now obvious that not all injuries lead to the same level of host-graft connectivity in the long-term.

Several previous studies have shown that neuronal grafts in the injured postnatal or adult rodent cortex receive inputs from the host brain (5,6,13,60,63). However, a comprehensive comparison of the connectivity levels with native circuits has been a more challenging endeavor addressed by only a few studies (5,6). Based on our findings herein using the SW injury, we could hypothesize that an excessive input is also observed in transplants developing in another inflammatory condition like stroke. However, using a semi-quantitative analysis, Tornero et al. (6) have reported input connectivity that resembles the normal circuitry when quantifying the contribution from each input region in 3 categories. This quantification may not resolve all the differences we found here. In addition, possible differences may be attributable to several reasons. First and foremost, transplantation of human induced pluripotent stem cells (iPSC)-derived neurons requires immune suppression and hence alters the environment. Second, the time and site of transplantation differs, since the authors grafted iPSC-derived cortically fated neurons nearby the stroke area and only 48h after the stroke (compared to 1 week post-injury in our study) when microglia numbers and reactivity at the site of transplantation is peaking and thus synapse pruning may be exaggerated leaving the initial connectome at lower metrics. Third, the degree of maturation differs profoundly with a much slower maturation speed of human neurons that are still rather immature at the time of RABV injection (2 mpt; only 20% NeuN) (64) and thus the mapped connectivity may be a premature assessment of an expanding connectome. Importantly, data from brain imaging and network neuroscience has been converging into the view that TBI is characterized by an increased endogenous connectivity degree, while the transience of this hyperconnectivity is still under debate (65). Slicing the brain also correlates with a substantial, rapid increase in synapse number compared to tissue fixed *in vivo* (66). Most importantly, however, stroke differs in many parameters from TBI, and may hence indeed influence connectivity differently, highlighting the need to examine quantitative connectivity specifically in different injury conditions. This is also confirmed by our accompanying manuscript (Thomas et al.) showing the influence of the aging brain and amyloid-loaded host environment in promoting the input connectome locally.

Our data showed a sharp decline in the number of host-graft connections in the SW cortex at 3 mpt resulting in an extremely scarce connectome, likely insufficient for functional repair. These findings are difficult to compare with transplants of human PSC-derived neurons as these mature at a very different pace and develop in an immunesuppressed environment. Interestingly, however, previous work showed a surprisingly early innervation already at 1.5-2 mpt after transplantation that is comparable at more mature stages at 6 mpt, in stroke and Parkinson’s disease (PD) models (6,7). It will be intriguing to analyze these transplants at even later stages, given the slow pace of human neuron maturation (67,68).

Analysis of the proteome from environments that promote graft connectivity (SW) as compared with those where transplant connectivity remains under normal levels (intact/LPS) suggests that fibrinogen and complement activation pathways play a role in synapse yield. Both have been implicated in synapse pruning during brain development and/or neurological disease namely in TBI (48,51,69,70). Interestingly, gene expression analysis in the neuronal apoptosis model where we have previously observed an adequate and stable level of connectivity, showed all the three components of C1q (C1qa, C1qb and C1qc) differentially overexpressed in injured regions (8). Collectively, our findings indicate that complement protein levels and its temporal dynamics at a brain injury may play an essential role dictating the outcome of transplantation and stability of new synaptic connections.

Successful brain repair by cell transplantation does not only involve axonal outgrowth and generation of new synapses with the correct regions, but also a tight quantitative match of the new input and output connections. In the mammalian cortex, excessive excitatory connections may result in hyperexcitability and formation of epileptic foci in the transplanted site, while scarce connections may be insufficient to restore the functional deficit caused by the injury. Herein, we provide the first evidence that injury type-dependent extrinsic cues influence the integration of transplanted neurons into the host synaptic circuitry, in specific, their afferent connectivity. Future research is needed to investigate the functional relevance of the inputs mapped herein, or in another model of acute brain injury, such as stroke. This work further advances our understanding of the prerequisites for neuronal replacement in damaged brain circuits and represents an important step in clinical translation of neuronal replacement strategies to restoring an authentic circuitry after brain damage and promote functional recovery in the injured brain.

## Materials and Methods

### EXPERIMENTAL DESIGN

#### Animals

Animal handling and experimental procedures were performed in accordance with German and European Union guidelines and were approved by the Government of Upper Bavaria. All efforts were made to minimize suffering and number of animals. Both male and female mice were used. Mice were kept in specific pathogen-free conditions and in 12:12 hour light/dark cycles with food and water *ad libitum*.

All mice were 2-4 months old at the time of the first surgery. C57BL/6J wildtype mice were used as host mice for all studies except those to test for the occurrence of cell fusion wherein tdTomato reporter mice were used (Ai9) (32). Surgeries were performed aseptically under anesthesia with a mixture of fentanyl (0.05 mg/kg, Janssen), midazolam (5 mg/kg, Roche) and medetomidine (0.5 mg/kg, Fort Dodge). After surgery, anesthesia was terminated with atipamezol (2.5 mg/kg, Janssen), flumazenil (0.5 mg/kg, Hexal) and buprenorphin (0.1 mg/kg, Essex). Meloxicam (1 mg/kg, Metacam) was administered for postoperative analgesia.

Cells for transplantation were obtained from C57BL/6J wildtype or Emx1-Cre/EGFP E14.5/E15.5 mouse embryos (30,31) and cultured (see below), or alternatively, they were mechanically dissociated from E18.5 Emx1-Cre/G-TVA/GFP embryos (30,31,34) and readily transplanted. Triple transgenic embryos amid the litter were identified by GFP expression in the dorsal telencephalon and resulted from matings involving a G-TVA homozygous parent.

#### Cell cultures

##### Primary culture of cortical neurons

Neocortex from E14.5/E15.5 mouse embryos was mechanically dissociated in Hanks’ balanced salt solution (HBSS) buffered with 10 mM HEPES (both from Life Technologies). Cells were plated in 20 μg/ml poly-D-lysine (PDL, Sigma-Aldrich) coated 24-well plates, at a density of 200 000 cells/well. Cells were initially kept in 10% fetal bovine serum (FBS, Pan Biotech)-containing DMEM high glucose (4.5 g/L) with glutamax, plus penicillin-streptomycin (both from Life Technologies). Serum was gradually removed by replacing half of the medium with B27 (Life Technologies)-containing DMEM high glucose (4.5 g/L) with glutamax plus penicillin-streptomycin on each of the following two days. Cells were harvested for transplantation after 4-5 days *in vitro* upon validation of reporter expression.

All cells for primary culture were obtained from wildtype C57BL/6J mice except those used to test for the occurrence of cell fusion with the host cells. For this purpose, Emx1-Cre/EGFP (30,31) embryos were used. Double transgenic embryos amid the litter were identified by GFP expression in the dorsal telencephalon.

#### Virus treatment in cell cultures

Neocortical neurons from C57BL/6J wildtype mouse embryos were cultured. Two to four hours after plating the cells were transduced with MMuLV-derived retroviral vectors CAG-EGFP or CAG-DsRedExpress2-2A-Glyco-IRES2-TVA (0.8-1 μl/well; titers ranged from 10^7^ to 10^11^ transducing units per mL). Cells were then transplanted into C57BL/6J wildtype adult mice. For cell fusion test experiments, neuronal cultures were prepared from Emx1-Cre/EGFP E14.5/E15.5 embryos and transplanted into tdTomato reporter mice. Although these cells are endogenously labeled we additionally treated them with CAG-EGFP retrovirus to account for a putative impact of retroviral transduction in the propensity of cells to fuse upon transplantation in the adult brain parenchyma.

#### Treatments and Surgical procedures

##### LPS inflammatory stimulus

C57BL/6J wildtype mice were subjected to an inflammatory stimulus by a single i.p. injection of 1 mg/kg or 3 mg/kg lipopolysaccharide (LPS) from E. coli (Sigma). A batch of mice was used to run a comparative analysis of cortical gliosis in SW, LPS and control conditions. For this purpose, a craniotomy was performed at the time of the LPS injection to account for glial reactivity due to the craniotomy *per se*. These mice were perfused 7 days after the LPS treatment, in order to monitor cortical gliosis at the time when transplants would normally be performed. Other mice were used for subsequent neuronal transplantation and connectivity analysis (see below).

##### Stab wound injury (SW)

C57BL/6J wildtype or tdTomato reporter mice were used for cortical SW injury as described in Mattugini *et al*. (26). Briefly, mice were anesthetized and a craniotomy of 2.5 mm diameter was open to expose the primary visual cortex (V1) of the left cortical hemisphere. Using an ophthalmological lancet, a 0.5 mm-deep / 1 mm-long incision was performed within V1 borders (coordinates from lambda: 0.0±0.2 anteroposterior, 2.0±0.2 to 3.0±0.2 mediolateral; coordinates were chosen to avoid large pial vasculature). The bone flap was placed back and the skin sutured. For analysis of cortical gliosis, mice were perfused 7 days after SW, while for analysis of graft integration the mice were subjected to further surgical procedures (see below).

##### Cell transplantation

Cells were transplanted unilaterally into one V1 of the mouse cerebral cortex. Host mice belonged to three groups: SW, intact (naïve) and mice subjected to an inflammatory stimulus by i.p. injection of 3 mg/kg LPS. Transplantation into SW or LPS mice was performed 7 days after the respective insult. In SW group, donor cells were placed right into the center of the incision, still lightly visible.

Donor cells were fluorescently labelled in Emx1-Cre/EGFP or Emx1Cre/G-TVA/EGFP mouse lines or via *in vitro* viral transduction with the aforementioned constructs. Cultured cells were washed 3 times with pre-warmed phosphate-buffered saline (PBS) to remove any remaining viral particles and cell debris. Gentle trypsinization (0.025%, 10 min at 37 °C) detached the cells and trypsin was then inactivated by FBS-containing medium (1:1). The cell suspension was then prepared in B27-containing DMEM high glucose (4.5 g/L) with glutamax, plus penicillin-streptomycin. 25 000-50 000 donor cells were transplanted in a total volume of 1 μl of cell suspension using a ga33 Hamilton syringe (coordinates from lambda: 2.5 ±0.2 mm mediolateral, 0.0 ±0.2 mm anteroposterior). Cell deposits were distributed dorsoventrally filling up a depth between 0.5 to 0.2 mm that results in a cellular graft encompassing cortical layers 1 to 4/5. Injection coordinates and pattern of pial vasculature were noted for later identification of the transplantation site and injection of the RABV. The bone lid was repositioned and the skin was sutured. For analysis of transplanted neurons survival and early differentiation, some mice were perfused 5 days afterwards, while for analysis of neuronal integration mice were subjected to RABV injection as detailed next.

##### Rabies virus injection

We used retrograde monosynaptic tracing with a modified rabies virus (RABV: EnvA-pseudotyped ΔG-EGFP or ΔG-mCherry, complementary to the reporter in transplanted cells) (5,33) to map brain-wide synaptic input to grafted neurons. In short, the RABV was injected 1 or 3 months after cell transplantation, in three locations surrounding the transplantation site (200 nl/location), using an automated nanoinjector at slow delivery speed. RABV titers typically ranged between 1.5-3.5 × 10^8^ plaque forming units (pfu)/mL. Mice were sacrificed 7/8 days later for immunostainings and circuit analysis.

#### Immunostaining

Mice were deeply anaesthetized with ketamin (100 mg/kg) and xylazin (10 mg/kg) and perfused transcardially with PBS (5 min) followed by 4% PFA in PBS for 30-40 min. Brains were collected and post-fixed in 4% PFA overnight, at 4 °C, serially cut on a vibratome into 70 μm sagittal sections and slices were further processed as free-floating. Sections were washed and incubated in blocking and permeabilizing solution for 2 h (2-3% bovine serum albumin or 10% normal goat serum; 0.5% triton X-100). The following primary antibodies were then used: chicken anti-Green Fluorescent Protein (GFP, 1:1000; Aves Labs), rabbit anti-Red Fluorescence Protein (RFP, 1:1000; Rockland), goat mCherry (1:200; Sicgen), rabbit anti-Cux1 (1:200; Santa Cruz), guinea pig anti-Satb2 (1:250; Synaptic Systems), mouse anti-GFAP (1:1000; Sigma-Aldrich), rabbit anti-Iba1 (1:500; Wako), rabbit anti-Dcx (1:1000 Abcam), guinea pig anti-Dcx (1:1000, Merck), mouse anti-NeuN (1:200; Merck/Millipore), sheep anti-Fibrinogen (1:100, Biomol) and rabbit anti-C1q (1:1000, Abcam) for overnight to 48 h-incubation, at 4 °C). After washing, sections were incubated with appropriate species- and subclass-specific secondary antibodies conjugated to Cy3 or Cy5 (Dianova) or Alexa Fluor 488 or 647 (Invitrogen), used at 1:500 or 1:1000 depending on high (>1:500) or low (<1:500) concentration of the primary antibody. Sections were incubated for 10 min with 1 μg/ml 4,6-diamidino-2-phenylindole (DAPI; Sigma-Aldrich) for nuclear labeling and mounted on glass slides with Aqua-Poly/Mount (Polysciences).

For connectivity analysis, brain-wide, all sections were immunostained for GFP/RFP. For some brains, sections with the transplant were selected and subsequently stained for Cux1 colocalization analysis and all were mounted for microscopy and serial analysis.

#### Slice processing and imaging

For brain-wide connectivity analysis, brain sections were kept in serial order throughout their processing. Sections with one or more GFP (or mCherry)-labeled cell somas were scanned using an epifluorescence microscope with a motorized stage (Zeiss, Axio Imager M2) equipped with a 10x objective (NA 0.3). Automated scanning, tile alignment, and image stitching was performed to create a high-resolution image of the whole section. In sections with unclear cell numbers due to close apposition of two GFP (or mCherry) cell bodies or with high densities of GFP (or mCherry) cells, scanning of Z-stacks in a laser-scanning confocal microscope (Zeiss, LSM 710) with a 40x objective (NA 1.1) was carried out.

For all immunofluorescence studies, images were acquired using an epifluorescence microscope with a motorized stage (Zeiss, Axio ImagerM2) and a laser-scanning confocal microscope (Zeiss, LSM 710).

#### Mass spectrometry

3-4 months old male and female C57BL/6J mice (n = 10 intact controls, n = 10 SW-injured, n = 5 LPS injected) were sacrificed through cervical dislocation, brains were removed and placed into cold PBS. Biopsy punches (2.5 mm diameter) of the visual cortex of both hemispheres were dissected whereas meninges and white matter were carefully taken off. The contralateral (uninjured) cortices of the SW-injured brains were not considered for proteome analysis (intact cortical tissue: n = 20, SW: n = 10, LPS: n = 10). Samples were placed into low-protein binding Eppendorf tubes, frozen on dry ice, and stored at – 80 °C until further processing.

Tissue samples were lysed in NP40 buffer (1% NP40 in 10 mM Tris, pH 7.4, 150 mM NaCl) in a Precellys homogenizator (VWR) and 10 μg total protein per sample were proteolyzed with Lys-C and trypsin using a modified FASP procedure (71).

LC-MS/MS analysis was performed on a Q Exactive HF mass spectrometer (Thermo Fisher Scientific) online coupled to a nano-RSLC (Ultimate 3000 RSLC; Dionex). Tryptic peptides were accumulated on a nano trap column (Acclaim PepMap 100 C18, 5 μm, 100 Å, 300 μm inner diameter (i.d.) × 5 mm; Thermo Fisher Scientific) at a flow rate of 30 μl/min followed by separation by reversed phase chromatography (μPAC™ column, 200 cm length, with pillar array backbone at interpillar distance of 2.5 μm, PharmaFluidics) using a non-linear gradient for 240 min from 3 to 42% buffer B (acetonitrile [v/v]/0.1% formic acid [v/v] in HPLC-grade water) in buffer A (2% acetonitrile [v/v]/0.1% formic acid [v/v] in HPLC-grade water) at a flow rate of 300 nl/min. MS spectra were recorded at a resolution of 60,000 with an AGC target of 3 × 106 and a maximum injection time of 50 ms, at a range of 300 to 1500 m/z. From the MS scan, the 10 most abundant ions were selected for HCD fragmentation with a normalized collision energy of 27, an isolation window of 1.6 m/z, and a dynamic exclusion of 30 s. MS/MS spectra were recorded at a resolution of 15,000 with an AGC target of 105 and a maximum injection time of 50 ms.

### STATISTICAL ANALYSIS

#### General image analysis and statistics

Images were analyzed with ZEN (Zeiss) and ImageJ software. Cell countings were performed with the Cell Counter plug-in for ImageJ by careful inspection across serial optical sections (spaced at 1 μm interval) of confocal Z-stacks acquired with a 40x objective (NA 1.1). Image processing was performed with ImageJ after stitching in Imaris Stitcher (9.6.0, Bitplane) when needed and multipanel figures assembled in Adobe Photoshop/Illustrator (Adobe Systems).

Statistical analysis was performed using GraphPad Prism Version 8.0 Software (Graphpad). All biological replicates (n, mice) are derived from at least 2 independent experiments (see figure legends for details on each experiment). Values are reported as mean ± S.E.M. calculated between different mice. Statistical significance was defined at *p < 0.05, **p < 0.01, ***p < 0.001, and ****p < 0.0001. For comparison of connectivity ratio or graft volume analysis between two conditions non-parametric Mann-Whitney and Kruskal-Wallis one-way analysis of variance tests were used. Data visualization leveraged Microsoft Excel, GraphPad Prism Version 8.0 and Adobe Illustrator 2020.

Statistical significance for the proteomics experiment was ascertained employing the approach described in Navarro *et al*. (72), which is based on the presumption that we look for expression changes of proteins that are just a few in comparison to the number of total proteins being quantified. The quantification variability of the non-changing “background” proteins can be used to infer which proteins change their expression in a statistically significant manner. Proteins with p < 0.05 were considered significantly changed.

#### Connectivity analysis

Whole slice tilescans were used to identify brain regions with GFP (or mCherry) labeled cells, by alignment with the corresponding sections of the Allen Reference Atlas of the adult mouse brain (version 2; 2011; Allen Institute for Brain Science). Some sections of interest are not available in this reference atlas, namely in the sagittal atlas, which displays 21 sections spaced at 200 μm intervals, and only up to 4.0 mm lateral from bregma. In these cases, the Brain Explorer 2 software (Allen Institute for Brain Science) was used to retrieve the corresponding annotated section and overlap it with the experimental section to identify the anatomical location of the labeled cells. In sections with unclear cell numbers analysis of the confocal Z-stacks was carried out, and quantification was performed by careful inspection through serial optical sections spaced at 1 μm interval. In sections including transplanted cells, four categories were considered for counting: GFP-only (or mCherry-only) with neuronal morphology, GFP-only (or mCherry-only) with glial morphology, RFP/GFP (or GFP/mCherry) cells with neuronal morphology so called “starter” neurons, and RFP/GFP (or GFP/mCherry) cells with glial morphology. Connectivity ratio for a given anatomical region was calculated by computing the ratio of the total number of GFP-only cells with neuronal morphology counted in that region and the total number of GFP/RFP cells with neuronal morphology in V1, so called “starter cells” (or mCherry-only neurons in a region per number of starter GFP/mCherry neurons in V1). Results are represented as mean ± S.E.M. calculated between different mice.

Important technical consideration of this system relates with intragraft connectivity. Note that a few double labeled cells may have acquired the rabies virus *via* a synapse with another transplanted cell, instead of being primarily infected. However, as these express the rabies G, they are also starter neurons that propagate the virus brain-wide, and thus, this aspect does not interfere with our analysis. Moreover, due to a transduction rate *in vitro* lower than 100%, some transplanted cells are unlabeled. In case these cells innervate a starter cell they’ll be labeled GFP and accounted for in the “Vis” connectivity. Importantly, a minority of the GFP-only presynaptic cells counted in the full visual area are within the transplant and thus neglectable, as it can be appreciated in Fig. 2D and 5B. Note that the graft is only 250-300 um of diameter, while the visual cortex is about 2.5 mm wide (medio-lateral) and 2.5 mm long (anteroposterior).

#### Proteomic data processing – Label-free quantification

The individual raw-files were loaded to the Proteome Discoverer 2.4 software (Thermo Fisher Scientific; version 2.4.1.15) allowing for peptide identification and label-free quantification using the Minora node. Database search (Sequest HT search engine) were performed using Sequest HT as search engine in the Swiss Prot database, taxonomy mouse (17038 sequences) with the following search settings: 10 ppm precursor tolerance, 0.02 Da fragment tolerance, full tryptic specificity, two missed cleavages allowed, carbamidomethyl on cysteine as fixed modification, deamidation of glutamine and asparagine allowed as variable modifications, as well as oxidation of methionine and Met-loss combined with acetylation at the N-terminus of the protein. Carbamidomethylation of Cys was set as a static modification. Dynamic modifications included deamidation of Asn and Gln, oxidation of Met, and a combination of Met loss with acetylation on protein N-terminus. The Percolator node was used for validating peptide spectrum matches and peptides, accepting only the top-scoring hit for each spectrum, and satisfying a false discovery rate (FDR) <1% (high confidence). Protein groups were additionally filtered for an identification FDR <5% (target/decoy concatenated search validation). Peak intensities (at RT apex) of all allocated unique peptides were used for pairwise ratio calculations. A background-based t-test was employed for calculation of statistical significance of the reported ratios. The final list of proteins complied with the strict parsimony principle.

Peak intensities (at RT apex) for top 3 unique peptides were used for pairwise ratio calculations. Abundance values were normalized to the total peptide amount to account for sample load errors. The protein abundances were calculated summing the abundance values for admissible peptides. The final protein ratio was calculated using median peptide ratios of at least 10 biological replicates each (20 replicates intact, 10 replicates SW, and 10 replicates LPS). P-values were calculated based on the approach described (72). Data was filtered to ensure direct identifications (not based on match-between run) in at least 30 % of samples within at least one experimental group. To visualize the data, volcano plots with log2 abundance ratios of sample replicates of SW and LPS versus intact control and the corresponding log10 p-values were created with Microsoft Excel. For gene ontology (GO) enrichment, significantly differentially expressed up- or downregulated proteins were run against a background list of all detected proteins using the webserver GO-rilla (http://cbl-gorilla.cs.technion.ac.il/; *(73)*. A heatmap of all significantly regulated proteins of the SW condition was created with GraphPad Prism Version 8.0, by plotting all significantly regulated proteins in SW condition and subsequently these were aligned with the corresponding proteins of the LPS condition.

## Supporting information

Supplementary Data S1 - Connectivity analysis

Supplementary Data S2a - SW vs Intact

Supplementary Data S2b - GO terms SW

Supplementary Data S2c - LPS vs Intact

Supplementary Data S2d - GO terms LPS

Supplementary Data S2e - SW-specific proteins

## Acknowledgments

We are grateful for technical assistance by Tatiana Simon-Ebert and Detlef Franzen. We also greatly appreciate the expert input by Jacob Kjell and Miriam Esgleas in the proteome analysis. We are particularly thankful to Maria Fernanda Martinez-Reza for performing a crucial experiment on testing inflammatory stimuli in regard to regulation of the G protein which we included in the accompanying manuscript by Thomas et al..

## Author contributions

M.G. initially conceived the idea. M.G. and S.G. conceived and coordinated the project. S.G. designed, performed and analyzed all experiments except the proteomics. J.T. and S.M.H. conducted the proteome study: J.T. performed injuries, collected samples and post-analyzed the data. S.M.H. prepared the samples, performed measurements and quantitative analysis. For the proteome validation, glial reactivity analysis and characterization of the identity of transplanted cells, M.T. and S.G performed LPS and SW surgeries, and Y.Z. and S.G. performed SW followed by transplantation for the 5dpt, 2wpt and 4wpt. M.T. and S.G stained sections and M.T., Y.Z. and S.G. acquired images. S.G. quantified fibrinogen/C1q as well as GFAP/Iba1 while Y.Z. quantified Cux1 transplant composition. K.-K.C. provided the expertise and viral vectors for monosynaptic tracing. S.G. and M.G. wrote the manuscript with input from all co-authors.

## Data and materials availability

All data needed to evaluate the conclusions in the paper are present in the paper and/or the Supplementary Materials. The mass spectrometry proteomics data have been deposited to the ProteomeXchange Consortium via the PRIDE (74) partner repository with the dataset identifier PXD023660 and 10.6019/PXD023660.

