## Supplementary Figures and Table S1 for "Brain injury environment critically influences the connectivity of transplanted neurons"

**Fig. S1.**

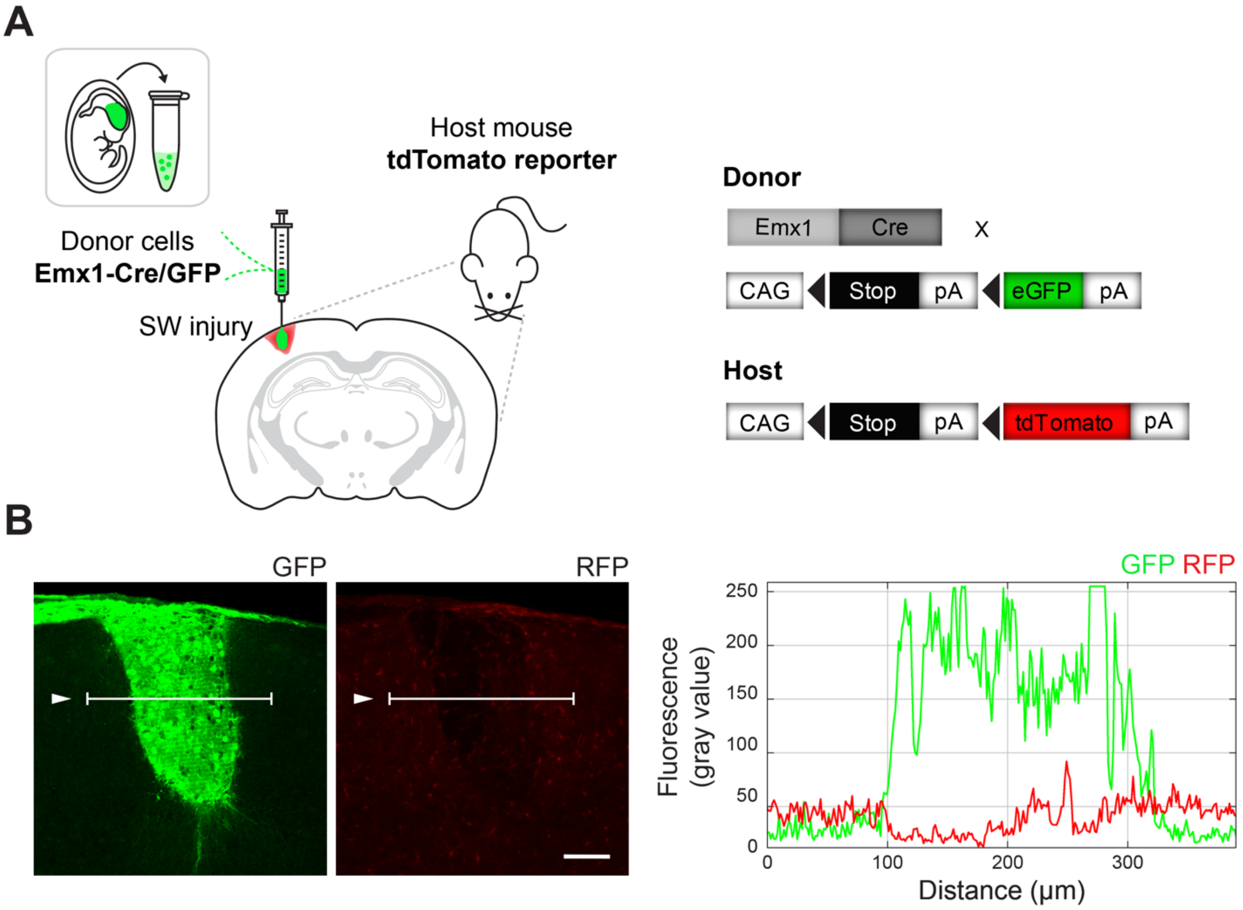


**Control for cell fusion between host and graft cells.**
(**A**) Genetic strategy: Emx1-Cre/GFP cells were transplanted into the SW-injured visual cortex of tdTomato reporter mice (n=4). (**B**) Confocal images and fluorescence intensity analysis along a line drawn across the transplant show absence of tdTomato fluorescence in GFP transplants. Scale bar: (B) 100 µm.

**Fig. S2.**

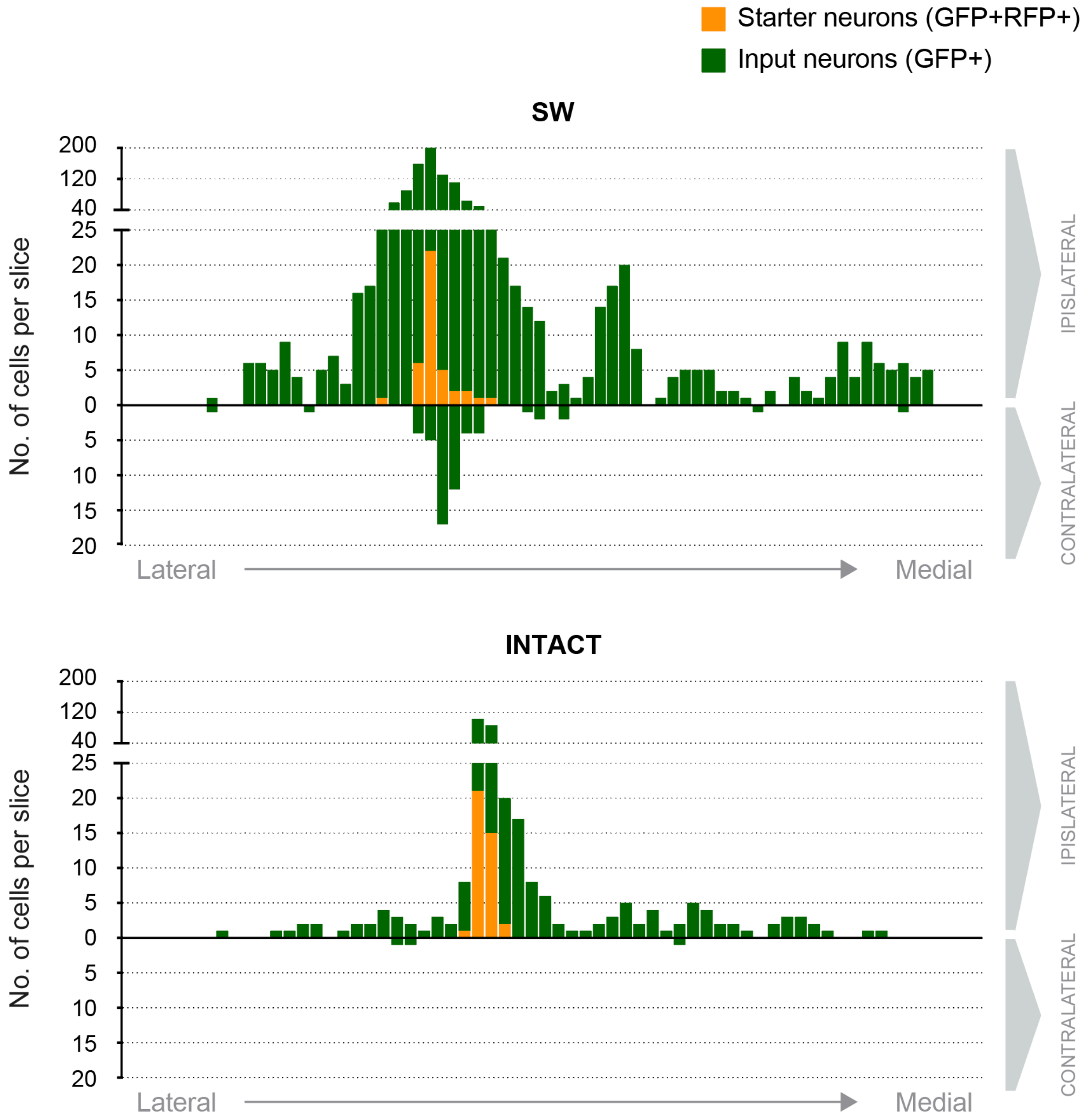


**Brain-wide distribution of input neurons in SW and intact cortex, at 4 wpt.** Example from both experimental groups shows (y axis) the number of input neurons (GFP-only; in green) and "starter" neurons (double-labelled GFP/RFP; in yellow) throughout the brain. Each bar corresponds to one sagittal section, from the most lateral (left) to the most medial (right) in the mouse brain (x axis). The transplanted brain hemisphere (ipsilateral) and the contralateral are represented above and below the x axis respectively. Note the overrepresentation of local connections compared to long-range in both conditions, and the scarce amount of input neurons for transplants in the intact brain, for a comparable number of “starter” neurons.

**Fig. S3.**

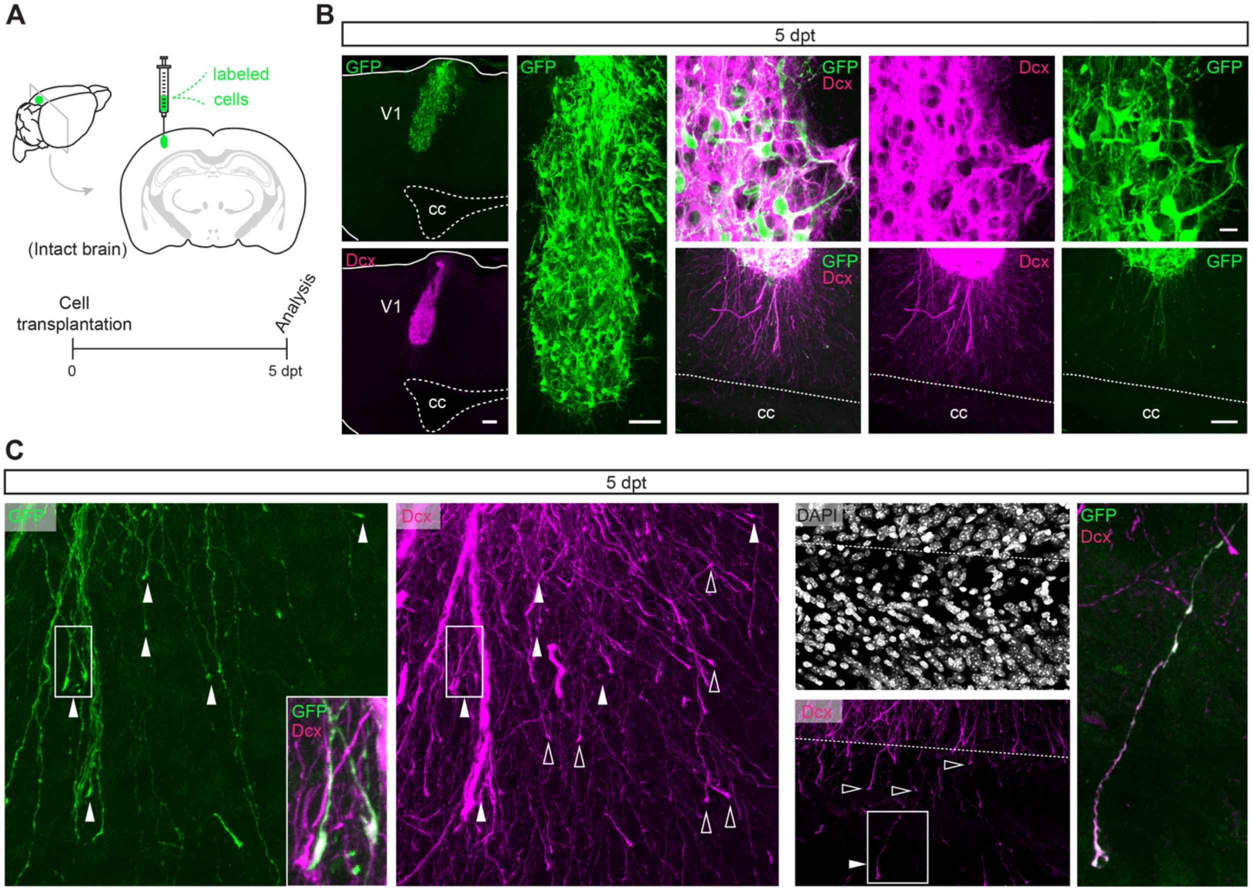


**Early development of transplanted neurons in the intact brain.** (**A**) Experimental procedure and time of analysis. (**B-C**) Confocal images of GFP-labeled neuronal transplants at 5 days after transplantation (dpt) immunostained for the immature neuronal marker Dcx and DAPI for nuclear labeling (the latter only in C). (**B**) Note the immature nature of the transplant, largely Dcx-positive, but robust outgrowth of neurites. (**C**) Neurites display growth cone like structures at their tips (see white and empty arrowheads for GFP+/Dcx+ and GFP-/Dcx+ growth cones respectively) and some have reached the corpus callosum (cc; evidenced by a distinct alignment and density of the DAPI nuclei characteristic of the white matter). Boxed areas are magnified in insets on the right. Scale bars: (B) left, 100 µm; right, 10 and 50 µm (top/bottom).

Fig. S4.

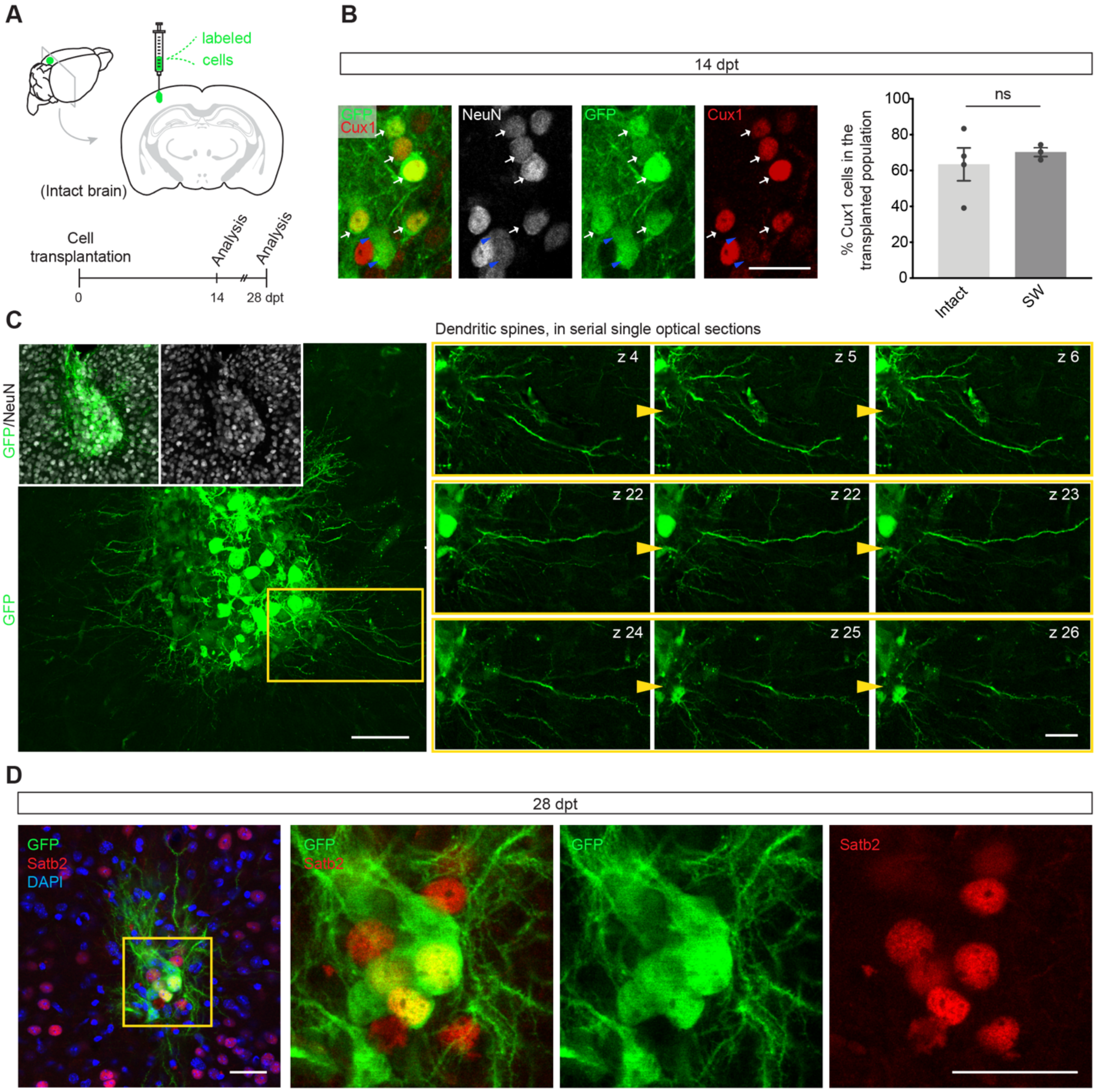


**Transplanted neurons express cortical markers and develop dendritic spines in the intact brain.** (**A**) Experimental procedure and time of analysis. (**B**) High magnification confocal images within the transplant show that GFP grafted cells express the mature neuronal marker NeuN and a large fraction co-express the marker of upper layer cortical identity Cux1 (white arrows; blue arrows for GFP+/NeuN+/Cux1- cells) at 14 dpt (n=3/4 for SW and intact respectively; ns, non-significant, using Mann-Whitney test). (**C**) At this time, transplanted neurons already bear spines in their dendrites. Left, low magnification confocal images show an example of NeuN+ transplant at 14dpt with the yellow boxed area magnified on the right. Right, each set of 3 serial optical sections shows a different dendrite where individual spines are visible across the three planes) (**D**) Confocal images show that some of the GFP grafted cells also express the callosal projection marker Satb2; yellow boxed area is zoomed on the right. Scale bars: (A) 25 μm (C) left, 50 μm, right, 20 μm. (D) left, 50 μm, right, 20 μm.

**Fig. S5.**

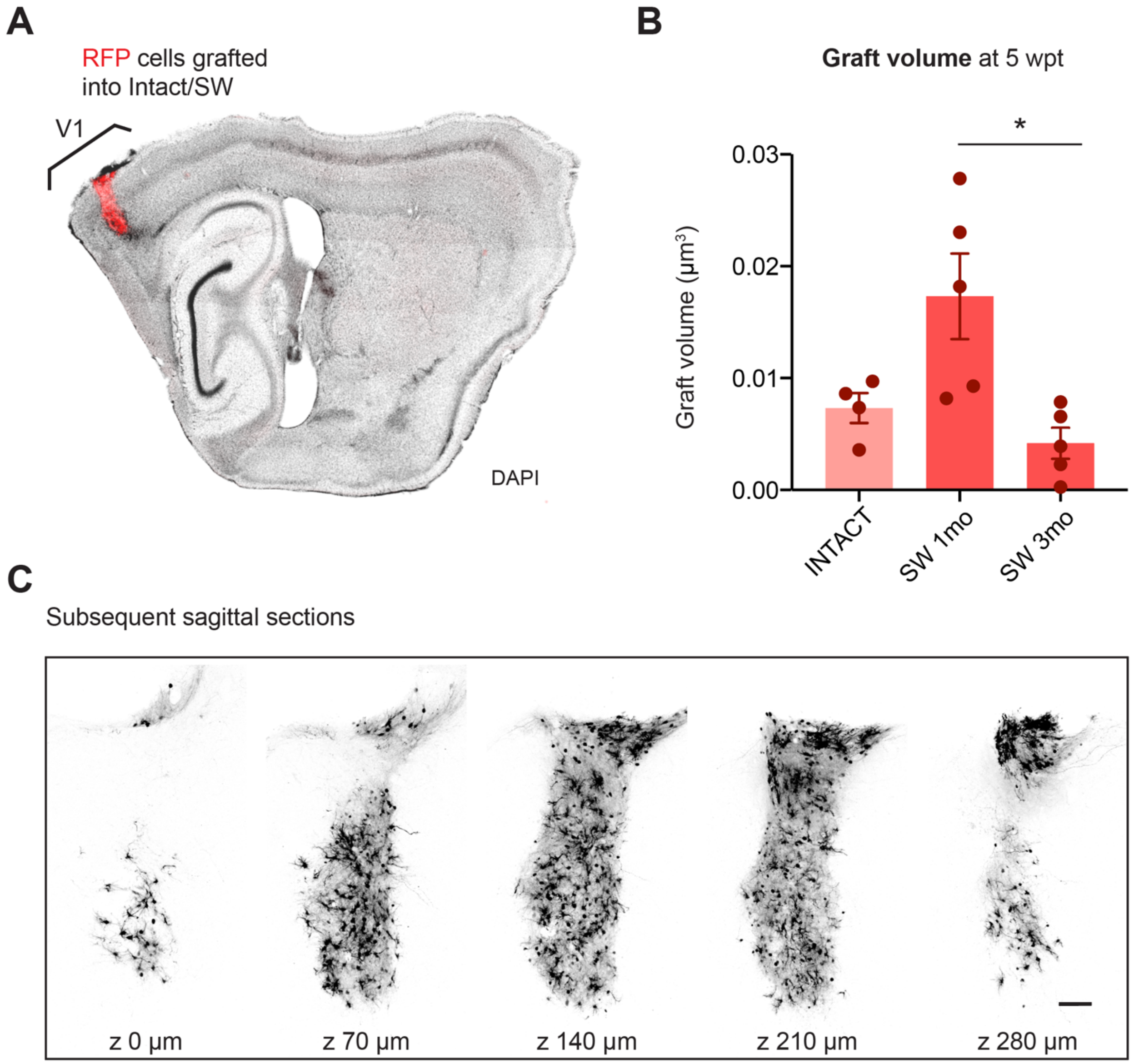


**Analysis of the graft size in SW and intact cortex.** (**A**) Sagittal section from a mouse brain transplanted with RFP-labeled cells in the primary visual cortex (V1), either intact or previously inflicted with a SW. (**B**) Graft volume calculated by summing the graft volumes per slice: the area of graft (RFP area) per slice is measured in Fiji and multiplied by the slice thickness. (n=4 for intact and n=5 for both SW 1mpt and 3mpt). (**C**) Sections with clear RFP cells within the cortical parenchyma were considered for the analysis, and those with cells above L1 or meningeal pia that result from leakage or reflux while retracting the transplantation needle were excluded. *$p < 0.05$ using Kruskal-Wallis multiple comparison test.

**Fig. S6.**

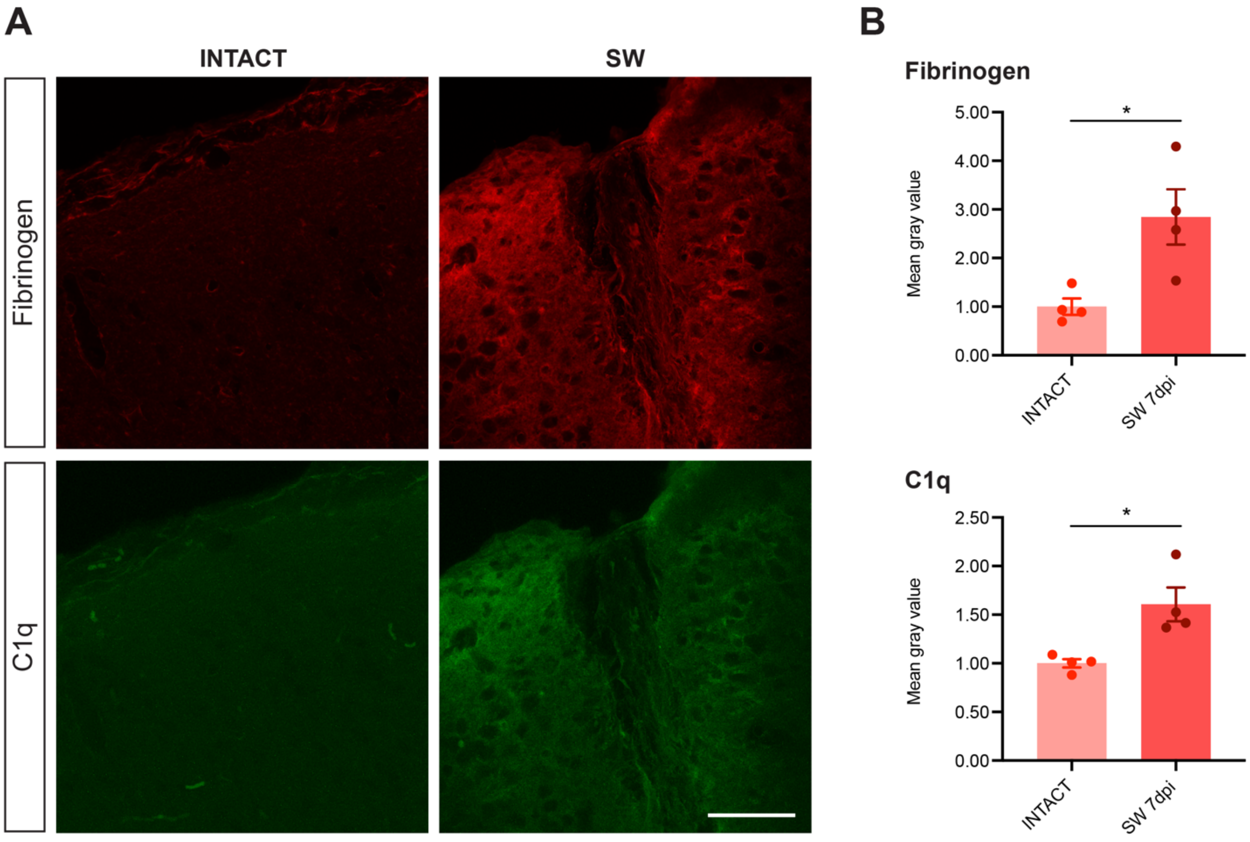


**Increased levels of fibrinogen and C1q in SW at 7dpi.** (**A**) Representative confocal image of the SW-inflicted (7dpi) or intact visual cortex stained for fibrinogen and C1q. Note the pronounced increase in immunopositive signal in SW verified by (**B**) quantification of the mean gray value of all pixels, calculated using Z-stack projections of similar thickness (n=4; note that the same area size and equivalent position in the visual cortex were selected for quantification). *$p < 0.05$ using Mann-Whitney test.

**Table S1.**

| Abbreviation | Anatomical region |
|---|---|
| AMY | Amygdala |
| Aud | Auditory cortex |
| Cg | Cingulate cortex |
| DBN | Diagonal band nucleus |
| ECT | Ectorhinal cortex |
| ENT | Entorhinal cortex |
| GP | Globus pallidus |
| HLZ | Hypothalamic lateral zone |
| Hipp | Hippocampus |
| HPF | Hippocampal formation |
| MB | Midbrain |
| MO | Motor cortex |
| Orb | Orbital cortex |
| Prh | Perirhinal cortex |
| Pons | Pons |
| PtPa | Posterior parietal association area |
| RS | Retrosplenial cortex |
| SI | Substantia innominata |
| SS | Somatosensory cortex |
| TEa | Temporal association areas |
| Thal dLGN | Thalamic dorsal lateral geniculate nucleus |
| Thal other | Other thalamic nucleus |
| Vis | Visual cortex |

Abbreviations of anatomical regions.

**Movie S1.**
Z-series stack shows the complex neuronal morphology 5 weeks after transplantation in a cortical stab injury.

**Data S1. (separate file)**
Absolute number of starter and pre-synaptic neurons counted per brain, for all the experimental groups.

**Data S2. (separate file)**
Protein and GO term enrichment analysis.
