## Supplementary Data S2b - GO terms SW for "Brain injury environment critically influences the connectivity of transplanted neurons"

### enriched GO terms of increased proteins (biological process)

| GO Term | Description | P-value | FDR q-value | Enrichment | N | B | n | b | Genes |
|---|---|---|---|---|---|---|---|---|---|
| GO:0072376 | protein activation cascade | 5.01E-20 | 5.7E-16 | 20,17 | 4935 | 26 | 160 | 17 | [Igkc - immunoglobulin kappa constant, Krt1 - keratin 1, C3 - complement component 3, C4b - complement component 4b (chido blood group), Fn1 - fibronectin 1, Cfb - complement factor b, C1qc - complement component 1, q subcomponent, c chain, C1qa - complement component 1, q subcomponent, alpha polypeptide, C1qb - complement component 1, q subcomponent, beta polypeptide, Apoh - apolipoprotein h, Ighg2b - immunoglobulin heavy constant gamma 2b, Ighg1 - immunoglobulin heavy constant gamma 1 (g1m marker), Fga - fibrinogen alpha chain, Fgg - fibrinogen gamma chain, Ighm - immunoglobulin heavy constant mu, Cfh - complement component factor h, Fgb - fibrinogen beta chain] |
| GO:0006959 | humoral immune response | 5.1E-16 | 2.9E-12 | 14,95 | 4935 | 33 | 160 | 16 | [C3 - complement component 3, Krt1 - keratin 1, Igkc - immunoglobulin kappa constant, C4b - complement component 4b (chido blood group), Cfb - complement factor b, C1qc - complement component 1, q subcomponent, c chain, F2 - coagulation factor ii, C1qa - complement component 1, q subcomponent, alpha polypeptide, Hrg - histidine-rich glycoprotein, C1qb - complement component 1, q subcomponent, beta polypeptide, Ighg2b - immunoglobulin heavy constant gamma 2b, Ighg1 - immunoglobulin heavy constant gamma 1 (g1m marker), Fga - fibrinogen alpha chain, Ighm - immunoglobulin heavy constant mu, Cfh - complement component factor h, Fgb - fibrinogen beta chain] |
| GO:0052547 | regulation of peptidase activity | 2.45E-15 | 9.28E-12 | 6,12 | 4935 | 141 | 160 | 28 | [Pzp - pregnancy zone protein, Serpina1b - serine (or cysteine) preptidase inhibitor, clade a, member 1b, Serpina6 - serine (or cysteine) peptidase inhibitor, clade a, member 6, Serpina1a - serine (or cysteine) peptidase inhibitor, clade a, member 1a, Ahsg - alpha-2-hs-glycoprotein, Serpinh1 - serine (or cysteine) peptidase inhibitor, clade h, member 1, Gsn - gelsolin, Cst3 - cystatin c, Serpina1d - serine (or cysteine) peptidase inhibitor, clade a, member 1d, Serpina1e - serine (or cysteine) peptidase inhibitor, clade a, member 1e, Serpinc1 - serine (or cysteine) peptidase inhibitor, clade c (antithrombin), member 1, Lgmn - legumain, Serpina3k - serine (or cysteine) peptidase inhibitor, clade a, member 3k, Ctsd - cathepsin d, Kng1 - kininogen 1, Hrg - histidine-rich glycoprotein, Psme1 - proteasome (prosome, macropain) activator subunit 1 (pa28 alpha), Ddrgk1 - ddrgk domain containing 1, Vtn - vitronectin, Itih3 - inter-alpha trypsin inhibitor, heavy chain 3, Serpinb1a - serine (or cysteine) peptidase inhibitor, clade b, member 1a, Itih2 - inter-alpha trypsin inhibitor, heavy chain 2, Itih1 - inter-alpha trypsin inhibitor, heavy chain 1, Pdcd5 - programmed cell death 5, Ncstn - nicastrin, Mug1 - murinoglobulin 1, Stat3 - signal transducer and activator of transcription 3, Stat1 - signal transducer and activator of transcription 1] |
| GO:0002376 | immune system process | 8.95E-15 | 2.55E-11 | 3,78 | 4935 | 343 | 160 | 42 | [Scg2 - secretogranin ii, Anxa3 - annexin a3, Plg - plasminogen, Aif1 - allograft inflammatory factor 1, Lcp1 - lymphocyte cytosolic protein 1, Irgm1 - immunity-related gtpase family m member 1, Fgg - fibrinogen gamma chain, Fcer1g - fc receptor, ige, high affinity i, gamma polypeptide, Cfh - complement component factor h, Sod1 - superoxide dismutase 1, soluble, Fgb - fibrinogen beta chain, Ide - insulin degrading enzyme, C3 - complement component 3, Krt1 - keratin 1, Igkc - immunoglobulin kappa constant, H2-D1 - histocompatibility 2, d region locus 1, C4b - complement component 4b (chido blood group), Itgam - integrin alpha m, Lgals1 - lectin, galactose binding, soluble 1, Hp - haptoglobin, Cfb - complement factor b, C1qc - complement component 1, q subcomponent, c chain, Itga6 - integrin alpha 6, F2 - coagulation factor ii, C1qa - complement component 1, q subcomponent, alpha polypeptide, C1qb - complement component 1, q subcomponent, beta polypeptide, Hrg - histidine-rich glycoprotein, Psme1 - proteasome (prosome, macropain) activator subunit 1 (pa28 alpha), Vtn - vitronectin, Itgb2 - integrin beta 2, Gba - glucosidase, beta, acid, Sema4a - sema domain, immunoglobulin domain (ig), transmembrane domain (tm) and short cytoplasmic domain, (semaphorin) 4a, Ighg2b - immunoglobulin heavy constant gamma 2b, Ighg1 - immunoglobulin heavy constant gamma 1 (g1m marker), Fga - fibrinogen alpha chain, Ampd3 - adenosine monophosphate deaminase 3, Ighm - immunoglobulin heavy constant mu, Ncstn - nicastrin, Clu - clusterin, Stat3 - signal transducer and activator of transcription 3, Cd151 - cd151 antigen, Stat1 - signal transducer and activator of transcription 1] |
| GO:0006956 | complement activation | 1.4E-14 | 3.2E-11 | 20,56 | 4935 | 18 | 160 | 12 | [Igkc - immunoglobulin kappa constant, Krt1 - keratin 1, C3 - complement component 3, C4b - complement component 4b (chido blood group), Cfb - complement factor b, Ighg2b - immunoglobulin heavy constant gamma 2b, Ighg1 - immunoglobulin heavy constant gamma 1 (g1m marker), C1qc - complement component 1, q subcomponent, c chain, Ighm - immunoglobulin heavy constant mu, Cfh - complement component factor h, C1qa - complement component 1, q subcomponent, alpha polypeptide, C1qb - complement component 1, q subcomponent, beta polypeptide] |
| GO:0030162 | regulation of proteolysis | 1.13E-13 | 2.14E-10 | 4,07 | 4935 | 273 | 160 | 36 | [Serpina1b - serine (or cysteine) preptidase inhibitor, clade a, member 1b, Pzp - pregnancy zone protein, Serpina6 - serine (or cysteine) peptidase inhibitor, clade a, member 6, Serpina1a - serine (or cysteine) peptidase inhibitor, clade a, member 1a, Ahsg - alpha-2-hs-glycoprotein, Serpinh1 - serine (or cysteine) peptidase inhibitor, clade h, member 1, Gsn - gelsolin, Cst3 - cystatin c, Serpina1d - serine (or cysteine) peptidase inhibitor, clade a, member 1d, Serpina1e - serine (or cysteine) peptidase inhibitor, clade a, member 1e, Lgmn - legumain, Serpinc1 - serine (or cysteine) peptidase inhibitor, clade c (antithrombin), member 1, Serpina3k - serine (or cysteine) peptidase inhibitor, clade a, member 3k, Fhit - fragile histidine triad gene, Cfh - complement component factor h, Ctsd - cathepsin d, Kng1 - kininogen 1, Ide - insulin degrading enzyme, C3 - complement component 3, F2 - coagulation factor ii, Psme1 - proteasome (prosome, macropain) activator subunit 1 (pa28 alpha), Hrg - histidine-rich glycoprotein, Ddrgk1 - ddrgk domain containing 1, Vtn - vitronectin, Gba - glucosidase, beta, acid, Itih3 - inter-alpha trypsin inhibitor, heavy chain 3, Serpinb1a - serine (or cysteine) peptidase inhibitor, clade b, member 1a, Itih2 - inter-alpha trypsin inhibitor, heavy chain 2, Itih1 - inter-alpha trypsin inhibitor, heavy chain 1, Pdcd5 - programmed cell death 5, Clu - clusterin, Ncstn - nicastrin, Commd1 - comm domain containing 1, Stat3 - signal transducer and activator of transcription 3, Mug1 - murinoglobulin 1, Stat1 - signal transducer and activator of transcription 1] |
| GO:0010466 | negative regulation of peptidase activity | 1.36E-12 | 2.21E-9 | 7,71 | 4935 | 76 | 160 | 19 | [Pzp - pregnancy zone protein, Serpina1b - serine (or cysteine) preptidase inhibitor, clade a, member 1b, Serpina6 - serine (or cysteine) peptidase inhibitor, clade a, member 6, Serpina1a - serine (or cysteine) peptidase inhibitor, clade a, member 1a, Ahsg - alpha-2-hs-glycoprotein, Serpinh1 - serine (or cysteine) peptidase inhibitor, clade h, member 1, Cst3 - cystatin c, Hrg - histidine-rich glycoprotein, Vtn - vitronectin, Serpina1d - serine (or cysteine) peptidase inhibitor, clade a, member 1d, Serpina1e - serine (or cysteine) peptidase inhibitor, clade a, member 1e, Serpinc1 - serine (or cysteine) peptidase inhibitor, clade c (antithrombin), member 1, Itih3 - inter-alpha trypsin inhibitor, heavy chain 3, Serpinb1a - serine (or cysteine) peptidase inhibitor, clade b, member 1a, Serpina3k - serine (or cysteine) peptidase inhibitor, clade a, member 3k, Itih2 - inter-alpha trypsin inhibitor, heavy chain 2, Itih1 - inter-alpha trypsin inhibitor, heavy chain 1, Mug1 - murinoglobulin 1, Kng1 - kininogen 1] |

| | | | | | | | | | |
|---|---|---|---|---|---|---|---|---|---|
| GO:0006952 | defense response | 2.8E-12 | 3.99E-9 | 4,36 | 4935 | 212 | 160 | 30 | [Serpina1b - serine (or cysteine) preptidase inhibitor, clade a, member 1b, Anxa3 - annexin a3, Ahsg - alpha-2-hs-glycoprotein, Fn1 - fibronectin 1, Cst3 - cystatin c, Irgm1 - immunity-related gtpase family m member 1, Fgg - fibrinogen gamma chain, Fcer1g - fc receptor, ige, high affinity i, gamma polypeptide, Cfh - complement component factor h, Kng1 - kininogen 1, Fgb - fibrinogen beta chain, C3 - complement component 3, Igkc - immunoglobulin kappa constant, Krt1 - keratin 1, Hp - haptoglobin, Cfb - complement factor b, C1qc - complement component 1, q subcomponent, c chain, F2 - coagulation factor ii, C1qa - complement component 1, q subcomponent, alpha polypeptide, C1qb - complement component 1, q subcomponent, beta polypeptide, Hrg - histidine-rich glycoprotein, Ighg2b - immunoglobulin heavy constant gamma 2b, Serpinb1a - serine (or cysteine) peptidase inhibitor, clade b, member 1a, Orm1 - orosomucoid 1, Ighg1 - immunoglobulin heavy constant gamma 1 (g1m marker), Fga - fibrinogen alpha chain, Ighm - immunoglobulin heavy constant mu, Stat3 - signal transducer and activator of transcription 3, Tspan2 - tetraspanin 2, Stat1 - signal transducer and activator of transcription 1] |
| GO:0045861 | negative regulation of proteolysis | 7.43E-12 | 9.39E-9 | 5,59 | 4935 | 127 | 160 | 23 | [Pzp - pregnancy zone protein, Serpina1b - serine (or cysteine) preptidase inhibitor, clade a, member 1b, Serpina6 - serine (or cysteine) peptidase inhibitor, clade a, member 6, Serpina1a - serine (or cysteine) peptidase inhibitor, clade a, member 1a, Ahsg - alpha-2-hs-glycoprotein, Serpinh1 - serine (or cysteine) peptidase inhibitor, clade h, member 1, F2 - coagulation factor ii, Cst3 - cystatin c, Hrg - histidine-rich glycoprotein, Ddrgk1 - ddrgk domain containing 1, Vtn - vitronectin, Serpina1d - serine (or cysteine) peptidase inhibitor, clade a, member 1d, Serpina1e - serine (or cysteine) peptidase inhibitor, clade a, member 1e, Serpinb1a - serine (or cysteine) peptidase inhibitor, clade b, member 1a, Itih3 - inter-alpha trypsin inhibitor, heavy chain 3, Serpinc1 - serine (or cysteine) peptidase inhibitor, clade c (antithrombin), member 1, Itih2 - inter-alpha trypsin inhibitor, heavy chain 2, Serpina3k - serine (or cysteine) peptidase inhibitor, clade a, member 3k, Fhit - fragile histidine triad gene, Itih1 - inter-alpha trypsin inhibitor, heavy chain 1, Mug1 - murinoglobulin 1, Kng1 - kininogen 1, Ide - insulin degrading enzyme] |
| GO:0002252 | immune effector process | 1.61E-11 | 1.83E-8 | 6,36 | 4935 | 97 | 160 | 20 | [Krt1 - keratin 1, Igkc - immunoglobulin kappa constant, C3 - complement component 3, Anxa3 - annexin a3, Lgals1 - lectin, galactose binding, soluble 1, C4b - complement component 4b (chido blood group), Cfb - complement factor b, C1qc - complement component 1, q subcomponent, c chain, F2 - coagulation factor ii, Lcp1 - lymphocyte cytosolic protein 1, C1qa - complement component 1, q subcomponent, alpha polypeptide, C1qb - complement component 1, q subcomponent, beta polypeptide, Sema4a - sema domain, immunoglobulin domain (ig), transmembrane domain (tm) and short cytoplasmic domain, (semaphorin) 4a, Ighg2b - immunoglobulin heavy constant gamma 2b, Ighg1 - immunoglobulin heavy constant gamma 1 (g1m marker), Fcer1g - fc receptor, ige, high affinity i, gamma polypeptide, Ighm - immunoglobulin heavy constant mu, Clu - clusterin, Cfh - complement component factor h, Stat1 - signal transducer and activator of transcription 1] |
| GO:0032101 | regulation of response to external stimulus | 5.73E-11 | 5.93E-8 | 3,89 | 4935 | 238 | 160 | 30 | [Scg2 - secretogranin ii, Plg - plasminogen, Fn1 - fibronectin 1, Ahsg - alpha-2-hs-glycoprotein, Aif1 - allograft inflammatory factor 1, Flna - filamin, alpha, Apoh - apolipoprotein h, Apod - apolipoprotein d, Lgmn - legumain, Serpinc1 - serine (or cysteine) peptidase inhibitor, clade c (antithrombin), member 1, Fgg - fibrinogen gamma chain, Fcer1g - fc receptor, ige, high affinity i, gamma polypeptide, Cfh - complement component factor h, Sod1 - superoxide dismutase 1, soluble, Kng1 - kininogen 1, Fgb - fibrinogen beta chain, C3 - complement component 3, Krt1 - keratin 1, Fabp7 - fatty acid binding protein 7, brain, F2 - coagulation factor ii, Hrg - histidine-rich glycoprotein, Sema4a - sema domain, immunoglobulin domain (ig), transmembrane domain (tm) and short cytoplasmic domain, (semaphorin) 4a, Tgm2 - transglutaminase 2, c polypeptide, Ighg2b - immunoglobulin heavy constant gamma 2b, Ighg1 - immunoglobulin heavy constant gamma 1 (g1m marker), Fga - fibrinogen alpha chain, Apoa1 - apolipoprotein a-i, Stat3 - signal transducer and activator of transcription 3, Stat1 - signal transducer and activator of transcription 1, Anxa2 - annexin a2] |
| GO:0006955 | immune response | 1.93E-10 | 1.83E-7 | 4,79 | 4935 | 148 | 160 | 23 | [C3 - complement component 3, Krt1 - keratin 1, Igkc - immunoglobulin kappa constant, H2-D1 - histocompatibility 2, d region locus 1, C4b - complement component 4b (chido blood group), Cfb - complement factor b, C1qc - complement component 1, q subcomponent, c chain, F2 - coagulation factor ii, C1qa - complement component 1, q subcomponent, alpha polypeptide, C1qb - complement component 1, q subcomponent, beta polypeptide, Hrg - histidine-rich glycoprotein, Vtn - vitronectin, Sema4a - sema domain, immunoglobulin domain (ig), transmembrane domain (tm) and short cytoplasmic domain, (semaphorin) 4a, Irgm1 - immunity-related gtpase family m member 1, Ighg2b - immunoglobulin heavy constant gamma 2b, Fga - fibrinogen alpha chain, Fgg - fibrinogen gamma chain, Ighg1 - immunoglobulin heavy constant gamma 1 (g1m marker), Fcer1g - fc receptor, ige, high affinity i, gamma polypeptide, Ighm - immunoglobulin heavy constant mu, Cfh - complement component factor h, Stat3 - signal transducer and activator of transcription 3, Fgb - fibrinogen beta chain] |
| GO:0042730 | fibrinolysis | 2.58E-10 | 2.26E-7 | 26,99 | 4935 | 8 | 160 | 7 | [Plg - plasminogen, Fga - fibrinogen alpha chain, Fgg - fibrinogen gamma chain, F2 - coagulation factor ii, Hrg - histidine-rich glycoprotein, Fgb - fibrinogen beta chain, Anxa2 - annexin a2] |
| GO:0052548 | regulation of endopeptidase activity | 2.88E-10 | 2.34E-7 | 5,18 | 4935 | 125 | 160 | 21 | [Serpina1b - serine (or cysteine) preptidase inhibitor, clade a, member 1b, Serpina6 - serine (or cysteine) peptidase inhibitor, clade a, member 6, Ahsg - alpha-2-hs-glycoprotein, Serpina1a - serine (or cysteine) peptidase inhibitor, clade a, member 1a, Serpinh1 - serine (or cysteine) peptidase inhibitor, clade h, member 1, Gsn - gelsolin, Psme1 - proteasome (prosome, macropain) activator subunit 1 (pa28 alpha), Hrg - histidine-rich glycoprotein, Vtn - vitronectin, Serpina1d - serine (or cysteine) peptidase inhibitor, clade a, member 1d, Serpina1e - serine (or cysteine) peptidase inhibitor, clade a, member 1e, Lgmn - legumain, Serpinb1a - serine (or cysteine) peptidase inhibitor, clade b, member 1a, Serpinc1 - serine (or cysteine) peptidase inhibitor, clade c (antithrombin), member 1, Serpina3k - serine (or cysteine) peptidase inhibitor, clade a, member 3k, Pdcd5 - programmed cell death 5, Ncstn - nicastrin, Ctsd - cathepsin d, Stat3 - signal transducer and activator of transcription 3, Kng1 - kininogen 1, Stat1 - signal transducer and activator of transcription 1] |
| GO:0002684 | positive regulation of immune system process | 4.26E-10 | 3.23E-7 | 3,95 | 4935 | 211 | 160 | 27 | [Aif1 - allograft inflammatory factor 1, Lamp1 - lysosomal-associated membrane protein 1, Irgm1 - immunity-related gtpase family m member 1, Lgmn - legumain, Fcer1g - fc receptor, ige, high affinity i, gamma polypeptide, Cfh - complement component factor h, Igkc - immunoglobulin kappa constant, Krt1 - keratin 1, C3 - complement component 3, H2-D1 - histocompatibility 2, d region locus 1, Lgals1 - lectin, galactose binding, soluble 1, Itgam - integrin alpha m, C4b - complement component 4b (chido blood group), Cfb - complement factor b, C1qc - complement component 1, q subcomponent, c chain, C1qa - complement component 1, q subcomponent, alpha polypeptide, C1qb - complement component 1, q subcomponent, beta polypeptide, Hrg - histidine-rich glycoprotein, Itgb3 - integrin beta 3, Itgb2 - integrin beta 2, Hpx - hemopexin, Ighg2b - immunoglobulin heavy constant gamma 2b, Ighg1 - immunoglobulin heavy constant gamma 1 (g1m marker), Ighm - immunoglobulin heavy constant mu, Irs2 - insulin receptor substrate 2, Stat3 - signal transducer and activator of transcription 3, Stat1 - signal transducer and activator of transcription 1] |

| | | | | | | | | | |
|---|---|---|---|---|---|---|---|---|---|
| GO:0050817 | coagulation | 4.49E-10 | 3.19E-7 | 14,02 | 4935 | 22 | 160 | 10 | [C3 - complement component 3, Apoh - apolipoprotein h, Plg - plasminogen, Serpinc1 - serine (or cysteine) peptidase inhibitor, clade c (antithrombin), member 1, Fgg - fibrinogen gamma chain, Fga - fibrinogen alpha chain, F2 - coagulation factor ii, Hrg - histidine-rich glycoprotein, Fgb - fibrinogen beta chain, Kng1 - kininogen 1] |
| GO:0007596 | blood coagulation | 4.49E-10 | 3,00E-07 | 14,02 | 4935 | 22 | 160 | 10 | [C3 - complement component 3, Apoh - apolipoprotein h, Plg - plasminogen, Serpinc1 - serine (or cysteine) peptidase inhibitor, clade c (antithrombin), member 1, Fga - fibrinogen alpha chain, Fgg - fibrinogen gamma chain, F2 - coagulation factor ii, Hrg - histidine-rich glycoprotein, Kng1 - kininogen 1, Fgb - fibrinogen beta chain] |
| GO:0007599 | hemostasis | 7.72E-10 | 4.88E-7 | 13,41 | 4935 | 23 | 160 | 10 | [C3 - complement component 3, Apoh - apolipoprotein h, Plg - plasminogen, Serpinc1 - serine (or cysteine) peptidase inhibitor, clade c (antithrombin), member 1, Fgg - fibrinogen gamma chain, Fga - fibrinogen alpha chain, F2 - coagulation factor ii, Hrg - histidine-rich glycoprotein, Fgb - fibrinogen beta chain, Kng1 - kininogen 1] |
| GO:1900047 | negative regulation of hemostasis | 1.2E-9 | 7.18E-7 | 15,42 | 4935 | 18 | 160 | 9 | [Apoh - apolipoprotein h, Plg - plasminogen, Fga - fibrinogen alpha chain, Fgg - fibrinogen gamma chain, F2 - coagulation factor ii, Hrg - histidine-rich glycoprotein, Kng1 - kininogen 1, Fgb - fibrinogen beta chain, Anxa2 - annexin a2] |
| GO:0030195 | negative regulation of blood coagulation | 1.2E-9 | 6.82E-7 | 15,42 | 4935 | 18 | 160 | 9 | [Apoh - apolipoprotein h, Plg - plasminogen, Fga - fibrinogen alpha chain, Fgg - fibrinogen gamma chain, F2 - coagulation factor ii, Hrg - histidine-rich glycoprotein, Fgb - fibrinogen beta chain, Kng1 - kininogen 1, Anxa2 - annexin a2] |
| GO:0050819 | negative regulation of coagulation | 1.2E-9 | 6.49E-7 | 15,42 | 4935 | 18 | 160 | 9 | [Apoh - apolipoprotein h, Plg - plasminogen, Fga - fibrinogen alpha chain, Fgg - fibrinogen gamma chain, F2 - coagulation factor ii, Hrg - histidine-rich glycoprotein, Kng1 - kininogen 1, Fgb - fibrinogen beta chain, Anxa2 - annexin a2] |
| GO:0002526 | acute inflammatory response | 2.61E-9 | 1.35E-6 | 17,62 | 4935 | 14 | 160 | 8 | [Serpina1b - serine (or cysteine) preptidase inhibitor, clade a, member 1b, Ahsg - alpha-2-hs-glycoprotein, Fn1 - fibronectin 1, Hp - haptoglobin, Orm1 - orosomucoid 1, Ighg1 - immunoglobulin heavy constant gamma 1 (g1m marker), F2 - coagulation factor ii, Stat3 - signal transducer and activator of transcription 3] |
| GO:0006958 | complement activation, classical pathway | 2.61E-9 | 1.29E-6 | 17,62 | 4935 | 14 | 160 | 8 | [Igkc - immunoglobulin kappa constant, C3 - complement component 3, Ighg2b - immunoglobulin heavy constant gamma 2b, Ighg1 - immunoglobulin heavy constant gamma 1 (g1m marker), C1qc - complement component 1, q subcomponent, c chain, Ighm - immunoglobulin heavy constant mu, C1qa - complement component 1, q subcomponent, alpha polypeptide, C1qb - complement component 1, q subcomponent, beta polypeptide] |
| GO:1900046 | regulation of hemostasis | 3.09E-9 | 1.46E-6 | 10,28 | 4935 | 33 | 160 | 11 | [Apoh - apolipoprotein h, Plg - plasminogen, Serpinc1 - serine (or cysteine) peptidase inhibitor, clade c (antithrombin), member 1, Fga - fibrinogen alpha chain, Fgg - fibrinogen gamma chain, F2 - coagulation factor ii, Fcer1g - fc receptor, ige, high affinity i, gamma polypeptide, Hrg - histidine-rich glycoprotein, Kng1 - kininogen 1, Fgb - fibrinogen beta chain, Anxa2 - annexin a2] |
| GO:0030193 | regulation of blood coagulation | 3.09E-9 | 1.4E-6 | 10,28 | 4935 | 33 | 160 | 11 | [Apoh - apolipoprotein h, Plg - plasminogen, Serpinc1 - serine (or cysteine) peptidase inhibitor, clade c (antithrombin), member 1, Fga - fibrinogen alpha chain, Fgg - fibrinogen gamma chain, F2 - coagulation factor ii, Fcer1g - fc receptor, ige, high affinity i, gamma polypeptide, Hrg - histidine-rich glycoprotein, Fgb - fibrinogen beta chain, Kng1 - kininogen 1, Anxa2 - annexin a2] |
| GO:0051336 | regulation of hydrolase activity | 3.19E-9 | 1.4E-6 | 2,82 | 4935 | 405 | 160 | 37 | [Pzp - pregnancy zone protein, Serpina1b - serine (or cysteine) preptidase inhibitor, clade a, member 1b, Serpina6 - serine (or cysteine) peptidase inhibitor, clade a, member 6, Ahsg - alpha-2-hs-glycoprotein, Serpina1a - serine (or cysteine) peptidase inhibitor, clade a, member 1a, Serpinh1 - serine (or cysteine) peptidase inhibitor, clade h, member 1, Gsn - gelsolin, Cst3 - cystatin c, Serpina1d - serine (or cysteine) peptidase inhibitor, clade a, member 1d, Serpina1e - serine (or cysteine) peptidase inhibitor, clade a, member 1e, Apoh - apolipoprotein h, Serpinc1 - serine (or cysteine) peptidase inhibitor, clade c (antithrombin), member 1, Lgmn - legumain, Ppp1r1b - protein phosphatase 1, regulatory (inhibitor) subunit 1b, Serpina3k - serine (or cysteine) peptidase inhibitor, clade a, member 3k, Ctsd - cathepsin d, Sod1 - superoxide dismutase 1, soluble, Kng1 - kininogen 1, Itgb1 - integrin beta 1 (fibronectin receptor beta), Itga6 - integrin alpha 6, Arhgap22 - rho gtpase activating protein 22, Hrg - histidine-rich glycoprotein, Psme1 - proteasome (prosome, macropain) activator subunit 1 (pa28 alpha), Ddrgk1 - ddrgk domain containing 1, Vtn - vitronectin, Tpm1 - tropomyosin 1, alpha, Scarb2 - scavenger receptor class b, member 2, Itih3 - inter-alpha trypsin inhibitor, heavy chain 3, Serpinb1a - serine (or cysteine) peptidase inhibitor, clade b, member 1a, Itih2 - inter-alpha trypsin inhibitor, heavy chain 2, Itih1 - inter-alpha trypsin inhibitor, heavy chain 1, Pdcd5 - programmed cell death 5, Ncstn - nicastrin, Apoa1 - apolipoprotein a-i, Mug1 - murinoglobulin 1, Stat3 - signal transducer and activator of transcription 3, Stat1 - signal transducer and activator of transcription 1] |
| GO:0002682 | regulation of immune system process | 3.25E-9 | 1.37E-6 | 3,07 | 4935 | 332 | 160 | 33 | [Aif1 - allograft inflammatory factor 1, Gsn - gelsolin, Lamp1 - lysosomal-associated membrane protein 1, Apod - apolipoprotein d, Lgmn - legumain, Irgm1 - immunity-related gtpase family m member 1, Fcer1g - fc receptor, ige, high affinity i, gamma polypeptide, Cfh - complement component factor h, C3 - complement component 3, Igkc - immunoglobulin kappa constant, Krt1 - keratin 1, H2-D1 - histocompatibility 2, d region locus 1, C4b - complement component 4b (chido blood group), Lgals1 - lectin, galactose binding, soluble 1, Itgam - integrin alpha m, Cfb - complement factor b, C1qc - complement component 1, q subcomponent, c chain, C1qa - complement component 1, q subcomponent, alpha polypeptide, C1qb - complement component 1, q subcomponent, beta polypeptide, Hrg - histidine-rich glycoprotein, Itgb3 - integrin beta 3, Rbp1 - retinol binding protein 1, cellular, Itgb2 - integrin beta 2, Hpx - hemopexin, Orm1 - orosomucoid 1, Ighg2b - immunoglobulin heavy constant gamma 2b, Serpinb1a - serine (or cysteine) peptidase inhibitor, clade b, member 1a, Ighg1 - immunoglobulin heavy constant gamma 1 (g1m marker), Ighm - immunoglobulin heavy constant mu, Apoa1 - apolipoprotein a-i, Irs2 - insulin receptor substrate 2, Stat3 - signal transducer and activator of transcription 3, Stat1 - signal transducer and activator of transcription 1] |
| GO:0006953 | acute-phase response | 3.66E-9 | 1.49E-6 | 21,59 | 4935 | 10 | 160 | 7 | [Serpina1b - serine (or cysteine) preptidase inhibitor, clade a, member 1b, Ahsg - alpha-2-hs-glycoprotein, Fn1 - fibronectin 1, Hp - haptoglobin, Orm1 - orosomucoid 1, F2 - coagulation factor ii, Stat3 - signal transducer and activator of transcription 3] |

| | | | | | | | | | |
|---|---|---|---|---|---|---|---|---|---|
| GO:0051346 | negative regulation of hydrolase activity | 3.72E-9 | 1.46E-6 | 4,53 | 4935 | 143 | 160 | 21 | [Pzp - pregnancy zone protein, Serpina1b - serine (or cysteine) preptidase inhibitor, clade a, member 1b, Serpina6 - serine (or cysteine) peptidase inhibitor, clade a, member 6, Serpina1a - serine (or cysteine) peptidase inhibitor, clade a, member 1a, Ahsg - alpha-2-hs-glycoprotein, Serpinh1 - serine (or cysteine) peptidase inhibitor, clade h, member 1, Cst3 - cystatin c, Hrg - histidine-rich glycoprotein, Vtn - vitronectin, Serpina1d - serine (or cysteine) peptidase inhibitor, clade a, member 1d, Serpina1e - serine (or cysteine) peptidase inhibitor, clade a, member 1e, Serpinb1a - serine (or cysteine) peptidase inhibitor, clade b, member 1a, Itih3 - inter-alpha trypsin inhibitor, heavy chain 3, Serpinc1 - serine (or cysteine) peptidase inhibitor, clade c (antithrombin), member 1, Itih2 - inter-alpha trypsin inhibitor, heavy chain 2, Serpina3k - serine (or cysteine) peptidase inhibitor, clade a, member 3k, Ppp1r1b - protein phosphatase 1, regulatory (inhibitor) subunit 1b, Itih1 - inter-alpha trypsin inhibitor, heavy chain 1, Apoa1 - apolipoprotein a-i, Mug1 - murinoglobulin 1, Kng1 - kininogen 1] |
| GO:0050778 | positive regulation of immune response | 3.95E-9 | 1.5E-6 | 4,75 | 4935 | 130 | 160 | 20 | [Krt1 - keratin 1, Igkc - immunoglobulin kappa constant, C3 - complement component 3, H2-D1 - histocompatibility 2, d region locus 1, C4b - complement component 4b (chido blood group), Itgam - integrin alpha m, Cfb - complement factor b, C1qc - complement component 1, q subcomponent, c chain, C1qa - complement component 1, q subcomponent, alpha polypeptide, C1qb - complement component 1, q subcomponent, beta polypeptide, Hrg - histidine-rich glycoprotein, Lamp1 - lysosomal-associated membrane protein 1, Itgb2 - integrin beta 2, Hpx - hemopexin, Irgm1 - immunity-related gtpase family m member 1, Ighg2b - immunoglobulin heavy constant gamma 2b, Ighg1 - immunoglobulin heavy constant gamma 1 (g1m marker), Fcer1g - fc receptor, ige, high affinity i, gamma polypeptide, Ighm - immunoglobulin heavy constant mu, Cfh - complement component factor h] |
| GO:0050818 | regulation of coagulation | 4.44E-9 | 1.63E-6 | 9,98 | 4935 | 34 | 160 | 11 | [Apoh - apolipoprotein h, Plg - plasminogen, Serpinc1 - serine (or cysteine) peptidase inhibitor, clade c (antithrombin), member 1, Fga - fibrinogen alpha chain, Fgg - fibrinogen gamma chain, F2 - coagulation factor ii, Fcer1g - fc receptor, ige, high affinity i, gamma polypeptide, Hrg - histidine-rich glycoprotein, Kng1 - kininogen 1, Fgb - fibrinogen beta chain, Anxa2 - annexin a2] |
| GO:0001775 | cell activation | 4.53E-9 | 1.61E-6 | 4,71 | 4935 | 131 | 160 | 20 | [Igkc - immunoglobulin kappa constant, Anxa3 - annexin a3, Lgals1 - lectin, galactose binding, soluble 1, Itgam - integrin alpha m, Fn1 - fibronectin 1, Aif1 - allograft inflammatory factor 1, F2 - coagulation factor ii, Lcp1 - lymphocyte cytosolic protein 1, C1qa - complement component 1, q subcomponent, alpha polypeptide, Itgb3 - integrin beta 3, Itgb2 - integrin beta 2, Gba - glucosidase, beta, acid, Sema4a - sema domain, immunoglobulin domain (ig), transmembrane domain (tm) and short cytoplasmic domain, (semaphorin) 4a, Fga - fibrinogen alpha chain, Fgg - fibrinogen gamma chain, Fcer1g - fc receptor, ige, high affinity i, gamma polypeptide, Clu - clusterin, Ncstn - nicastrin, Cd151 - cd151 antigen, Fgb - fibrinogen beta chain] |
| GO:0050896 | response to stimulus | 6.55E-9 | 2.26E-6 | 1,76 | 4935 | 1364 | 160 | 78 | [Serpina1b - serine (or cysteine) preptidase inhibitor, clade a, member 1b, Wfs1 - wolfram syndrome 1 homolog (human), Fn1 - fibronectin 1, Serpina1a - serine (or cysteine) peptidase inhibitor, clade a, member 1a, Ahsg - alpha-2-hs-glycoprotein, Aif1 - allograft inflammatory factor 1, Gsn - gelsolin, Serpina1d - serine (or cysteine) peptidase inhibitor, clade a, member 1d, Serpina1e - serine (or cysteine) peptidase inhibitor, clade a, member 1e, Apoh - apolipoprotein h, Apod - apolipoprotein d, P4hb - prolyl 4-hydroxylase, beta polypeptide, Serpina3k - serine (or cysteine) peptidase inhibitor, clade a, member 3k, Ppp1r1b - protein phosphatase 1, regulatory (inhibitor) subunit 1b, Fcer1g - fc receptor, ige, high affinity i, gamma polypeptide, Alb - albumin, Kng1 - kininogen 1, Igkc - immunoglobulin kappa constant, Itgb1 - integrin beta 1 (fibronectin receptor beta), Vim - vimentin, Itgam - integrin alpha m, Cp - ceruloplasmin, F2 - coagulation factor ii, Fabp7 - fatty acid binding protein 7, brain, Itga6 - integrin alpha 6, Mgst1 - microsomal glutathione s-transferase 1, Vtn - vitronectin, Itgb3 - integrin beta 3, Tpm1 - tropomyosin 1, alpha, Gba - glucosidase, beta, acid, Gss - glutathione synthetase, Rbp1 - retinol binding protein 1, cellular, Itgb2 - integrin beta 2, Orm1 - orosomucoid 1, Serpinb1a - serine (or cysteine) peptidase inhibitor, clade b, member 1a, Ighg2b - immunoglobulin heavy constant gamma 2b, Ighg1 - immunoglobulin heavy constant gamma 1 (g1m marker), Ighm - immunoglobulin heavy constant mu, Irs2 - insulin receptor substrate 2, Tspan2 - tetraspanin 2, Stat3 - signal transducer and activator of transcription 3, Stat1 - signal transducer and activator of transcription 1, Anxa2 - annexin a2, Anxa3 - annexin a3, S100a16 - s100 calcium binding protein a16, Lcp1 - lymphocyte cytosolic protein 1, Cst3 - cystatin c, Lamp2 - lysosomal-associated membrane protein 2, Lgmn - legumain, Irgm1 - immunity-related gtpase family m member 1, Serpinc1 - serine (or cysteine) peptidase inhibitor, clade c (antithrombin), member 1, Fgg - fibrinogen gamma chain, Cfh - complement component factor h, Sod1 - superoxide dismutase 1, soluble, Fgb - fibrinogen beta chain, Ide - insulin degrading enzyme, C3 - complement component 3, Krt1 - keratin 1, H2-D1 - histocompatibility 2, d region locus 1, C4b - complement component 4b (chido blood group), Crot - carnitine o-octanoyltransferase, Cfb - complement factor b, Hp - haptoglobin, C1qc - complement component 1, q subcomponent, c chain, C1qa - complement component 1, q subcomponent, alpha polypeptide, Hrg - histidine-rich glycoprotein, C1qb - complement component 1, q subcomponent, beta polypeptide, Ddrgk1 - ddrgk domain containing 1, Ybx1 - y box protein 1, Sema4a - sema domain, immunoglobulin domain (ig), transmembrane domain (tm) and short cytoplasmic domain, (semaphorin) 4a, Gfap - glial fibrillary acidic protein, Fga - fibrinogen alpha chain, Pdcd5 - programmed cell death 5, Ncstn - nicastrin, Clu - clusterin, Commd1 - comm domain containing 1, Tbl2 - transducin (beta)-like 2, Ces1c - carboxylesterase 1c] |
| GO:0016064 | immunoglobulin mediated immune response | 2.81E-8 | 9.4E-6 | 23,13 | 4935 | 8 | 160 | 6 | [C4b - complement component 4b (chido blood group), Ighg2b - immunoglobulin heavy constant gamma 2b, Ighg1 - immunoglobulin heavy constant gamma 1 (g1m marker), Ighm - immunoglobulin heavy constant mu, Fcer1g - fc receptor, ige, high affinity i, gamma polypeptide, Cfh - complement component factor h] |
| GO:0019724 | B cell mediated immunity | 2.81E-8 | 9.14E-6 | 23,13 | 4935 | 8 | 160 | 6 | [C4b - complement component 4b (chido blood group), Ighg2b - immunoglobulin heavy constant gamma 2b, Ighg1 - immunoglobulin heavy constant gamma 1 (g1m marker), Fcer1g - fc receptor, ige, high affinity i, gamma polypeptide, Ighm - immunoglobulin heavy constant mu, Cfh - complement component factor h] |
| GO:0009617 | response to bacterium | 3.71E-8 | 1.17E-5 | 6,68 | 4935 | 60 | 160 | 13 | [Igkc - immunoglobulin kappa constant, C3 - complement component 3, Anxa3 - annexin a3, Hp - haptoglobin, Ighg2b - immunoglobulin heavy constant gamma 2b, Irgm1 - immunity-related gtpase family m member 1, Ighg1 - immunoglobulin heavy constant gamma 1 (g1m marker), Fga - fibrinogen alpha chain, Fcer1g - fc receptor, ige, high affinity i, gamma polypeptide, Ighm - immunoglobulin heavy constant mu, Stat1 - signal transducer and activator of transcription 1, Fgb - fibrinogen beta chain, Ces1c - carboxylesterase 1c] |

| | | | | | | | | | |
|---|---|---|---|---|---|---|---|---|---|
| GO:0006911 | phagocytosis, engulfment | 4.14E-8 | 1.27E-5 | 11,10 | 4935 | 25 | 160 | 9 | [Igkc - immunoglobulin kappa constant, Itgam - integrin alpha m, Ighg2b - immunoglobulin heavy constant gamma 2b, Ighg1 - immunoglobulin heavy constant gamma 1 (g1m marker), Aif1 - allograft inflammatory factor 1, Ighm - immunoglobulin heavy constant mu, Fcer1g - fc receptor, ige, high affinity i, gamma polypeptide, Gsn - gelsolin, Itgb2 - integrin beta 2] |
| GO:0045087 | innate immune response | 4.24E-8 | 1.27E-5 | 5,19 | 4935 | 95 | 160 | 16 | [Igkc - immunoglobulin kappa constant, C3 - complement component 3, Krt1 - keratin 1, Cfb - complement factor b, C1qc - complement component 1, q subcomponent, c chain, C1qa - complement component 1, q subcomponent, alpha polypeptide, C1qb - complement component 1, q subcomponent, beta polypeptide, Ighg2b - immunoglobulin heavy constant gamma 2b, Irgm1 - immunity-related gtpase family m member 1, Fga - fibrinogen alpha chain, Ighg1 - immunoglobulin heavy constant gamma 1 (g1m marker), Fgg - fibrinogen gamma chain, Fcer1g - fc receptor, ige, high affinity i, gamma polypeptide, Ighm - immunoglobulin heavy constant mu, Cfh - complement component factor h, Fgb - fibrinogen beta chain] |
| GO:0050878 | regulation of body fluid levels | 4.3E-8 | 1.25E-5 | 5,57 | 4935 | 83 | 160 | 15 | [Wfs1 - wolfram syndrome 1 homolog (human), C3 - complement component 3, Krt1 - keratin 1, Plg - plasminogen, F2 - coagulation factor ii, Hrg - histidine-rich glycoprotein, Gba - glucosidase, beta, acid, Apoh - apolipoprotein h, Serpinc1 - serine (or cysteine) peptidase inhibitor, clade c (antithrombin), member 1, Fga - fibrinogen alpha chain, Fgg - fibrinogen gamma chain, Fcer1g - fc receptor, ige, high affinity i, gamma polypeptide, Kng1 - kininogen 1, Fgb - fibrinogen beta chain, Anxa2 - annexin a2] |
| GO:0008284 | positive regulation of cell proliferation | 6.03E-8 | 1.71E-5 | 3,35 | 4935 | 230 | 160 | 25 | [Scg2 - secretogranin ii, Fn1 - fibronectin 1, Mvd - mevalonate (diphospho) decarboxylase, Aif1 - allograft inflammatory factor 1, Cst3 - cystatin c, Flna - filamin, alpha, Lgmn - legumain, Crip2 - cysteine rich protein 2, Marcksl1 - marcks-like 1, Itgb1 - integrin beta 1 (fibronectin receptor beta), Vim - vimentin, F2 - coagulation factor ii, S100a13 - s100 calcium binding protein a13, Ddrgk1 - ddrgk domain containing 1, Itgb3 - integrin beta 3, Ybx1 - y box protein 1, Tgm1 - transglutaminase 1, k polypeptide, Tgm2 - transglutaminase 2, c polypeptide, Gfap - glial fibrillary acidic protein, Ighm - immunoglobulin heavy constant mu, Irs2 - insulin receptor substrate 2, Clu - clusterin, Stat3 - signal transducer and activator of transcription 3, Stat1 - signal transducer and activator of transcription 1, Anxa2 - annexin a2] |
| GO:0080134 | regulation of response to stress | 6.74E-8 | 1.87E-5 | 2,56 | 4935 | 433 | 160 | 36 | [Wfs1 - wolfram syndrome 1 homolog (human), Plg - plasminogen, Ahsg - alpha-2-hs-glycoprotein, Lamp1 - lysosomal-associated membrane protein 1, Flna - filamin, alpha, Apoh - apolipoprotein h, Apod - apolipoprotein d, Serpinc1 - serine (or cysteine) peptidase inhibitor, clade c (antithrombin), member 1, Irgm1 - immunity-related gtpase family m member 1, P4hb - prolyl 4-hydroxylase, beta polypeptide, Fgg - fibrinogen gamma chain, Fcer1g - fc receptor, ige, high affinity i, gamma polypeptide, Cfh - complement component factor h, Sod1 - superoxide dismutase 1, soluble, Fgb - fibrinogen beta chain, Kng1 - kininogen 1, Krt1 - keratin 1, C3 - complement component 3, Itgb1 - integrin beta 1 (fibronectin receptor beta), Itgam - integrin alpha m, F2 - coagulation factor ii, Hrg - histidine-rich glycoprotein, Ddrgk1 - ddrgk domain containing 1, Ybx1 - y box protein 1, Hpx - hemopexin, Tgm2 - transglutaminase 2, c polypeptide, Ighg2b - immunoglobulin heavy constant gamma 2b, Serpinb1a - serine (or cysteine) peptidase inhibitor, clade b, member 1a, Ighg1 - immunoglobulin heavy constant gamma 1 (g1m marker), Fga - fibrinogen alpha chain, Clu - clusterin, Apoa1 - apolipoprotein a-i, Commd1 - comm domain containing 1, Stat3 - signal transducer and activator of transcription 3, Stat1 - signal transducer and activator of transcription 1, Anxa2 - annexin a2] |
| GO:0010951 | negative regulation of endopeptidase activity | 6.9E-8 | 1.87E-5 | 6,36 | 4935 | 63 | 160 | 13 | [Serpina1b - serine (or cysteine) preptidase inhibitor, clade a, member 1b, Serpina6 - serine (or cysteine) peptidase inhibitor, clade a, member 6, Serpina1a - serine (or cysteine) peptidase inhibitor, clade a, member 1a, Ahsg - alpha-2-hs-glycoprotein, Serpinh1 - serine (or cysteine) peptidase inhibitor, clade h, member 1, Hrg - histidine-rich glycoprotein, Vtn - vitronectin, Serpina1d - serine (or cysteine) peptidase inhibitor, clade a, member 1d, Serpina1e - serine (or cysteine) peptidase inhibitor, clade a, member 1e, Serpinb1a - serine (or cysteine) peptidase inhibitor, clade b, member 1a, Serpinc1 - serine (or cysteine) peptidase inhibitor, clade c (antithrombin), member 1, Serpina3k - serine (or cysteine) peptidase inhibitor, clade a, member 3k, Kng1 - kininogen 1] |
| GO:0061045 | negative regulation of wound healing | 8.99E-8 | 2.38E-5 | 10,28 | 4935 | 27 | 160 | 9 | [Apoh - apolipoprotein h, Plg - plasminogen, Fga - fibrinogen alpha chain, Fgg - fibrinogen gamma chain, F2 - coagulation factor ii, Hrg - histidine-rich glycoprotein, Fgb - fibrinogen beta chain, Kng1 - kininogen 1, Anxa2 - annexin a2] |
| GO:0002250 | adaptive immune response | 1.16E-7 | 2.99E-5 | 7,54 | 4935 | 45 | 160 | 11 | [Sema4a - sema domain, immunoglobulin domain (ig), transmembrane domain (tm) and short cytoplasmic domain, (semaphorin) 4a, C4b - complement component 4b (chido blood group), Ighg2b - immunoglobulin heavy constant gamma 2b, Ighg1 - immunoglobulin heavy constant gamma 1 (g1m marker), Fga - fibrinogen alpha chain, Fgg - fibrinogen gamma chain, Ighm - immunoglobulin heavy constant mu, Fcer1g - fc receptor, ige, high affinity i, gamma polypeptide, Cfh - complement component factor h, Stat3 - signal transducer and activator of transcription 3, Fgb - fibrinogen beta chain] |
| GO:0048584 | positive regulation of response to stimulus | 1.23E-7 | 3.11E-5 | 2,22 | 4935 | 611 | 160 | 44 | [Plg - plasminogen, Scg2 - secretogranin ii, Fn1 - fibronectin 1, Aif1 - allograft inflammatory factor 1, Gsn - gelsolin, Lamp1 - lysosomal-associated membrane protein 1, Flna - filamin, alpha, Apoh - apolipoprotein h, Lgmn - legumain, Irgm1 - immunity-related gtpase family m member 1, Fgg - fibrinogen gamma chain, Fcer1g - fc receptor, ige, high affinity i, gamma polypeptide, Cfh - complement component factor h, Sod1 - superoxide dismutase 1, soluble, Fgb - fibrinogen beta chain, Krt1 - keratin 1, Igkc - immunoglobulin kappa constant, Glipr2 - gli pathogenesis-related 2, C3 - complement component 3, Itgb1 - integrin beta 1 (fibronectin receptor beta), H2-D1 - histocompatibility 2, d region locus 1, Itgam - integrin alpha m, C4b - complement component 4b (chido blood group), Cfb - complement factor b, C1qc - complement component 1, q subcomponent, c chain, F2 - coagulation factor ii, Fabp5 - fatty acid binding protein 5, epidermal, S100a13 - s100 calcium binding protein a13, C1qa - complement component 1, q subcomponent, alpha polypeptide, Hrg - histidine-rich glycoprotein, C1qb - complement component 1, q subcomponent, beta polypeptide, Ddrgk1 - ddrgk domain containing 1, Itgb3 - integrin beta 3, Itgb2 - integrin beta 2, Hpx - hemopexin, Tgm2 - transglutaminase 2, c polypeptide, Ighg2b - immunoglobulin heavy constant gamma 2b, Fga - fibrinogen alpha chain, Ighg1 - immunoglobulin heavy constant gamma 1 (g1m marker), Ighm - immunoglobulin heavy constant mu, Clu - clusterin, Pdcd5 - programmed cell death 5, Apoa1 - apolipoprotein a-i, Stat3 - signal transducer and activator of transcription 3] |

| | | | | | | | | | |
|---|---|---|---|---|---|---|---|---|---|
| GO:0032102 | negative regulation of response to external stimulus | 1.58E-7 | 3.91E-5 | 4,75 | 4935 | 104 | 160 | 16 | [Krt1 - keratin 1, Plg - plasminogen, Aif1 - allograft inflammatory factor 1, F2 - coagulation factor ii, Fabp7 - fatty acid binding protein 7, brain, Hrg - histidine-rich glycoprotein, Sema4a - sema domain, immunoglobulin domain (ig), transmembrane domain (tm) and short cytoplasmic domain, (semaphorin) 4a, Apoh - apolipoprotein h, Apod - apolipoprotein d, Fga - fibrinogen alpha chain, Fgg - fibrinogen gamma chain, Apoa1 - apolipoprotein a-i, Sod1 - superoxide dismutase 1, soluble, Kng1 - kininogen 1, Fgb - fibrinogen beta chain, Anxa2 - annexin a2] |
| GO:0065009 | regulation of molecular function | 1.62E-7 | 3.92E-5 | 1,92 | 4935 | 917 | 160 | 57 | [Pzp - pregnancy zone protein, Wfs1 - wolfram syndrome 1 homolog (human), Serpina1b - serine (or cysteine) preptidase inhibitor, clade a, member 1b, Ahsg - alpha-2-hs-glycoprotein, Serpina1a - serine (or cysteine) peptidase inhibitor, clade a, member 1a, Gsn - gelsolin, Serpina1d - serine (or cysteine) peptidase inhibitor, clade a, member 1d, Serpina1e - serine (or cysteine) peptidase inhibitor, clade a, member 1e, Flna - filamin, alpha, Apoh - apolipoprotein h, Ppp1r1b - protein phosphatase 1, regulatory (inhibitor) subunit 1b, Serpina3k - serine (or cysteine) peptidase inhibitor, clade a, member 3k, Kng1 - kininogen 1, Itgb1 - integrin beta 1 (fibronectin receptor beta), Itgam - integrin alpha m, Itga6 - integrin alpha 6, F2 - coagulation factor ii, Arhgap22 - rho gtpase activating protein 22, Psme1 - proteasome (prosome, macropain) activator subunit 1 (pa28 alpha), Vtn - vitronectin, Itgb3 - integrin beta 3, Tpm1 - tropomyosin 1, alpha, Gba - glucosidase, beta, acid, Itgb2 - integrin beta 2, Scarb2 - scavenger receptor class b, member 2, Serpinb1a - serine (or cysteine) peptidase inhibitor, clade b, member 1a, Itih3 - inter-alpha trypsin inhibitor, heavy chain 3, Itih2 - inter-alpha trypsin inhibitor, heavy chain 2, Ighm - immunoglobulin heavy constant mu, Itih1 - inter-alpha trypsin inhibitor, heavy chain 1, Irs2 - insulin receptor substrate 2, Nptxr - neuronal pentraxin receptor, Stat3 - signal transducer and activator of transcription 3, Stat1 - signal transducer and activator of transcription 1, Anxa2 - annexin a2, Scn3b - sodium channel, voltage-gated, type iii, beta, Serpina6 - serine (or cysteine) peptidase inhibitor, clade a, member 6, Anxa3 - annexin a3, Serpinh1 - serine (or cysteine) peptidase inhibitor, clade h, member 1, Ctsb - cathepsin b, Cst3 - cystatin c, Lgmn - legumain, Serpinc1 - serine (or cysteine) peptidase inhibitor, clade c (antithrombin), member 1, Ctsd - cathepsin d, Pex19 - peroxisomal biogenesis factor 19, Sod1 - superoxide dismutase 1, soluble, Ide - insulin degrading enzyme, Hp - haptoglobin, Hrg - histidine-rich glycoprotein, Ddrgk1 - ddrgk domain containing 1, Pdcd5 - programmed cell death 5, Apoa1 - apolipoprotein a-i, Clu - clusterin, Ncstn - nicastrin, Nptx2 - neuronal pentraxin 2, Commd1 - comm domain containing 1, Mug1 - murinoglobulin 1] |
| GO:0050776 | regulation of immune response | 1.76E-7 | 4.18E-5 | 3,52 | 4935 | 193 | 160 | 22 | [C3 - complement component 3, Igkc - immunoglobulin kappa constant, Krt1 - keratin 1, H2-D1 - histocompatibility 2, d region locus 1, C4b - complement component 4b (chido blood group), Itgam - integrin alpha m, Cfb - complement factor b, C1qc - complement component 1, q subcomponent, c chain, C1qa - complement component 1, q subcomponent, alpha polypeptide, C1qb - complement component 1, q subcomponent, beta polypeptide, Hrg - histidine-rich glycoprotein, Lamp1 - lysosomal-associated membrane protein 1, Itgb2 - integrin beta 2, Hpx - hemopexin, Ighg2b - immunoglobulin heavy constant gamma 2b, Serpinb1a - serine (or cysteine) peptidase inhibitor, clade b, member 1a, Irgm1 - immunity-related gtpase family m member 1, Ighg1 - immunoglobulin heavy constant gamma 1 (g1m marker), Fcer1g - fc receptor, ige, high affinity i, gamma polypeptide, Ighm - immunoglobulin heavy constant mu, Cfh - complement component factor h, Apoa1 - apolipoprotein a-i] |
| GO:0061041 | regulation of wound healing | 1.79E-7 | 4.15E-5 | 6,49 | 4935 | 57 | 160 | 12 | [Plg - plasminogen, Itgb1 - integrin beta 1 (fibronectin receptor beta), Apoh - apolipoprotein h, Serpinc1 - serine (or cysteine) peptidase inhibitor, clade c (antithrombin), member 1, Fga - fibrinogen alpha chain, Fgg - fibrinogen gamma chain, F2 - coagulation factor ii, Fcer1g - fc receptor, ige, high affinity i, gamma polypeptide, Hrg - histidine-rich glycoprotein, Fgb - fibrinogen beta chain, Kng1 - kininogen 1, Anxa2 - annexin a2] |
| GO:0051246 | regulation of protein metabolic process | 2.12E-7 | 4.83E-5 | 1,90 | 4935 | 924 | 160 | 57 | [Pzp - pregnancy zone protein, Wfs1 - wolfram syndrome 1 homolog (human), Serpina1b - serine (or cysteine) preptidase inhibitor, clade a, member 1b, Ahsg - alpha-2-hs-glycoprotein, Serpina1a - serine (or cysteine) peptidase inhibitor, clade a, member 1a, Fn1 - fibronectin 1, Aif1 - allograft inflammatory factor 1, Gsn - gelsolin, Serpina1d - serine (or cysteine) peptidase inhibitor, clade a, member 1d, Serpina1e - serine (or cysteine) peptidase inhibitor, clade a, member 1e, Flna - filamin, alpha, Apod - apolipoprotein d, Ppp1r1b - protein phosphatase 1, regulatory (inhibitor) subunit 1b, Serpina3k - serine (or cysteine) peptidase inhibitor, clade a, member 3k, Ctsa - cathepsin a, Kng1 - kininogen 1, Glipr2 - gli pathogenesis-related 2, Itgb1 - integrin beta 1 (fibronectin receptor beta), Vim - vimentin, F2 - coagulation factor ii, Psme1 - proteasome (prosome, macropain) activator subunit 1 (pa28 alpha), Vtn - vitronectin, Itgb3 - integrin beta 3, Gba - glucosidase, beta, acid, Itgb2 - integrin beta 2, Serpinb1a - serine (or cysteine) peptidase inhibitor, clade b, member 1a, Itih3 - inter-alpha trypsin inhibitor, heavy chain 3, Itih2 - inter-alpha trypsin inhibitor, heavy chain 2, Ighm - immunoglobulin heavy constant mu, Itih1 - inter-alpha trypsin inhibitor, heavy chain 1, Stat3 - signal transducer and activator of transcription 3, Stat1 - signal transducer and activator of transcription 1, Anxa2 - annexin a2, Serpina6 - serine (or cysteine) peptidase inhibitor, clade a, member 6, Serpinh1 - serine (or cysteine) peptidase inhibitor, clade h, member 1, Cst3 - cystatin c, Lgmn - legumain, Serpinc1 - serine (or cysteine) peptidase inhibitor, clade c (antithrombin), member 1, Fgg - fibrinogen gamma chain, Fhit - fragile histidine triad gene, Cfh - complement component factor h, Ctsd - cathepsin d, Sod1 - superoxide dismutase 1, soluble, Fgb - fibrinogen beta chain, Ide - insulin degrading enzyme, C3 - complement component 3, Hrg - histidine-rich glycoprotein, Ddrgk1 - ddrgk domain containing 1, Hpx - hemopexin, Gfap - glial fibrillary acidic protein, Fga - fibrinogen alpha chain, Pdcd5 - programmed cell death 5, Apoa1 - apolipoprotein a-i, Clu - clusterin, Ncstn - nicastrin, Commd1 - comm domain containing 1, Mug1 - murinoglobulin 1] |
| GO:0099024 | plasma membrane invagination | 2.52E-7 | 5.63E-5 | 9,25 | 4935 | 30 | 160 | 9 | [Igkc - immunoglobulin kappa constant, Itgam - integrin alpha m, Ighg2b - immunoglobulin heavy constant gamma 2b, Ighg1 - immunoglobulin heavy constant gamma 1 (g1m marker), Aif1 - allograft inflammatory factor 1, Ighm - immunoglobulin heavy constant mu, Fcer1g - fc receptor, ige, high affinity i, gamma polypeptide, Gsn - gelsolin, Itgb2 - integrin beta 2] |

| | | | | | | | | | |
|---|---|---|---|---|---|---|---|---|---|
| GO:0050790 | regulation of catalytic activity | 3.05E-7 | 6.68E-5 | 2,10 | 4935 | 675 | 160 | 46 | [Pzp - pregnancy zone protein, Serpina1b - serine (or cysteine) preptidase inhibitor, clade a, member 1b, Serpina6 - serine (or cysteine) peptidase inhibitor, clade a, member 6, Ahsg - alpha-2-hs-glycoprotein, Serpina1a - serine (or cysteine) peptidase inhibitor, clade a, member 1a, Serpinh1 - serine (or cysteine) peptidase inhibitor, clade h, member 1, Ctsb - cathepsin b, Gsn - gelsolin, Cst3 - cystatin c, Serpina1d - serine (or cysteine) peptidase inhibitor, clade a, member 1d, Serpina1e - serine (or cysteine) peptidase inhibitor, clade a, member 1e, Apoh - apolipoprotein h, Serpinc1 - serine (or cysteine) peptidase inhibitor, clade c (antithrombin), member 1, Lgmn - legumain, Serpina3k - serine (or cysteine) peptidase inhibitor, clade a, member 3k, Ppp1r1b - protein phosphatase 1, regulatory (inhibitor) subunit 1b, Ctsd - cathepsin d, Sod1 - superoxide dismutase 1, soluble, Kng1 - kininogen 1, Itgb1 - integrin beta 1 (fibronectin receptor beta), Itgam - integrin alpha m, Hp - haptoglobin, Itga6 - integrin alpha 6, F2 - coagulation factor ii, Arhgap22 - rho gtpase activating protein 22, Hrg - histidine-rich glycoprotein, Psme1 - proteasome (prosome, macropain) activator subunit 1 (pa28 alpha), Ddrgk1 - ddrgk domain containing 1, Vtn - vitronectin, Itgb3 - integrin beta 3, Tpm1 - tropomyosin 1, alpha, Gba - glucosidase, beta, acid, Scarb2 - scavenger receptor class b, member 2, Itih3 - inter-alpha trypsin inhibitor, heavy chain 3, Serpinb1a - serine (or cysteine) peptidase inhibitor, clade b, member 1a, Itih2 - inter-alpha trypsin inhibitor, heavy chain 2, Itih1 - inter-alpha trypsin inhibitor, heavy chain 1, Ighm - immunoglobulin heavy constant mu, Clu - clusterin, Pdcd5 - programmed cell death 5, Irs2 - insulin receptor substrate 2, Apoa1 - apolipoprotein a-i, Ncstn - nicastrin, Mug1 - murinoglobulin 1, Stat3 - signal transducer and activator of transcription 3, Stat1 - signal transducer and activator of transcription 1] |
| GO:0002253 | activation of immune response | 3.63E-7 | 7.79E-5 | 5,57 | 4935 | 72 | 160 | 13 | [Igkc - immunoglobulin kappa constant, Krt1 - keratin 1, C3 - complement component 3, C4b - complement component 4b (chido blood group), Cfb - complement factor b, C1qc - complement component 1, q subcomponent, c chain, C1qa - complement component 1, q subcomponent, alpha polypeptide, C1qb - complement component 1, q subcomponent, beta polypeptide, Ighg2b - immunoglobulin heavy constant gamma 2b, Ighg1 - immunoglobulin heavy constant gamma 1 (g1m marker), Fcer1g - fc receptor, ige, high affinity i, gamma polypeptide, Ighm - immunoglobulin heavy constant mu, Cfh - complement component factor h] |
| GO:0002705 | positive regulation of leukocyte mediated immunity | 4.68E-7 | 9.86E-5 | 8,67 | 4935 | 32 | 160 | 9 | [Hpx - hemopexin, C3 - complement component 3, H2-D1 - histocompatibility 2, d region locus 1, Itgam - integrin alpha m, Ighg2b - immunoglobulin heavy constant gamma 2b, Ighg1 - immunoglobulin heavy constant gamma 1 (g1m marker), Fcer1g - fc receptor, ige, high affinity i, gamma polypeptide, Lamp1 - lysosomal-associated membrane protein 1, Itgb2 - integrin beta 2] |
| GO:0006954 | inflammatory response | 5.95E-7 | 1.23E-4 | 5,35 | 4935 | 75 | 160 | 13 | [Serpina1b - serine (or cysteine) preptidase inhibitor, clade a, member 1b, C3 - complement component 3, Fn1 - fibronectin 1, Ahsg - alpha-2-hs-glycoprotein, Hp - haptoglobin, F2 - coagulation factor ii, Orm1 - orosomucoid 1, Serpinb1a - serine (or cysteine) peptidase inhibitor, clade b, member 1a, Ighg1 - immunoglobulin heavy constant gamma 1 (g1m marker), Cfh - complement component factor h, Tspan2 - tetraspanin 2, Stat3 - signal transducer and activator of transcription 3, Kng1 - kininogen 1] |
| GO:0002714 | positive regulation of B cell mediated immunity | 6.71E-7 | 1.36E-4 | 22,03 | 4935 | 7 | 160 | 5 | [C3 - complement component 3, Hpx - hemopexin, Ighg2b - immunoglobulin heavy constant gamma 2b, Ighg1 - immunoglobulin heavy constant gamma 1 (g1m marker), Fcer1g - fc receptor, ige, high affinity i, gamma polypeptide] |
| GO:0002891 | positive regulation of immunoglobulin mediated immune response | 6.71E-7 | 1.34E-4 | 22,03 | 4935 | 7 | 160 | 5 | [Hpx - hemopexin, C3 - complement component 3, Ighg2b - immunoglobulin heavy constant gamma 2b, Ighg1 - immunoglobulin heavy constant gamma 1 (g1m marker), Fcer1g - fc receptor, ige, high affinity i, gamma polypeptide] |
| GO:0010324 | membrane invagination | 8.27E-7 | 1.62E-4 | 8,16 | 4935 | 34 | 160 | 9 | [Igkc - immunoglobulin kappa constant, Itgam - integrin alpha m, Ighg2b - immunoglobulin heavy constant gamma 2b, Ighg1 - immunoglobulin heavy constant gamma 1 (g1m marker), Aif1 - allograft inflammatory factor 1, Ighm - immunoglobulin heavy constant mu, Fcer1g - fc receptor, ige, high affinity i, gamma polypeptide, Gsn - gelsolin, Itgb2 - integrin beta 2] |
| GO:0044092 | negative regulation of molecular function | 1,00E-06 | 1.93E-4 | 2,57 | 4935 | 360 | 160 | 30 | [Wfs1 - wolfram syndrome 1 homolog (human), Pzp - pregnancy zone protein, Serpina1b - serine (or cysteine) preptidase inhibitor, clade a, member 1b, Serpina6 - serine (or cysteine) peptidase inhibitor, clade a, member 6, Ahsg - alpha-2-hs-glycoprotein, Serpina1a - serine (or cysteine) peptidase inhibitor, clade a, member 1a, Serpinh1 - serine (or cysteine) peptidase inhibitor, clade h, member 1, Cst3 - cystatin c, Serpina1d - serine (or cysteine) peptidase inhibitor, clade a, member 1d, Serpina1e - serine (or cysteine) peptidase inhibitor, clade a, member 1e, Flna - filamin, alpha, Serpinc1 - serine (or cysteine) peptidase inhibitor, clade c (antithrombin), member 1, Serpina3k - serine (or cysteine) peptidase inhibitor, clade a, member 3k, Ppp1r1b - protein phosphatase 1, regulatory (inhibitor) subunit 1b, Pex19 - peroxisomal biogenesis factor 19, Kng1 - kininogen 1, Hp - haptoglobin, Hrg - histidine-rich glycoprotein, Vtn - vitronectin, Itgb3 - integrin beta 3, Gba - glucosidase, beta, acid, Itih3 - inter-alpha trypsin inhibitor, heavy chain 3, Serpinb1a - serine (or cysteine) peptidase inhibitor, clade b, member 1a, Itih2 - inter-alpha trypsin inhibitor, heavy chain 2, Itih1 - inter-alpha trypsin inhibitor, heavy chain 1, Apoa1 - apolipoprotein a-i, Irs2 - insulin receptor substrate 2, Commd1 - comm domain containing 1, Mug1 - murinoglobulin 1, Anxa2 - annexin a2] |
| GO:0001796 | regulation of type IIa hypersensitivity | 1.07E-6 | 2.02E-4 | 30,84 | 4935 | 4 | 160 | 4 | [C3 - complement component 3, Ighg2b - immunoglobulin heavy constant gamma 2b, Ighg1 - immunoglobulin heavy constant gamma 1 (g1m marker), Fcer1g - fc receptor, ige, high affinity i, gamma polypeptide] |

| | | | | | | | | | |
|---|---|---|---|---|---|---|---|---|---|
| GO:0001798 | positive regulation of type IIa hypersensitivity | 1.07E-6 | 1.99E-4 | 30,84 | 4935 | 4 | 160 | 4 | [C3 - complement component 3, Ighg2b - immunoglobulin heavy constant gamma 2b, Ighg1 - immunoglobulin heavy constant gamma 1 (g1m marker), Fcer1g - fc receptor, ige, high affinity i, gamma polypeptide] |
| GO:0002885 | positive regulation of hypersensitivity | 1.07E-6 | 1.96E-4 | 30,84 | 4935 | 4 | 160 | 4 | [C3 - complement component 3, Ighg2b - immunoglobulin heavy constant gamma 2b, Ighg1 - immunoglobulin heavy constant gamma 1 (g1m marker), Fcer1g - fc receptor, ige, high affinity i, gamma polypeptide] |
| GO:0002892 | regulation of type II hypersensitivity | 1.07E-6 | 1.92E-4 | 30,84 | 4935 | 4 | 160 | 4 | [C3 - complement component 3, Ighg2b - immunoglobulin heavy constant gamma 2b, Ighg1 - immunoglobulin heavy constant gamma 1 (g1m marker), Fcer1g - fc receptor, ige, high affinity i, gamma polypeptide] |
| GO:0002894 | positive regulation of type II hypersensitivity | 1.07E-6 | 1.89E-4 | 30,84 | 4935 | 4 | 160 | 4 | [C3 - complement component 3, Ighg2b - immunoglobulin heavy constant gamma 2b, Ighg1 - immunoglobulin heavy constant gamma 1 (g1m marker), Fcer1g - fc receptor, ige, high affinity i, gamma polypeptide] |
| GO:0002883 | regulation of hypersensitivity | 1.07E-6 | 1.86E-4 | 30,84 | 4935 | 4 | 160 | 4 | [C3 - complement component 3, Ighg2b - immunoglobulin heavy constant gamma 2b, Ighg1 - immunoglobulin heavy constant gamma 1 (g1m marker), Fcer1g - fc receptor, ige, high affinity i, gamma polypeptide] |
| GO:0007160 | cell-matrix adhesion | 1.08E-6 | 1.87E-4 | 7,93 | 4935 | 35 | 160 | 9 | [Itgb1 - integrin beta 1 (fibronectin receptor beta), Fn1 - fibronectin 1, Fga - fibrinogen alpha chain, Fgg - fibrinogen gamma chain, Itga6 - integrin alpha 6, Vtn - vitronectin, Itgb3 - integrin beta 3, Fgb - fibrinogen beta chain, Itgb2 - integrin beta 2] |
| GO:0065008 | regulation of biological quality | 1.12E-6 | 1.91E-4 | 1,64 | 4935 | 1358 | 160 | 72 | [Wfs1 - wolfram syndrome 1 homolog (human), Fn1 - fibronectin 1, Ahsg - alpha-2-hs-glycoprotein, Clic1 - chloride intracellular channel 1, Aif1 - allograft inflammatory factor 1, Gsn - gelsolin, Stx12 - syntaxin 12, Flna - filamin, alpha, Apoh - apolipoprotein h, P4hb - prolyl 4-hydroxylase, beta polypeptide, Ppp1r1b - protein phosphatase 1, regulatory (inhibitor) subunit 1b, Fcer1g - fc receptor, ige, high affinity i, gamma polypeptide, Alb - albumin, Ctsa - cathepsin a, Cpe - carboxypeptidase e, Kng1 - kininogen 1, Itgb1 - integrin beta 1 (fibronectin receptor beta), Vim - vimentin, Cp - ceruloplasmin, F2 - coagulation factor ii, Fabp5 - fatty acid binding protein 5, epidermal, Itgb3 - integrin beta 3, Sh3bgrl3 - sh3 domain binding glutamic acid-rich protein-like 3, Tpm1 - tropomyosin 1, alpha, Gba - glucosidase, beta, acid, Itgb2 - integrin beta 2, Rbp1 - retinol binding protein 1, cellular, Serpinb1a - serine (or cysteine) peptidase inhibitor, clade b, member 1a, Ampd3 - adenosine monophosphate deaminase 3, Irs2 - insulin receptor substrate 2, Nptxr - neuronal pentraxin receptor, Stat3 - signal transducer and activator of transcription 3, Arpc1b - actin related protein 2/3 complex, subunit 1b, Stat1 - signal transducer and activator of transcription 1, Scn3b - sodium channel, voltage-gated, type iii, beta, Anxa2 - annexin a2, Plg - plasminogen, Serpina6 - serine (or cysteine) peptidase inhibitor, clade a, member 6, Ctsb - cathepsin b, Lamp2 - lysosomal-associated membrane protein 2, Lamp1 - lysosomal-associated membrane protein 1, Anpep - alanyl (membrane) aminopeptidase, Lgmn - legumain, Serpinc1 - serine (or cysteine) peptidase inhibitor, clade c (antithrombin), member 1, Fgg - fibrinogen gamma chain, Cfh - complement component factor h, Ttr - transthyretin, Pex19 - peroxisomal biogenesis factor 19, Gaa - glucosidase, alpha, acid, Sod1 - superoxide dismutase 1, soluble, Fgb - fibrinogen beta chain, Ide - insulin degrading enzyme, C3 - complement component 3, Krt1 - keratin 1, S100a13 - s100 calcium binding protein a13, Hrg - histidine-rich glycoprotein, Kcnj10 - potassium inwardly-rectifying channel, subfamily j, member 10, Ddrgk1 - ddrgk domain containing 1, Hexb - hexosaminidase b, Ybx1 - y box protein 1, Hpx - hemopexin, Tgm2 - transglutaminase 2, c polypeptide, Sema4a - sema domain, immunoglobulin domain (ig), transmembrane domain (tm) and short cytoplasmic domain, (semaphorin) 4a, Gfap - glial fibrillary acidic protein, Fga - fibrinogen alpha chain, Ncstn - nicastrin, Apoa1 - apolipoprotein a-i, Car14 - carbonic anhydrase 14, Clu - clusterin, Nptx2 - neuronal pentraxin 2, Txndc5 - thioredoxin domain containing 5, Commd1 - comm domain containing 1] |
| GO:0043086 | negative regulation of catalytic activity | 1.17E-6 | 1.96E-4 | 2,96 | 4935 | 250 | 160 | 24 | [Serpina1b - serine (or cysteine) preptidase inhibitor, clade a, member 1b, Pzp - pregnancy zone protein, Serpina6 - serine (or cysteine) peptidase inhibitor, clade a, member 6, Hp - haptoglobin, Serpina1a - serine (or cysteine) peptidase inhibitor, clade a, member 1a, Ahsg - alpha-2-hs-glycoprotein, Serpinh1 - serine (or cysteine) peptidase inhibitor, clade h, member 1, Cst3 - cystatin c, Hrg - histidine-rich glycoprotein, Vtn - vitronectin, Serpina1d - serine (or cysteine) peptidase inhibitor, clade a, member 1d, Serpina1e - serine (or cysteine) peptidase inhibitor, clade a, member 1e, Gba - glucosidase, beta, acid, Itih3 - inter-alpha trypsin inhibitor, heavy chain 3, Serpinb1a - serine (or cysteine) peptidase inhibitor, clade b, member 1a, Serpinc1 - serine (or cysteine) peptidase inhibitor, clade c (antithrombin), member 1, Itih2 - inter-alpha trypsin inhibitor, heavy chain 2, Serpina3k - serine (or cysteine) peptidase inhibitor, clade a, member 3k, Ppp1r1b - protein phosphatase 1, regulatory (inhibitor) subunit 1b, Itih1 - inter-alpha trypsin inhibitor, heavy chain 1, Apoa1 - apolipoprotein a-i, Irs2 - insulin receptor substrate 2, Mug1 - murinoglobulin 1, Kng1 - kininogen 1] |
| GO:1903034 | regulation of response to wounding | 1.29E-6 | 2.12E-4 | 5,01 | 4935 | 80 | 160 | 13 | [Itgb1 - integrin beta 1 (fibronectin receptor beta), Plg - plasminogen, F2 - coagulation factor ii, Hrg - histidine-rich glycoprotein, Flna - filamin, alpha, Apoh - apolipoprotein h, Serpinc1 - serine (or cysteine) peptidase inhibitor, clade c (antithrombin), member 1, Fgg - fibrinogen gamma chain, Fga - fibrinogen alpha chain, Fcer1g - fc receptor, ige, high affinity i, gamma polypeptide, Kng1 - kininogen 1, Fgb - fibrinogen beta chain, Anxa2 - annexin a2] |
| GO:0042742 | defense response to bacterium | 1.4E-6 | 2.28E-4 | 7,71 | 4935 | 36 | 160 | 9 | [Igkc - immunoglobulin kappa constant, Anxa3 - annexin a3, Hp - haptoglobin, Ighg2b - immunoglobulin heavy constant gamma 2b, Ighg1 - immunoglobulin heavy constant gamma 1 (g1m marker), Fga - fibrinogen alpha chain, Ighm - immunoglobulin heavy constant mu, Fcer1g - fc receptor, ige, high affinity i, gamma polypeptide, Fgb - fibrinogen beta chain] |

| | | | | | | | | | |
|---|---|---|---|---|---|---|---|---|---|
| GO:1903035 | negative regulation of response to wounding | 1.4E-6 | 2.25E-4 | 7,71 | 4935 | 36 | 160 | 9 | [Apoh - apolipoprotein h, Plg - plasminogen, Fga - fibrinogen alpha chain, Fgg - fibrinogen gamma chain, F2 - coagulation factor ii, Hrg - histidine-rich glycoprotein, Fgb - fibrinogen beta chain, Kng1 - kininogen 1, Anxa2 - annexin a2] |
| GO:0032103 | positive regulation of response to external stimulus | 1.48E-6 | 2.33E-4 | 4,59 | 4935 | 94 | 160 | 14 | [C3 - complement component 3, Scg2 - secretogranin ii, Plg - plasminogen, Fn1 - fibronectin 1, Aif1 - allograft inflammatory factor 1, Hrg - histidine-rich glycoprotein, Flna - filamin, alpha, Tgm2 - transglutaminase 2, c polypeptide, Apoh - apolipoprotein h, Ighg2b - immunoglobulin heavy constant gamma 2b, Lgmn - legumain, Ighg1 - immunoglobulin heavy constant gamma 1 (g1m marker), Fcer1g - fc receptor, ige, high affinity i, gamma polypeptide, Stat3 - signal transducer and activator of transcription 3] |
| GO:0051707 | response to other organism | 1.73E-6 | 2.69E-4 | 4,24 | 4935 | 109 | 160 | 15 | [Igkc - immunoglobulin kappa constant, C3 - complement component 3, Anxa3 - annexin a3, Hp - haptoglobin, F2 - coagulation factor ii, Hrg - histidine-rich glycoprotein, Irgm1 - immunity-related gtpase family m member 1, Ighg2b - immunoglobulin heavy constant gamma 2b, Fga - fibrinogen alpha chain, Ighg1 - immunoglobulin heavy constant gamma 1 (g1m marker), Fcer1g - fc receptor, ige, high affinity i, gamma polypeptide, Ighm - immunoglobulin heavy constant mu, Stat1 - signal transducer and activator of transcription 1, Fgb - fibrinogen beta chain, Ces1c - carboxylesterase 1c] |
| GO:0098883 | synapse pruning | 1.74E-6 | 2.68E-4 | 19,28 | 4935 | 8 | 160 | 5 | [C3 - complement component 3, Itgam - integrin alpha m, C1qc - complement component 1, q subcomponent, c chain, C1qa - complement component 1, q subcomponent, alpha polypeptide, C1qb - complement component 1, q subcomponent, beta polypeptide] |
| GO:0033627 | cell adhesion mediated by integrin | 1.74E-6 | 2.64E-4 | 19,28 | 4935 | 8 | 160 | 5 | [Itgb1 - integrin beta 1 (fibronectin receptor beta), Itga6 - integrin alpha 6, Vtn - vitronectin, Itgb3 - integrin beta 3, Itgb2 - integrin beta 2] |
| GO:0043207 | response to external biotic stimulus | 1.77E-6 | 2.65E-4 | 3,61 | 4935 | 154 | 160 | 18 | [C3 - complement component 3, Igkc - immunoglobulin kappa constant, Anxa3 - annexin a3, Vim - vimentin, Hp - haptoglobin, F2 - coagulation factor ii, Mgst1 - microsomal glutathione s-transferase 1, Hrg - histidine-rich glycoprotein, Irgm1 - immunity-related gtpase family m member 1, Ighg2b - immunoglobulin heavy constant gamma 2b, Fga - fibrinogen alpha chain, Ighg1 - immunoglobulin heavy constant gamma 1 (g1m marker), Fcer1g - fc receptor, ige, high affinity i, gamma polypeptide, Ighm - immunoglobulin heavy constant mu, Cfh - complement component factor h, Stat1 - signal transducer and activator of transcription 1, Fgb - fibrinogen beta chain, Ces1c - carboxylesterase 1c] |
| GO:0051241 | negative regulation of multicellular organismal process | 1.79E-6 | 2.64E-4 | 2,50 | 4935 | 370 | 160 | 30 | [Plg - plasminogen, Fn1 - fibronectin 1, Ahsg - alpha-2-hs-glycoprotein, Anpep - alanyl (membrane) aminopeptidase, Flna - filamin, alpha, Apoh - apolipoprotein h, Apod - apolipoprotein d, Lgmn - legumain, Fgg - fibrinogen gamma chain, Sod1 - superoxide dismutase 1, soluble, Cpe - carboxypeptidase e, Kng1 - kininogen 1, Fgb - fibrinogen beta chain, Itgb1 - integrin beta 1 (fibronectin receptor beta), H2-D1 - histocompatibility 2, d region locus 1, Lgals1 - lectin, galactose binding, soluble 1, Vim - vimentin, C1qc - complement component 1, q subcomponent, c chain, F2 - coagulation factor ii, Hrg - histidine-rich glycoprotein, Gba - glucosidase, beta, acid, Sema4a - sema domain, immunoglobulin domain (ig), transmembrane domain (tm) and short cytoplasmic domain, (semaphorin) 4a, Serpinb1a - serine (or cysteine) peptidase inhibitor, clade b, member 1a, Gfap - glial fibrillary acidic protein, Fga - fibrinogen alpha chain, Arhgdib - rho, gdp dissociation inhibitor (gdi) beta, Apoa1 - apolipoprotein a-i, Stat3 - signal transducer and activator of transcription 3, Stat1 - signal transducer and activator of transcription 1, Anxa2 - annexin a2] |
| GO:0002708 | positive regulation of lymphocyte mediated immunity | 1.79E-6 | 2.62E-4 | 10,80 | 4935 | 20 | 160 | 7 | [Hpx - hemopexin, C3 - complement component 3, H2-D1 - histocompatibility 2, d region locus 1, Ighg2b - immunoglobulin heavy constant gamma 2b, Ighg1 - immunoglobulin heavy constant gamma 1 (g1m marker), Fcer1g - fc receptor, ige, high affinity i, gamma polypeptide, Lamp1 - lysosomal-associated membrane protein 1] |
| GO:0032268 | regulation of cellular protein metabolic process | 1.89E-6 | 2.72E-4 | 1,86 | 4935 | 861 | 160 | 52 | [Pzp - pregnancy zone protein, Wfs1 - wolfram syndrome 1 homolog (human), Serpina1b - serine (or cysteine) preptidase inhibitor, clade a, member 1b, Ahsg - alpha-2-hs-glycoprotein, Serpina1a - serine (or cysteine) peptidase inhibitor, clade a, member 1a, Fn1 - fibronectin 1, Aif1 - allograft inflammatory factor 1, Gsn - gelsolin, Serpina1d - serine (or cysteine) peptidase inhibitor, clade a, member 1d, Serpina1e - serine (or cysteine) peptidase inhibitor, clade a, member 1e, Ppp1r1b - protein phosphatase 1, regulatory (inhibitor) subunit 1b, Serpina3k - serine (or cysteine) peptidase inhibitor, clade a, member 3k, Kng1 - kininogen 1, Glipr2 - gli pathogenesis-related 2, Itgb1 - integrin beta 1 (fibronectin receptor beta), Vim - vimentin, F2 - coagulation factor ii, Psme1 - proteasome (prosome, macropain) activator subunit 1 (pa28 alpha), Vtn - vitronectin, Itgb3 - integrin beta 3, Itgb2 - integrin beta 2, Gba - glucosidase, beta, acid, Serpinb1a - serine (or cysteine) peptidase inhibitor, clade b, member 1a, Itih3 - inter-alpha trypsin inhibitor, heavy chain 3, Itih2 - inter-alpha trypsin inhibitor, heavy chain 2, Ighm - immunoglobulin heavy constant mu, Itih1 - inter-alpha trypsin inhibitor, heavy chain 1, Stat3 - signal transducer and activator of transcription 3, Stat1 - signal transducer and activator of transcription 1, Anxa2 - annexin a2, Serpina6 - serine (or cysteine) peptidase inhibitor, clade a, member 6, Serpinh1 - serine (or cysteine) peptidase inhibitor, clade h, member 1, Cst3 - cystatin c, Serpinc1 - serine (or cysteine) peptidase inhibitor, clade c (antithrombin), member 1, Lgmn - legumain, Fgg - fibrinogen gamma chain, Fhit - fragile histidine triad gene, Ctsd - cathepsin d, Sod1 - superoxide dismutase 1, soluble, Fgb - fibrinogen beta chain, Ide - insulin degrading enzyme, C3 - complement component 3, Hrg - histidine-rich glycoprotein, Ddrgk1 - ddrgk domain containing 1, Hpx - hemopexin, Fga - fibrinogen alpha chain, Clu - clusterin, Apoa1 - apolipoprotein a-i, Ncstn - nicastrin, Pdcd5 - programmed cell death 5, Commd1 - comm domain containing 1, Mug1 - murinoglobulin 1] |

| | | | | | | | | | |
|---|---|---|---|---|---|---|---|---|---|
| GO:0065007 | biological regulation | 1.96E-6 | 2.78E-4 | 1,26 | 4935 | 3185 | 160 | 130 | [illegible] apolipoprotein h, Apod - apolipoprotein d, Psmg2 - proteasome (prosome, macropain) assembly chaperone 2, Fcer1g - fc receptor, ige, high affinity i, gamma polypeptide, Kng1 - kininogen 1, F2 - coagulation factor ii, Arhgap22 - rho gtpase activating protein 22, Fabp5 - fatty acid binding protein 5, epidermal, Psme1 - proteasome (prosome, macropain) activator subunit 1 (pa28 alpha), Sh3bgrl3 - sh3 domain binding glutamic acid-rich protein-like 3, Gba - glucosidase, beta, acid, Arhgdib - rho, gdp dissociation inhibitor (gdi) beta, Irs2 - insulin receptor substrate 2, Stat3 - signal transducer and activator of transcription 3, Nptxr - neuronal pentraxin receptor, Arpc1b - actin related protein 2/3 complex, subunit 1b, Stat1 - signal transducer and activator of transcription 1, Anxa2 - annexin a2, Serpina6 - serine (or cysteine) peptidase inhibitor, clade a, member 6, Plg - plasminogen, Serpinh1 - serine (or cysteine) peptidase inhibitor, clade h, member 1, Ctsb - cathepsin b, Lcp1 - lymphocyte cytosolic protein 1, Cst3 - cystatin c, Serpinc1 - serine (or cysteine) peptidase inhibitor, clade c (antithrombin), member 1, Irgm1 - immunity-related gtpase family m member 1, Fgg - fibrinogen gamma chain, Ctsd - cathepsin d, Pex19 - peroxisomal biogenesis factor 19, Gaa - glucosidase, alpha, acid, Fgb - fibrinogen beta chain, Ide - insulin degrading enzyme, Krt1 - keratin 1, S100a13 - s100 calcium binding protein a13, Hrg - histidine-rich glycoprotein, Hexb - hexosaminidase b, Car14 - carbonic anhydrase 14, Ncstn - nicastrin, Clu - clusterin, Wfs1 - wolfram syndrome 1 homolog (human), Serpina1b - serine (or cysteine) preptidase inhibitor, clade a, member 1b, Scg2 - secretogranin ii, Serpina1a - serine (or cysteine) peptidase inhibitor, clade a, member 1a, Fn1 - fibronectin 1, Ahsg - alpha-2-hs-glycoprotein, Aif1 - allograft inflammatory factor 1, Clic1 - chloride intracellular channel 1, Acot1 - acyl-coa thioesterase 1, Nes - nestin, Serpina1d - serine (or cysteine) peptidase inhibitor, clade a, member 1d, Serpina1e - serine (or cysteine) peptidase inhibitor, clade a, member 1e, Mcam - melanoma cell adhesion molecule, Serpina3k - serine (or cysteine) peptidase inhibitor, clade a, member 3k, P4hb - prolyl 4-hydroxylase, beta polypeptide, Ppp1r1b - protein phosphatase 1, regulatory (inhibitor) subunit 1b, Crip2 - cysteine rich protein 2, Alb - albumin, Ctsa - cathepsin a, Marcksl1 - marcks-like 1, Cpe - carboxypeptidase e, Glipr2 - gli pathogenesis-related 2, Igkc - immunoglobulin kappa constant, Itgb1 - integrin beta 1 (fibronectin receptor beta), Vim - vimentin, Itgam - integrin alpha m, Cp - ceruloplasmin, Fabp7 - fatty acid binding protein 7, brain, Itga6 - integrin alpha 6, Vtn - vitronectin, Itgb3 - integrin beta 3, Tpm1 - tropomyosin 1, alpha, Rbp1 - retinol binding protein 1, cellular, Itgb2 - integrin beta 2, Scarb2 - scavenger receptor class b, member 2, Serpinb1a - serine (or cysteine) peptidase inhibitor, clade b, member 1a, Ighg2b - immunoglobulin heavy constant gamma 2b, Itih3 - inter-alpha trypsin inhibitor, heavy chain 3, Orm1 - orosomucoid 1, Ighg1 - immunoglobulin heavy constant gamma 1 (g1m marker), Itih2 - inter-alpha trypsin inhibitor, heavy chain 2, Ampd3 - adenosine monophosphate deaminase 3, Ighm - immunoglobulin heavy constant mu, Itih1 - inter-alpha trypsin inhibitor, heavy chain 1, Cd151 - cd151 antigen, Scn3b - sodium channel, voltage-gated, type iii, beta, Anxa3 - annexin a3, Ier3ip1 - immediate early response 3 interacting protein 1, Lamp2 - lysosomal-associated membrane protein 2, Lamp1 - lysosomal-associated membrane protein 1, Anpep - alanyl (membrane) aminopeptidase, Gjb6 - gap junction protein, beta 6, Lgmn - legumain, Fhit - fragile histidine triad gene, Cfh - complement component factor h, Ttr - transthyretin, Sod1 - superoxide dismutase 1, soluble, C3 - complement component 3, H2-D1 - histocompatibility 2, d region locus 1, C4b - complement component 4b (chido blood group), Lgals1 - lectin, galactose binding, soluble 1, Ppp1r1a - protein phosphatase 1, regulatory (inhibitor) subunit 1a, Cfb - complement factor b, Hp - haptoglobin, C1qc - complement component 1, q subcomponent, c chain, Lingo2 - leucine rich repeat and ig domain containing 2, C1qa - complement component 1, q subcomponent, alpha polypeptide, C1qb - complement component 1, q subcomponent, beta polypeptide, Kcnj10 - potassium inwardly-rectifying channel, subfamily j, member 10, Ddrgk1 - ddrgk domain containing 1, Lsm6 - lsm6 homolog, u6 small nuclear rna associated (s. cerevisiae), Ybx1 - y box protein 1, Tgm1 - transglutaminase 1, k polypeptide, Hpx - hemopexin, Tgm2 - transglutaminase 2, c polypeptide, Sema4a - sema domain, |
| GO:0098542 | defense response to other organism | 2.18E-6 | 3.06E-4 | 5,21 | 4935 | 71 | 160 | 12 | [Igkc - immunoglobulin kappa constant, Anxa3 - annexin a3, Hp - haptoglobin, Ighg2b - immunoglobulin heavy constant gamma 2b, Fga - fibrinogen alpha chain, Ighg1 - immunoglobulin heavy constant gamma 1 (g1m marker), Ighm - immunoglobulin heavy constant mu, F2 - coagulation factor ii, Fcer1g - fc receptor, ige, high affinity i, gamma polypeptide, Hrg - histidine-rich glycoprotein, Fgb - fibrinogen beta chain, Stat1 - signal transducer and activator of transcription 1] |
| GO:0006950 | response to stress | 2.18E-6 | 3.03E-4 | 1,93 | 4935 | 769 | 160 | 48 | [Serpina1b - serine (or cysteine) preptidase inhibitor, clade a, member 1b, Wfs1 - wolfram syndrome 1 homolog (human), Anxa3 - annexin a3, Fn1 - fibronectin 1, Ahsg - alpha-2-hs-glycoprotein, Aif1 - allograft inflammatory factor 1, Lcp1 - lymphocyte cytosolic protein 1, Cst3 - cystatin c, Lamp2 - lysosomal-associated membrane protein 2, Apod - apolipoprotein d, Irgm1 - immunity-related gtpase family m member 1, P4hb - prolyl 4-hydroxylase, beta polypeptide, Fgg - fibrinogen gamma chain, Fcer1g - fc receptor, ige, high affinity i, gamma polypeptide, Alb - albumin, Cfh - complement component factor h, Sod1 - superoxide dismutase 1, soluble, Fgb - fibrinogen beta chain, Kng1 - kininogen 1, Ide - insulin degrading enzyme, Krt1 - keratin 1, C3 - complement component 3, Igkc - immunoglobulin kappa constant, Itgam - integrin alpha m, Cfb - complement factor b, Hp - haptoglobin, C1qc - complement component 1, q subcomponent, c chain, F2 - coagulation factor ii, Mgst1 - microsomal glutathione s-transferase 1, C1qa - complement component 1, q subcomponent, alpha polypeptide, Hrg - histidine-rich glycoprotein, C1qb - complement component 1, q subcomponent, beta polypeptide, Ddrgk1 - ddrgk domain containing 1, Tpm1 - tropomyosin 1, alpha, Gba - glucosidase, beta, acid, Orm1 - orosomucoid 1, Ighg2b - immunoglobulin heavy constant gamma 2b, Serpinb1a - serine (or cysteine) peptidase inhibitor, clade b, member 1a, Gfap - glial fibrillary acidic protein, Fga - fibrinogen alpha chain, Ighg1 - immunoglobulin heavy constant gamma 1 (g1m marker), Ighm - immunoglobulin heavy constant mu, Clu - clusterin, Commd1 - comm domain containing 1, Tspan2 - tetraspanin 2, Stat3 - signal transducer and activator of transcription 3, Stat1 - signal transducer and activator of transcription 1, Tbl2 - transducin (beta)-like 2] |
| GO:0045785 | positive regulation of cell adhesion | 2.55E-6 | 3.49E-4 | 3,89 | 4935 | 127 | 160 | 16 | [Itgb1 - integrin beta 1 (fibronectin receptor beta), Lgals1 - lectin, galactose binding, soluble 1, Fn1 - fibronectin 1, Aif1 - allograft inflammatory factor 1, Itga6 - integrin alpha 6, Vtn - vitronectin, Itgb3 - integrin beta 3, Tpm1 - tropomyosin 1, alpha, Itgb2 - integrin beta 2, Flna - filamin, alpha, Tgm2 - transglutaminase 2, c polypeptide, Fga - fibrinogen alpha chain, Fgg - fibrinogen gamma chain, P4hb - prolyl 4-hydroxylase, beta polypeptide, Apoa1 - apolipoprotein a-i, Fgb - fibrinogen beta chain] |
| GO:0031638 | zymogen activation | 2.56E-6 | 3.47E-4 | 13,22 | 4935 | 14 | 160 | 6 | [Apoh - apolipoprotein h, Hp - haptoglobin, Lgmn - legumain, Fga - fibrinogen alpha chain, Fgg - fibrinogen gamma chain, Fgb - fibrinogen beta chain] |

| | | | | | | | | | |
|---|---|---|---|---|---|---|---|---|---|
| GO:0002460 | adaptive immune response based on somatic recombination of immune receptors built from immunoglobulin superfamily domains | 2.62E-6 | 3.51E-4 | 10,28 | 4935 | 21 | 160 | 7 | [C4b - complement component 4b (chido blood group), Ighg2b - immunoglobulin heavy constant gamma 2b, Ighg1 - immunoglobulin heavy constant gamma 1 (g1m marker), Fcer1g - fc receptor, ige, high affinity i, gamma polypeptide, Ighm - immunoglobulin heavy constant mu, Cfh - complement component factor h, Stat3 - signal transducer and activator of transcription 3] |
| GO:0009605 | response to external stimulus | 2.92E-6 | 3.86E-4 | 2,55 | 4935 | 339 | 160 | 28 | [Anxa3 - annexin a3, Gsn - gelsolin, Lamp2 - lysosomal-associated membrane protein 2, Irgm1 - immunity-related gtpase family m member 1, Serpinc1 - serine (or cysteine) peptidase inhibitor, clade c (antithrombin), member 1, Alb - albumin, Fcer1g - fc receptor, ige, high affinity i, gamma polypeptide, Cfh - complement component factor h, Sod1 - superoxide dismutase 1, soluble, Fgb - fibrinogen beta chain, C3 - complement component 3, Igkc - immunoglobulin kappa constant, Vim - vimentin, Hp - haptoglobin, F2 - coagulation factor ii, Itga6 - integrin alpha 6, Fabp7 - fatty acid binding protein 7, brain, Mgst1 - microsomal glutathione s-transferase 1, Hrg - histidine-rich glycoprotein, Gba - glucosidase, beta, acid, Rbp1 - retinol binding protein 1, cellular, Ighg2b - immunoglobulin heavy constant gamma 2b, Ighg1 - immunoglobulin heavy constant gamma 1 (g1m marker), Fga - fibrinogen alpha chain, Ighm - immunoglobulin heavy constant mu, Stat1 - signal transducer and activator of transcription 1, Tbl2 - transducin (beta)-like 2, Ces1c - carboxylesterase 1c] |
| GO:0009607 | response to biotic stimulus | 3.7E-6 | 4.84E-4 | 3,43 | 4935 | 162 | 160 | 18 | [C3 - complement component 3, Igkc - immunoglobulin kappa constant, Anxa3 - annexin a3, Vim - vimentin, Hp - haptoglobin, F2 - coagulation factor ii, Mgst1 - microsomal glutathione s-transferase 1, Hrg - histidine-rich glycoprotein, Irgm1 - immunity-related gtpase family m member 1, Ighg2b - immunoglobulin heavy constant gamma 2b, Fga - fibrinogen alpha chain, Ighg1 - immunoglobulin heavy constant gamma 1 (g1m marker), Fcer1g - fc receptor, ige, high affinity i, gamma polypeptide, Ighm - immunoglobulin heavy constant mu, Cfh - complement component factor h, Stat1 - signal transducer and activator of transcription 1, Fgb - fibrinogen beta chain, Ces1c - carboxylesterase 1c] |
| GO:0002712 | regulation of B cell mediated immunity | 3.82E-6 | 4.94E-4 | 17,14 | 4935 | 9 | 160 | 5 | [Hpx - hemopexin, C3 - complement component 3, Ighg2b - immunoglobulin heavy constant gamma 2b, Ighg1 - immunoglobulin heavy constant gamma 1 (g1m marker), Fcer1g - fc receptor, ige, high affinity i, gamma polypeptide] |
| GO:0002889 | regulation of immunoglobulin mediated immune response | 3.82E-6 | 4.88E-4 | 17,14 | 4935 | 9 | 160 | 5 | [Hpx - hemopexin, C3 - complement component 3, Ighg2b - immunoglobulin heavy constant gamma 2b, Ighg1 - immunoglobulin heavy constant gamma 1 (g1m marker), Fcer1g - fc receptor, ige, high affinity i, gamma polypeptide] |
| GO:0051248 | negative regulation of protein metabolic process | 4.44E-6 | 5.61E-4 | 2,35 | 4935 | 407 | 160 | 31 | [Serpina1b - serine (or cysteine) preptidase inhibitor, clade a, member 1b, Pzp - pregnancy zone protein, Serpina6 - serine (or cysteine) peptidase inhibitor, clade a, member 6, Serpina1a - serine (or cysteine) peptidase inhibitor, clade a, member 1a, Ahsg - alpha-2-hs-glycoprotein, Serpinh1 - serine (or cysteine) peptidase inhibitor, clade h, member 1, Cst3 - cystatin c, Serpina1d - serine (or cysteine) peptidase inhibitor, clade a, member 1d, Serpina1e - serine (or cysteine) peptidase inhibitor, clade a, member 1e, Flna - filamin, alpha, Apod - apolipoprotein d, Serpinc1 - serine (or cysteine) peptidase inhibitor, clade c (antithrombin), member 1, Ppp1r1b - protein phosphatase 1, regulatory (inhibitor) subunit 1b, Serpina3k - serine (or cysteine) peptidase inhibitor, clade a, member 3k, Fhit - fragile histidine triad gene, Ctsa - cathepsin a, Kng1 - kininogen 1, Ide - insulin degrading enzyme, F2 - coagulation factor ii, Hrg - histidine-rich glycoprotein, Ddrgk1 - ddrgk domain containing 1, Vtn - vitronectin, Itgb3 - integrin beta 3, Gba - glucosidase, beta, acid, Itih3 - inter-alpha trypsin inhibitor, heavy chain 3, Serpinb1a - serine (or cysteine) peptidase inhibitor, clade b, member 1a, Itih2 - inter-alpha trypsin inhibitor, heavy chain 2, Itih1 - inter-alpha trypsin inhibitor, heavy chain 1, Clu - clusterin, Mug1 - murinoglobulin 1, Anxa2 - annexin a2] |

| | | | | | | | | | |
|---|---|---|---|---|---|---|---|---|---|
| GO:0032501 | multicellular organismal process | 5.15E-6 | 6.44E-4 | 1,77 | 4935 | 939 | 160 | 54 | [Wfs1 - wolfram syndrome 1 homolog (human), Serpina1b - serine (or cysteine) preptidase inhibitor, clade a, member 1b, Ahsg - alpha-2-hs-glycoprotein, Aif1 - allograft inflammatory factor 1, Gsn - gelsolin, Nes - nestin, Flna - filamin, alpha, Apoh - apolipoprotein h, Ppp1r1b - protein phosphatase 1, regulatory (inhibitor) subunit 1b, Fcer1g - fc receptor, ige, high affinity i, gamma polypeptide, Ndufs6 - nadh dehydrogenase (ubiquinone) fe-s protein 6, Kng1 - kininogen 1, Itgb1 - integrin beta 1 (fibronectin receptor beta), Itgam - integrin alpha m, Fabp7 - fatty acid binding protein 7, brain, F2 - coagulation factor ii, Alg5 - asparagine-linked glycosylation 5 (dolichyl-phosphate beta-glucosyltransferase), Arhgap22 - rho gtpase activating protein 22, Tpm1 - tropomyosin 1, alpha, Itgb2 - integrin beta 2, Gba - glucosidase, beta, acid, Ampd3 - adenosine monophosphate deaminase 3, Stat3 - signal transducer and activator of transcription 3, Stat1 - signal transducer and activator of transcription 1, Scn3b - sodium channel, voltage-gated, type iii, beta, Plg - plasminogen, Tpm4 - tropomyosin 4, Anpep - alanyl (membrane) aminopeptidase, Gjb6 - gap junction protein, beta 6, Lgmn - legumain, Serpinc1 - serine (or cysteine) peptidase inhibitor, clade c (antithrombin), member 1, Fgg - fibrinogen gamma chain, Cfh - complement component factor h, Gaa - glucosidase, alpha, acid, Sod1 - superoxide dismutase 1, soluble, Fgb - fibrinogen beta chain, Ide - insulin degrading enzyme, C3 - complement component 3, Krt1 - keratin 1, S100a13 - s100 calcium binding protein a13, C1qa - complement component 1, q subcomponent, alpha polypeptide, Hrg - histidine-rich glycoprotein, Kcnj10 - potassium inwardly-rectifying channel, subfamily j, member 10, Hexb - hexosaminidase b, Ybx1 - y box protein 1, Tgm1 - transglutaminase 1, k polypeptide, Sema4a - sema domain, immunoglobulin domain (ig), transmembrane domain (tm) and short cytoplasmic domain, (semaphorin) 4a, Tgm2 - transglutaminase 2, c polypeptide, Fga - fibrinogen alpha chain, Krt71 - keratin 71, Apoa1 - apolipoprotein a-i, Ncstn - nicastrin, Nptx2 - neuronal pentraxin 2, Commd1 - comm domain containing 1] |
| GO:0072378 | blood coagulation, fibrin clot formation | 5.19E-6 | 6.42E-4 | 24,68 | 4935 | 5 | 160 | 4 | [Fn1 - fibronectin 1, Fga - fibrinogen alpha chain, Fgg - fibrinogen gamma chain, Fgb - fibrinogen beta chain] |
| GO:0002443 | leukocyte mediated immunity | 5.23E-6 | 6.4E-4 | 9,39 | 4935 | 23 | 160 | 7 | [C4b - complement component 4b (chido blood group), Ighg2b - immunoglobulin heavy constant gamma 2b, Ighg1 - immunoglobulin heavy constant gamma 1 (g1m marker), Ighm - immunoglobulin heavy constant mu, Fcer1g - fc receptor, ige, high affinity i, gamma polypeptide, F2 - coagulation factor ii, Cfh - complement component factor h] |
| GO:0050789 | regulation of biological process | 5.34E-6 | 6.47E-4 | 1,27 | 4935 | 3013 | 160 | 124 | [illegible] apolipoprotein d, Psmg2 - proteasome (prosome, macropain) assembly chaperone 2, Fcer1g - fc receptor, ige, high affinity i, gamma polypeptide, Kng1 - kininogen 1, F2 - coagulation factor ii, Arhgap22 - rho gtpase activating protein 22, Fabp5 - fatty acid binding protein 5, epidermal, Psme1 - proteasome (prosome, macropain) activator subunit 1 (pa28 alpha), Sh3bgrl3 - sh3 domain binding glutamic acid-rich protein-like 3, Gba - glucosidase, beta, acid, Arhgdib - rho, gdp dissociation inhibitor (gdi) beta, Irs2 - insulin receptor substrate 2, Stat3 - signal transducer and activator of transcription 3, Nptxr - neuronal pentraxin receptor, Arpc1b - actin related protein 2/3 complex, subunit 1b, Stat1 - signal transducer and activator of transcription 1, Anxa2 - annexin a2, Serpina6 - serine (or cysteine) peptidase inhibitor, clade a, member 6, Plg - plasminogen, Serpinh1 - serine (or cysteine) peptidase inhibitor, clade h, member 1, Ctsb - cathepsin b, Lcp1 - lymphocyte cytosolic protein 1, Cst3 - cystatin c, Serpinc1 - serine (or cysteine) peptidase inhibitor, clade c (antithrombin), member 1, Irgm1 - immunity-related gtpase family m member 1, Fgg - fibrinogen gamma chain, Ctsd - cathepsin d, Gaa - glucosidase, alpha, acid, Fgb - fibrinogen beta chain, Ide - insulin degrading enzyme, Krt1 - keratin 1, S100a13 - s100 calcium binding protein a13, Hrg - histidine-rich glycoprotein, Hexb - hexosaminidase b, Clu - clusterin, Ncstn - nicastrin, Serpina1b - serine (or cysteine) preptidase inhibitor, clade a, member 1b, Wfs1 - wolfram syndrome 1 homolog (human), Scg2 - secretogranin ii, Ahsg - alpha-2-hs-glycoprotein, Serpina1a - serine (or cysteine) peptidase inhibitor, clade a, member 1a, Fn1 - fibronectin 1, Clic1 - chloride intracellular channel 1, Aif1 - allograft inflammatory factor 1, Acot1 - acyl-coa thioesterase 1, Nes - nestin, Serpina1d - serine (or cysteine) peptidase inhibitor, clade a, member 1d, Serpina1e - serine (or cysteine) peptidase inhibitor, clade a, member 1e, Mcam - melanoma cell adhesion molecule, P4hb - prolyl 4-hydroxylase, beta polypeptide, Ppp1r1b - protein phosphatase 1, regulatory (inhibitor) subunit 1b, Serpina3k - serine (or cysteine) peptidase inhibitor, clade a, member 3k, Crip2 - cysteine rich protein 2, Alb - albumin, Ctsa - cathepsin a, Marcksl1 - marcks-like 1, Cpe - carboxypeptidase e, Glipr2 - gli pathogenesis-related 2, Igkc - immunoglobulin kappa constant, Itgb1 - integrin beta 1 (fibronectin receptor beta), Itgam - integrin alpha m, Vim - vimentin, Itga6 - integrin alpha 6, Fabp7 - fatty acid binding protein 7, brain, Vtn - vitronectin, Itgb3 - integrin beta 3, Tpm1 - tropomyosin 1, alpha, Itgb2 - integrin beta 2, Rbp1 - retinol binding protein 1, cellular, Scarb2 - scavenger receptor class b, member 2, Orm1 - orosomucoid 1, Itih3 - inter-alpha trypsin inhibitor, heavy chain 3, Serpinb1a - serine (or cysteine) peptidase inhibitor, clade b, member 1a, Ighg2b - immunoglobulin heavy constant gamma 2b, Itih2 - inter-alpha trypsin inhibitor, heavy chain 2, Ighg1 - immunoglobulin heavy constant gamma 1 (g1m marker), Itih1 - inter-alpha trypsin inhibitor, heavy chain 1, Ighm - immunoglobulin heavy constant mu, Cd151 - cd151 antigen, Scn3b - sodium channel, voltage-gated, type iii, beta, Anxa3 - annexin a3, Ier3ip1 - immediate early response 3 interacting protein 1, Lamp2 - lysosomal-associated membrane protein 2, Lamp1 - lysosomal-associated membrane protein 1, Anpep - alanyl (membrane) aminopeptidase, Gjb6 - gap junction protein, beta 6, Lgmn - legumain, Fhit - fragile histidine triad gene, Cfh - complement component factor h, Sod1 - superoxide dismutase 1, soluble, C3 - complement component 3, H2-D1 - histocompatibility 2, d region locus 1, Lgals1 - lectin, galactose binding, soluble 1, C4b - complement component 4b (chido blood group), Ppp1r1a - protein phosphatase 1, regulatory (inhibitor) subunit 1a, Cfb - complement factor b, Hp - haptoglobin, C1qc - complement component 1, q subcomponent, c chain, Lingo2 - leucine rich repeat and ig domain containing 2, C1qa - complement component 1, q subcomponent, alpha polypeptide, Kcnj10 - potassium inwardly-rectifying channel, subfamily j, member 10, C1qb - complement component 1, q subcomponent, beta polypeptide, Ddrgk1 - ddrgk domain containing 1, Lsm6 - lsm6 homolog, u6 small nuclear rna associated (s. cerevisiae), Tgm1 - transglutaminase 1, k polypeptide, Ybx1 - y box protein 1, Hpx - hemopexin, Sema4a - sema domain, immunoglobulin domain (ig), transmembrane domain (tm) and short cytoplasmic domain, (semaphorin) 4a, Tgm2 - transglutaminase 2, c polypeptide, Fga - fibrinogen alpha chain, Gfap - glial fibrillary acidic |

| | | | | | | | | | |
|---|---|---|---|---|---|---|---|---|---|
| GO:0031347 | regulation of defense response | 6.24E-6 | 7.48E-4 | 3,30 | 4935 | 168 | 160 | 18 | [C3 - complement component 3, Krt1 - keratin 1, Ahsg - alpha-2-hs-glycoprotein, F2 - coagulation factor ii, Lamp1 - lysosomal-associated membrane protein 1, Hpx - hemopexin, Tgm2 - transglutaminase 2, c polypeptide, Apod - apolipoprotein d, Irgm1 - immunity-related gtpase family m member 1, Serpinb1a - serine (or cysteine) peptidase inhibitor, clade b, member 1a, Ighg2b - immunoglobulin heavy constant gamma 2b, Ighg1 - immunoglobulin heavy constant gamma 1 (g1m marker), Fcer1g - fc receptor, ige, high affinity i, gamma polypeptide, Apoa1 - apolipoprotein a-i, Cfh - complement component factor h, Stat3 - signal transducer and activator of transcription 3, Sod1 - superoxide dismutase 1, soluble, Stat1 - signal transducer and activator of transcription 1] |
| GO:0002888 | positive regulation of myeloid leukocyte mediated immunity | 6.47E-6 | 7.67E-4 | 11,57 | 4935 | 16 | 160 | 6 | [C3 - complement component 3, Itgam - integrin alpha m, Ighg2b - immunoglobulin heavy constant gamma 2b, Ighg1 - immunoglobulin heavy constant gamma 1 (g1m marker), Fcer1g - fc receptor, ige, high affinity i, gamma polypeptide, Itgb2 - integrin beta 2] |
| GO:0002697 | regulation of immune effector process | 6.5E-6 | 7.63E-4 | 4,36 | 4935 | 92 | 160 | 13 | [C3 - complement component 3, H2-D1 - histocompatibility 2, d region locus 1, Itgam - integrin alpha m, Lamp1 - lysosomal-associated membrane protein 1, Itgb2 - integrin beta 2, Hpx - hemopexin, Ighg2b - immunoglobulin heavy constant gamma 2b, Ighg1 - immunoglobulin heavy constant gamma 1 (g1m marker), Fcer1g - fc receptor, ige, high affinity i, gamma polypeptide, Ighm - immunoglobulin heavy constant mu, Cfh - complement component factor h, Apoa1 - apolipoprotein a-i, Stat1 - signal transducer and activator of transcription 1] |
| GO:0002696 | positive regulation of leukocyte activation | 6.78E-6 | 7.87E-4 | 5,14 | 4935 | 66 | 160 | 11 | [Igkc - immunoglobulin kappa constant, Lgals1 - lectin, galactose binding, soluble 1, Itgam - integrin alpha m, Ighg2b - immunoglobulin heavy constant gamma 2b, Ighg1 - immunoglobulin heavy constant gamma 1 (g1m marker), Aif1 - allograft inflammatory factor 1, Ighm - immunoglobulin heavy constant mu, Fcer1g - fc receptor, ige, high affinity i, gamma polypeptide, Irs2 - insulin receptor substrate 2, Lamp1 - lysosomal-associated membrane protein 1, Itgb2 - integrin beta 2] |
| GO:0045807 | positive regulation of endocytosis | 7.35E-6 | 8.45E-4 | 4,31 | 4935 | 93 | 160 | 13 | [C3 - complement component 3, Itgb1 - integrin beta 1 (fibronectin receptor beta), Ahsg - alpha-2-hs-glycoprotein, Vtn - vitronectin, Ighg2b - immunoglobulin heavy constant gamma 2b, Ighg1 - immunoglobulin heavy constant gamma 1 (g1m marker), Fcer1g - fc receptor, ige, high affinity i, gamma polypeptide, Ighm - immunoglobulin heavy constant mu, Apoa1 - apolipoprotein a-i, Clu - clusterin, Cd151 - cd151 antigen, Sod1 - superoxide dismutase 1, soluble, Anxa2 - annexin a2] |
| GO:0042127 | regulation of cell proliferation | 7.48E-6 | 8.51E-4 | 2,38 | 4935 | 376 | 160 | 29 | [Scg2 - secretogranin ii, Fn1 - fibronectin 1, Mvd - mevalonate (diphospho) decarboxylase, Aif1 - allograft inflammatory factor 1, Cst3 - cystatin c, Flna - filamin, alpha, Apoh - apolipoprotein h, Gjb6 - gap junction protein, beta 6, Apod - apolipoprotein d, Lgmn - legumain, Crip2 - cysteine rich protein 2, Marcksl1 - marcks-like 1, Itgb1 - integrin beta 1 (fibronectin receptor beta), Vim - vimentin, F2 - coagulation factor ii, S100a13 - s100 calcium binding protein a13, Ddrgk1 - ddrgk domain containing 1, Itgb3 - integrin beta 3, Ybx1 - y box protein 1, Tgm1 - transglutaminase 1, k polypeptide, Tgm2 - transglutaminase 2, c polypeptide, Gfap - glial fibrillary acidic protein, Ighm - immunoglobulin heavy constant mu, Pdcd5 - programmed cell death 5, Irs2 - insulin receptor substrate 2, Clu - clusterin, Stat3 - signal transducer and activator of transcription 3, Stat1 - signal transducer and activator of transcription 1, Anxa2 - annexin a2] |
| GO:0045321 | leukocyte activation | 7.98E-6 | 8.99E-4 | 4,00 | 4935 | 108 | 160 | 14 | [Igkc - immunoglobulin kappa constant, Anxa3 - annexin a3, Itgam - integrin alpha m, Lgals1 - lectin, galactose binding, soluble 1, Aif1 - allograft inflammatory factor 1, Lcp1 - lymphocyte cytosolic protein 1, C1qa - complement component 1, q subcomponent, alpha polypeptide, Gba - glucosidase, beta, acid, Itgb2 - integrin beta 2, Sema4a - sema domain, immunoglobulin domain (ig), transmembrane domain (tm) and short cytoplasmic domain, (semaphorin) 4a, Fcer1g - fc receptor, ige, high affinity i, gamma polypeptide, Clu - clusterin, Ncstn - nicastrin, Cd151 - cd151 antigen] |
| GO:0070527 | platelet aggregation | 9.73E-6 | 1.09E-3 | 10,89 | 4935 | 17 | 160 | 6 | [Fn1 - fibronectin 1, Fga - fibrinogen alpha chain, Fgg - fibrinogen gamma chain, Cfh - complement component factor h, Itgb3 - integrin beta 3, Fgb - fibrinogen beta chain] |
| GO:0002824 | positive regulation of adaptive immune response based on somatic recombination of immune receptors built from immunoglobulin superfamily domains | 9.73E-6 | 1.08E-3 | 10,89 | 4935 | 17 | 160 | 6 | [Hpx - hemopexin, C3 - complement component 3, H2-D1 - histocompatibility 2, d region locus 1, Ighg2b - immunoglobulin heavy constant gamma 2b, Ighg1 - immunoglobulin heavy constant gamma 1 (g1m marker), Fcer1g - fc receptor, ige, high affinity i, gamma polypeptide] |

| | | | | | | | | | |
|---|---|---|---|---|---|---|---|---|---|
| GO:0002703 | regulation of leukocyte mediated immunity | 1.03E-5 | 1.13E-3 | 6,17 | 4935 | 45 | 160 | 9 | [Hpx - hemopexin, C3 - complement component 3, H2-D1 - histocompatibility 2, d region locus 1, Itgam - integrin alpha m, Ighg2b - immunoglobulin heavy constant gamma 2b, Ighg1 - immunoglobulin heavy constant gamma 1 (g1m marker), Fcer1g - fc receptor, ige, high affinity i, gamma polypeptide, Lamp1 - lysosomal-associated membrane protein 1, Itgb2 - integrin beta 2] |
| GO:0051704 | multi-organism process | 1.03E-5 | 1.12E-3 | 2,77 | 4935 | 245 | 160 | 22 | [C3 - complement component 3, Igkc - immunoglobulin kappa constant, Plg - plasminogen, Anxa3 - annexin a3, Hp - haptoglobin, Fn1 - fibronectin 1, F2 - coagulation factor ii, Ctsb - cathepsin b, Hrg - histidine-rich glycoprotein, Itgb3 - integrin beta 3, Hexb - hexosaminidase b, Irgm1 - immunity-related gtpase family m member 1, Ighg2b - immunoglobulin heavy constant gamma 2b, Fga - fibrinogen alpha chain, Ighg1 - immunoglobulin heavy constant gamma 1 (g1m marker), Fcer1g - fc receptor, ige, high affinity i, gamma polypeptide, Ighm - immunoglobulin heavy constant mu, Stat3 - signal transducer and activator of transcription 3, Fgb - fibrinogen beta chain, Stat1 - signal transducer and activator of transcription 1, Anxa2 - annexin a2, Ces1c - carboxylesterase 1c] |
| GO:0050727 | regulation of inflammatory response | 1.05E-5 | 1.13E-3 | 4,18 | 4935 | 96 | 160 | 13 | [Krt1 - keratin 1, C3 - complement component 3, Ahsg - alpha-2-hs-glycoprotein, F2 - coagulation factor ii, Tgm2 - transglutaminase 2, c polypeptide, Apod - apolipoprotein d, Ighg2b - immunoglobulin heavy constant gamma 2b, Ighg1 - immunoglobulin heavy constant gamma 1 (g1m marker), Fcer1g - fc receptor, ige, high affinity i, gamma polypeptide, Cfh - complement component factor h, Apoa1 - apolipoprotein a-i, Stat3 - signal transducer and activator of transcription 3, Sod1 - superoxide dismutase 1, soluble] |
| GO:0048583 | regulation of response to stimulus | 1.18E-5 | 1.25E-3 | 1,63 | 4935 | 1171 | 160 | 62 | [Wfs1 - wolfram syndrome 1 homolog (human), Scg2 - secretogranin ii, Fn1 - fibronectin 1, Ahsg - alpha-2-hs-glycoprotein, Aif1 - allograft inflammatory factor 1, Gsn - gelsolin, Flna - filamin, alpha, Apoh - apolipoprotein h, Apod - apolipoprotein d, P4hb - prolyl 4-hydroxylase, beta polypeptide, Fcer1g - fc receptor, ige, high affinity i, gamma polypeptide, Kng1 - kininogen 1, Igkc - immunoglobulin kappa constant, Glipr2 - gli pathogenesis-related 2, Itgb1 - integrin beta 1 (fibronectin receptor beta), Itgam - integrin alpha m, F2 - coagulation factor ii, Fabp7 - fatty acid binding protein 7, brain, Itga6 - integrin alpha 6, Fabp5 - fatty acid binding protein 5, epidermal, Itgb3 - integrin beta 3, Gba - glucosidase, beta, acid, Itgb2 - integrin beta 2, Ighg2b - immunoglobulin heavy constant gamma 2b, Serpinb1a - serine (or cysteine) peptidase inhibitor, clade b, member 1a, Ighg1 - immunoglobulin heavy constant gamma 1 (g1m marker), Arhgdib - rho, gdp dissociation inhibitor (gdi) beta, Ighm - immunoglobulin heavy constant mu, Nptxr - neuronal pentraxin receptor, Stat3 - signal transducer and activator of transcription 3, Stat1 - signal transducer and activator of transcription 1, Anxa2 - annexin a2, Plg - plasminogen, Lamp1 - lysosomal-associated membrane protein 1, Lgmn - legumain, Irgm1 - immunity-related gtpase family m member 1, Serpinc1 - serine (or cysteine) peptidase inhibitor, clade c (antithrombin), member 1, Fgg - fibrinogen gamma chain, Cfh - complement component factor h, Sod1 - superoxide dismutase 1, soluble, Fgb - fibrinogen beta chain, C3 - complement component 3, Krt1 - keratin 1, H2-D1 - histocompatibility 2, d region locus 1, C4b - complement component 4b (chido blood group), Cfb - complement factor b, C1qc - complement component 1, q subcomponent, c chain, S100a13 - s100 calcium binding protein a13, C1qa - complement component 1, q subcomponent, alpha polypeptide, Hrg - histidine-rich glycoprotein, C1qb - complement component 1, q subcomponent, beta polypeptide, Ddrgk1 - ddrgk domain containing 1, Hpx - hemopexin, Ybx1 - y box protein 1, Tgm2 - transglutaminase 2, c polypeptide, Sema4a - sema domain, immunoglobulin domain (ig), transmembrane domain (tm) and short cytoplasmic domain, (semaphorin) 4a, Fga - fibrinogen alpha chain, Pdcd5 - programmed cell death 5, Apoa1 - apolipoprotein a-i, Clu - clusterin, Nptx2 - neuronal pentraxin 2, Commd1 - comm domain containing 1] |
| GO:0002449 | lymphocyte mediated immunity | 1.42E-5 | 1.5E-3 | 10,28 | 4935 | 18 | 160 | 6 | [C4b - complement component 4b (chido blood group), Ighg2b - immunoglobulin heavy constant gamma 2b, Ighg1 - immunoglobulin heavy constant gamma 1 (g1m marker), Fcer1g - fc receptor, ige, high affinity i, gamma polypeptide, Ighm - immunoglobulin heavy constant mu, Cfh - complement component factor h] |
| GO:0002866 | positive regulation of acute inflammatory response to antigenic stimulus | 1.52E-5 | 1.58E-3 | 20,56 | 4935 | 6 | 160 | 4 | [C3 - complement component 3, Ighg2b - immunoglobulin heavy constant gamma 2b, Ighg1 - immunoglobulin heavy constant gamma 1 (g1m marker), Fcer1g - fc receptor, ige, high affinity i, gamma polypeptide] |
| GO:0002864 | regulation of acute inflammatory response to antigenic stimulus | 1.52E-5 | 1.57E-3 | 20,56 | 4935 | 6 | 160 | 4 | [C3 - complement component 3, Ighg2b - immunoglobulin heavy constant gamma 2b, Ighg1 - immunoglobulin heavy constant gamma 1 (g1m marker), Fcer1g - fc receptor, ige, high affinity i, gamma polypeptide] |
| GO:0031639 | plasminogen activation | 1.52E-5 | 1.56E-3 | 20,56 | 4935 | 6 | 160 | 4 | [Apoh - apolipoprotein h, Fga - fibrinogen alpha chain, Fgg - fibrinogen gamma chain, Fgb - fibrinogen beta chain] |

| | | | | | | | | | |
|---|---|---|---|---|---|---|---|---|---|
| GO:0048519 | negative regulation of biological process | 1.61E-5 | 1.63E-3 | 1,50 | 4935 | 1564 | 160 | 76 | [Pzp - pregnancy zone protein, Serpina1b - serine (or cysteine) preptidase inhibitor, clade a, member 1b, Wfs1 - wolfram syndrome 1 homolog (human), Scg2 - secretogranin ii, Ahsg - alpha-2-hs-glycoprotein, Serpina1a - serine (or cysteine) peptidase inhibitor, clade a, member 1a, Fn1 - fibronectin 1, Aif1 - allograft inflammatory factor 1, Acot1 - acyl-coa thioesterase 1, Gsn - gelsolin, Nes - nestin, Serpina1d - serine (or cysteine) peptidase inhibitor, clade a, member 1d, Serpina1e - serine (or cysteine) peptidase inhibitor, clade a, member 1e, Flna - filamin, alpha, Apoh - apolipoprotein h, Apod - apolipoprotein d, Psmg2 - proteasome (prosome, macropain) assembly chaperone 2, Ppp1r1b - protein phosphatase 1, regulatory (inhibitor) subunit 1b, Serpina3k - serine (or cysteine) peptidase inhibitor, clade a, member 3k, Alb - albumin, Fcer1g - fc receptor, ige, high affinity i, gamma polypeptide, Ctsa - cathepsin a, Cpe - carboxypeptidase e, Kng1 - kininogen 1, Itgb1 - integrin beta 1 (fibronectin receptor beta), Itgam - integrin alpha m, Vim - vimentin, Itga6 - integrin alpha 6, Fabp7 - fatty acid binding protein 7, brain, F2 - coagulation factor ii, Fabp5 - fatty acid binding protein 5, epidermal, Vtn - vitronectin, Itgb3 - integrin beta 3, Tpm1 - tropomyosin 1, alpha, Gba - glucosidase, beta, acid, Itih3 - inter-alpha trypsin inhibitor, heavy chain 3, Serpinb1a - serine (or cysteine) peptidase inhibitor, clade b, member 1a, Itih2 - inter-alpha trypsin inhibitor, heavy chain 2, Itih1 - inter-alpha trypsin inhibitor, heavy chain 1, Arhgdib - rho, gdp dissociation inhibitor (gdi) beta, Irs2 - insulin receptor substrate 2, Stat3 - signal transducer and activator of transcription 3, Stat1 - signal transducer and activator of transcription 1, Anxa2 - annexin a2, Serpina6 - serine (or cysteine) peptidase inhibitor, clade a, member 6, Plg - plasminogen, Serpinh1 - serine (or cysteine) peptidase inhibitor, clade h, member 1, Ctsb - cathepsin b, Cst3 - cystatin c, Lamp2 - lysosomal-associated membrane protein 2, Anpep - alanyl (membrane) aminopeptidase, Gjb6 - gap junction protein, beta 6, Lgmn - legumain, Serpinc1 - serine (or cysteine) peptidase inhibitor, clade c (antithrombin), member 1, Fgg - fibrinogen gamma chain, Fhit - fragile histidine triad gene, Sod1 - superoxide dismutase 1, soluble, Fgb - fibrinogen beta chain, Ide - insulin degrading enzyme, Krt1 - keratin 1, H2-D1 - histocompatibility 2, d region locus 1, Lgals1 - lectin, galactose binding, soluble 1, Hp - haptoglobin, C1qc - complement component 1, q subcomponent, c chain, Hrg - histidine-rich glycoprotein, Ddrgk1 - ddrgk domain containing 1, Lsm6 - lsm6 homolog, u6 small nuclear rna associated (s. cerevisiae), Ybx1 - y box protein 1, Sema4a - sema domain, immunoglobulin domain (ig), transmembrane domain (tm) and short cytoplasmic domain, (semaphorin) 4a, Fga - fibrinogen alpha chain, Gfap - glial fibrillary acidic protein, Clu - clusterin, Apoa1 - apolipoprotein a-i, Pdcd5 - programmed cell death 5, Commd1 - comm domain containing 1, Mug1 - murinoglobulin 1] |
| GO:0050867 | positive regulation of cell activation | 1.84E-5 | 1.86E-3 | 4,65 | 4935 | 73 | 160 | 11 | [Igkc - immunoglobulin kappa constant, Lgals1 - lectin, galactose binding, soluble 1, Itgam - integrin alpha m, Ighg2b - immunoglobulin heavy constant gamma 2b, Ighg1 - immunoglobulin heavy constant gamma 1 (g1m marker), Aif1 - allograft inflammatory factor 1, Ighm - immunoglobulin heavy constant mu, Fcer1g - fc receptor, ige, high affinity i, gamma polypeptide, Irs2 - insulin receptor substrate 2, Lamp1 - lysosomal-associated membrane protein 1, Itgb2 - integrin beta 2] |
| GO:0051240 | positive regulation of multicellular organismal process | 2.11E-5 | 2.1E-3 | 2,00 | 4935 | 572 | 160 | 37 | [Plg - plasminogen, Anxa3 - annexin a3, Ahsg - alpha-2-hs-glycoprotein, Fn1 - fibronectin 1, Clic1 - chloride intracellular channel 1, Aif1 - allograft inflammatory factor 1, Flna - filamin, alpha, Apoh - apolipoprotein h, Lgmn - legumain, Fgg - fibrinogen gamma chain, Alb - albumin, Fcer1g - fc receptor, ige, high affinity i, gamma polypeptide, Sod1 - superoxide dismutase 1, soluble, Fgb - fibrinogen beta chain, Glipr2 - gli pathogenesis-related 2, C3 - complement component 3, Itgb1 - integrin beta 1 (fibronectin receptor beta), Itgam - integrin alpha m, Vim - vimentin, Itga6 - integrin alpha 6, F2 - coagulation factor ii, Fabp5 - fatty acid binding protein 5, epidermal, Lingo2 - leucine rich repeat and ig domain containing 2, Hrg - histidine-rich glycoprotein, Itgb3 - integrin beta 3, Tpm1 - tropomyosin 1, alpha, Itgb2 - integrin beta 2, Gba - glucosidase, beta, acid, Sema4a - sema domain, immunoglobulin domain (ig), transmembrane domain (tm) and short cytoplasmic domain, (semaphorin) 4a, Scarb2 - scavenger receptor class b, member 2, Gfap - glial fibrillary acidic protein, Fga - fibrinogen alpha chain, Clu - clusterin, Stat3 - signal transducer and activator of transcription 3, Stat1 - signal transducer and activator of transcription 1, Scn3b - sodium channel, voltage-gated, type iii, beta, Anxa2 - annexin a2] |
| GO:0002706 | regulation of lymphocyte mediated immunity | 2.2E-5 | 2.18E-3 | 7,71 | 4935 | 28 | 160 | 7 | [Hpx - hemopexin, C3 - complement component 3, H2-D1 - histocompatibility 2, d region locus 1, Ighg2b - immunoglobulin heavy constant gamma 2b, Ighg1 - immunoglobulin heavy constant gamma 1 (g1m marker), Fcer1g - fc receptor, ige, high affinity i, gamma polypeptide, Lamp1 - lysosomal-associated membrane protein 1] |
| GO:1904036 | negative regulation of epithelial cell apoptotic process | 2.22E-5 | 2.17E-3 | 12,85 | 4935 | 12 | 160 | 5 | [Wfs1 - wolfram syndrome 1 homolog (human), Scg2 - secretogranin ii, Fgg - fibrinogen gamma chain, Fga - fibrinogen alpha chain, Fgb - fibrinogen beta chain] |

| | | | | | | | | | |
|---|---|---|---|---|---|---|---|---|---|
| GO:0048523 | negative regulation of cellular process | 2.49E-5 | 2.42E-3 | 1,53 | 4935 | 1414 | 160 | 70 | [Pzp - pregnancy zone protein, Wfs1 - wolfram syndrome 1 homolog (human), Serpina1b - serine (or cysteine) preptidase inhibitor, clade a, member 1b, Scg2 - secretogranin ii, Fn1 - fibronectin 1, Serpina1a - serine (or cysteine) peptidase inhibitor, clade a, member 1a, Ahsg - alpha-2-hs-glycoprotein, Aif1 - allograft inflammatory factor 1, Acot1 - acyl-coa thioesterase 1, Gsn - gelsolin, Nes - nestin, Serpina1d - serine (or cysteine) peptidase inhibitor, clade a, member 1d, Serpina1e - serine (or cysteine) peptidase inhibitor, clade a, member 1e, Flna - filamin, alpha, Apoh - apolipoprotein h, Apod - apolipoprotein d, Psmg2 - proteasome (prosome, macropain) assembly chaperone 2, Serpina3k - serine (or cysteine) peptidase inhibitor, clade a, member 3k, Ppp1r1b - protein phosphatase 1, regulatory (inhibitor) subunit 1b, Fcer1g - fc receptor, ige, high affinity i, gamma polypeptide, Alb - albumin, Ctsa - cathepsin a, Kng1 - kininogen 1, Itgb1 - integrin beta 1 (fibronectin receptor beta), Vim - vimentin, Itgam - integrin alpha m, F2 - coagulation factor ii, Itga6 - integrin alpha 6, Vtn - vitronectin, Itgb3 - integrin beta 3, Tpm1 - tropomyosin 1, alpha, Gba - glucosidase, beta, acid, Serpinb1a - serine (or cysteine) peptidase inhibitor, clade b, member 1a, Itih3 - inter-alpha trypsin inhibitor, heavy chain 3, Itih2 - inter-alpha trypsin inhibitor, heavy chain 2, Arhgdib - rho, gdp dissociation inhibitor (gdi) beta, Itih1 - inter-alpha trypsin inhibitor, heavy chain 1, Irs2 - insulin receptor substrate 2, Stat3 - signal transducer and activator of transcription 3, Stat1 - signal transducer and activator of transcription 1, Anxa2 - annexin a2, Serpina6 - serine (or cysteine) peptidase inhibitor, clade a, member 6, Plg - plasminogen, Serpinh1 - serine (or cysteine) peptidase inhibitor, clade h, member 1, Ctsb - cathepsin b, Cst3 - cystatin c, Lamp2 - lysosomal-associated membrane protein 2, Gjb6 - gap junction protein, beta 6, Lgmn - legumain, Serpinc1 - serine (or cysteine) peptidase inhibitor, clade c (antithrombin), member 1, Fgg - fibrinogen gamma chain, Fhit - fragile histidine triad gene, Sod1 - superoxide dismutase 1, soluble, Fgb - fibrinogen beta chain, Ide - insulin degrading enzyme, H2-D1 - histocompatibility 2, d region locus 1, Lgals1 - lectin, galactose binding, soluble 1, Hp - haptoglobin, C1qc - complement component 1, q subcomponent, c chain, Hrg - histidine-rich glycoprotein, Ddrgk1 - ddrgk domain containing 1, Ybx1 - y box protein 1, Sema4a - sema domain, immunoglobulin domain (ig), transmembrane domain (tm) and short cytoplasmic domain, (semaphorin) 4a, Gfap - glial fibrillary acidic protein, Fga - fibrinogen alpha chain, Pdcd5 - programmed cell death 5, Apoa1 - apolipoprotein a-i, Clu - clusterin, Commd1 - comm domain containing 1, Mug1 - murinoglobulin 1] |
| GO:0010941 | regulation of cell death | 2.67E-5 | 2.57E-3 | 1,97 | 4935 | 578 | 160 | 37 | [Wfs1 - wolfram syndrome 1 homolog (human), Scg2 - secretogranin ii, Fn1 - fibronectin 1, Aif1 - allograft inflammatory factor 1, Ctsb - cathepsin b, Gsn - gelsolin, Acot1 - acyl-coa thioesterase 1, Cst3 - cystatin c, Nes - nestin, Ier3ip1 - immediate early response 3 interacting protein 1, Flna - filamin, alpha, Apoh - apolipoprotein h, Lgmn - legumain, Psmg2 - proteasome (prosome, macropain) assembly chaperone 2, P4hb - prolyl 4-hydroxylase, beta polypeptide, Fgg - fibrinogen gamma chain, Alb - albumin, Fcer1g - fc receptor, ige, high affinity i, gamma polypeptide, Ctsd - cathepsin d, Sod1 - superoxide dismutase 1, soluble, Fgb - fibrinogen beta chain, Itgb1 - integrin beta 1 (fibronectin receptor beta), Itgam - integrin alpha m, Hp - haptoglobin, Itga6 - integrin alpha 6, C1qa - complement component 1, q subcomponent, alpha polypeptide, Ddrgk1 - ddrgk domain containing 1, Itgb3 - integrin beta 3, Gba - glucosidase, beta, acid, Ybx1 - y box protein 1, Tgm2 - transglutaminase 2, c polypeptide, Fga - fibrinogen alpha chain, Clu - clusterin, Pdcd5 - programmed cell death 5, Irs2 - insulin receptor substrate 2, Stat3 - signal transducer and activator of transcription 3, Stat1 - signal transducer and activator of transcription 1] |
| GO:0002821 | positive regulation of adaptive immune response | 2.81E-5 | 2.69E-3 | 9,25 | 4935 | 20 | 160 | 6 | [Hpx - hemopexin, C3 - complement component 3, H2-D1 - histocompatibility 2, d region locus 1, Ighg2b - immunoglobulin heavy constant gamma 2b, Ighg1 - immunoglobulin heavy constant gamma 1 (g1m marker), Fcer1g - fc receptor, ige, high affinity i, gamma polypeptide] |
| GO:0050766 | positive regulation of phagocytosis | 2.82E-5 | 2.68E-3 | 7,45 | 4935 | 29 | 160 | 7 | [C3 - complement component 3, Ahsg - alpha-2-hs-glycoprotein, Ighg2b - immunoglobulin heavy constant gamma 2b, Ighg1 - immunoglobulin heavy constant gamma 1 (g1m marker), Fcer1g - fc receptor, ige, high affinity i, gamma polypeptide, Apoa1 - apolipoprotein a-i, Sod1 - superoxide dismutase 1, soluble] |
| GO:0001812 | positive regulation of type I hypersensitivity | 3.35E-5 | 3.15E-3 | 30,84 | 4935 | 3 | 160 | 3 | [Ighg2b - immunoglobulin heavy constant gamma 2b, Ighg1 - immunoglobulin heavy constant gamma 1 (g1m marker), Fcer1g - fc receptor, ige, high affinity i, gamma polypeptide] |
| GO:0001810 | regulation of type I hypersensitivity | 3.35E-5 | 3.12E-3 | 30,84 | 4935 | 3 | 160 | 3 | [Ighg2b - immunoglobulin heavy constant gamma 2b, Ighg1 - immunoglobulin heavy constant gamma 1 (g1m marker), Fcer1g - fc receptor, ige, high affinity i, gamma polypeptide] |
| GO:0150064 | vertebrate eye-specific patterning | 3.35E-5 | 3.1E-3 | 30,84 | 4935 | 3 | 160 | 3 | [C3 - complement component 3, Itgam - integrin alpha m, C1qa - complement component 1, q subcomponent, alpha polypeptide] |
| GO:0150062 | complement-mediated synapse pruning | 3.35E-5 | 3.07E-3 | 30,84 | 4935 | 3 | 160 | 3 | [C3 - complement component 3, Itgam - integrin alpha m, C1qa - complement component 1, q subcomponent, alpha polypeptide] |
| GO:0006910 | phagocytosis, recognition | 3.45E-5 | 3.14E-3 | 17,62 | 4935 | 7 | 160 | 4 | [Igkc - immunoglobulin kappa constant, Ighg2b - immunoglobulin heavy constant gamma 2b, Ighg1 - immunoglobulin heavy constant gamma 1 (g1m marker), Ighm - immunoglobulin heavy constant mu] |

| | | | | | | | | | |
|---|---|---|---|---|---|---|---|---|---|
| GO:0002675 | positive regulation of acute inflammatory response | 3.45E-5 | 3.12E-3 | 17,62 | 4935 | 7 | 160 | 4 | [C3 - complement component 3, Ighg2b - immunoglobulin heavy constant gamma 2b, Ighg1 - immunoglobulin heavy constant gamma 1 (g1m marker), Fcer1g - fc receptor, ige, high affinity i, gamma polypeptide] |
| GO:2000352 | negative regulation of endothelial cell apoptotic process | 3.45E-5 | 3.09E-3 | 17,62 | 4935 | 7 | 160 | 4 | [Scg2 - secretogranin ii, Fga - fibrinogen alpha chain, Fgg - fibrinogen gamma chain, Fgb - fibrinogen beta chain] |
| GO:0002863 | positive regulation of inflammatory response to antigenic stimulus | 3.45E-5 | 3.07E-3 | 17,62 | 4935 | 7 | 160 | 4 | [C3 - complement component 3, Ighg2b - immunoglobulin heavy constant gamma 2b, Ighg1 - immunoglobulin heavy constant gamma 1 (g1m marker), Fcer1g - fc receptor, ige, high affinity i, gamma polypeptide] |
| GO:0002699 | positive regulation of immune effector process | 4.12E-5 | 3.63E-3 | 5,24 | 4935 | 53 | 160 | 9 | [Hpx - hemopexin, C3 - complement component 3, H2-D1 - histocompatibility 2, d region locus 1, Itgam - integrin alpha m, Ighg2b - immunoglobulin heavy constant gamma 2b, Ighg1 - immunoglobulin heavy constant gamma 1 (g1m marker), Fcer1g - fc receptor, ige, high affinity i, gamma polypeptide, Lamp1 - lysosomal-associated membrane protein 1, Itgb2 - integrin beta 2] |
| GO:0040017 | positive regulation of locomotion | 4.2E-5 | 3.68E-3 | 2,78 | 4935 | 211 | 160 | 19 | [Glipr2 - gli pathogenesis-related 2, Itgb1 - integrin beta 1 (fibronectin receptor beta), Scg2 - secretogranin ii, Anxa3 - annexin a3, Plg - plasminogen, Fn1 - fibronectin 1, Aif1 - allograft inflammatory factor 1, Itga6 - integrin alpha 6, Ddrgk1 - ddrgk domain containing 1, Vtn - vitronectin, Itgb3 - integrin beta 3, Flna - filamin, alpha, Sema4a - sema domain, immunoglobulin domain (ig), transmembrane domain (tm) and short cytoplasmic domain, (semaphorin) 4a, Mcam - melanoma cell adhesion molecule, Lgmn - legumain, Fga - fibrinogen alpha chain, Irs2 - insulin receptor substrate 2, Stat3 - signal transducer and activator of transcription 3, Cd151 - cd151 antigen] |
| GO:0045765 | regulation of angiogenesis | 4.46E-5 | 3.88E-3 | 4,24 | 4935 | 80 | 160 | 11 | [C3 - complement component 3, Sema4a - sema domain, immunoglobulin domain (ig), transmembrane domain (tm) and short cytoplasmic domain, (semaphorin) 4a, Anxa3 - annexin a3, Plg - plasminogen, Itgb1 - integrin beta 1 (fibronectin receptor beta), Apoh - apolipoprotein h, Hrg - histidine-rich glycoprotein, Stat3 - signal transducer and activator of transcription 3, Itgb3 - integrin beta 3, Stat1 - signal transducer and activator of transcription 1, Itgb2 - integrin beta 2] |
| GO:0030335 | positive regulation of cell migration | 4.56E-5 | 3.93E-3 | 2,86 | 4935 | 194 | 160 | 18 | [Glipr2 - gli pathogenesis-related 2, Itgb1 - integrin beta 1 (fibronectin receptor beta), Anxa3 - annexin a3, Plg - plasminogen, Fn1 - fibronectin 1, Aif1 - allograft inflammatory factor 1, Itga6 - integrin alpha 6, Ddrgk1 - ddrgk domain containing 1, Vtn - vitronectin, Itgb3 - integrin beta 3, Flna - filamin, alpha, Sema4a - sema domain, immunoglobulin domain (ig), transmembrane domain (tm) and short cytoplasmic domain, (semaphorin) 4a, Mcam - melanoma cell adhesion molecule, Lgmn - legumain, Fga - fibrinogen alpha chain, Irs2 - insulin receptor substrate 2, Stat3 - signal transducer and activator of transcription 3, Cd151 - cd151 antigen] |
| GO:1904035 | regulation of epithelial cell apoptotic process | 5.13E-5 | 4.39E-3 | 8,41 | 4935 | 22 | 160 | 6 | [Wfs1 - wolfram syndrome 1 homolog (human), Scg2 - secretogranin ii, Fgg - fibrinogen gamma chain, Fga - fibrinogen alpha chain, Gsn - gelsolin, Fgb - fibrinogen beta chain] |
| GO:0002886 | regulation of myeloid leukocyte mediated immunity | 5.13E-5 | 4.35E-3 | 8,41 | 4935 | 22 | 160 | 6 | [C3 - complement component 3, Itgam - integrin alpha m, Ighg2b - immunoglobulin heavy constant gamma 2b, Ighg1 - immunoglobulin heavy constant gamma 1 (g1m marker), Fcer1g - fc receptor, ige, high affinity i, gamma polypeptide, Itgb2 - integrin beta 2] |
| GO:0019730 | antimicrobial humoral response | 5.32E-5 | 4.48E-3 | 11,02 | 4935 | 14 | 160 | 5 | [Fga - fibrinogen alpha chain, Ighg1 - immunoglobulin heavy constant gamma 1 (g1m marker), F2 - coagulation factor ii, Hrg - histidine-rich glycoprotein, Fgb - fibrinogen beta chain] |
| GO:0060548 | negative regulation of cell death | 5.59E-5 | 4.67E-3 | 2,22 | 4935 | 375 | 160 | 27 | [Wfs1 - wolfram syndrome 1 homolog (human), Scg2 - secretogranin ii, Fn1 - fibronectin 1, Aif1 - allograft inflammatory factor 1, Ctsb - cathepsin b, Acot1 - acyl-coa thioesterase 1, Nes - nestin, Cst3 - cystatin c, Flna - filamin, alpha, Apoh - apolipoprotein h, Psmg2 - proteasome (prosome, macropain) assembly chaperone 2, Lgmn - legumain, Fgg - fibrinogen gamma chain, Alb - albumin, Fcer1g - fc receptor, ige, high affinity i, gamma polypeptide, Sod1 - superoxide dismutase 1, soluble, Fgb - fibrinogen beta chain, Itgb1 - integrin beta 1 (fibronectin receptor beta), Itga6 - integrin alpha 6, Ddrgk1 - ddrgk domain containing 1, Itgb3 - integrin beta 3, Gba - glucosidase, beta, acid, Ybx1 - y box protein 1, Fga - fibrinogen alpha chain, Irs2 - insulin receptor substrate 2, Clu - clusterin, Stat3 - signal transducer and activator of transcription 3] |

| | | | | | | | | | |
|---|---|---|---|---|---|---|---|---|---|
| GO:2000147 | positive regulation of cell motility | 5.59E-5 | 4.65E-3 | 2,82 | 4935 | 197 | 160 | 18 | [Glipr2 - gli pathogenesis-related 2, Itgb1 - integrin beta 1 (fibronectin receptor beta), Anxa3 - annexin a3, Plg - plasminogen, Fn1 - fibronectin 1, Aif1 - allograft inflammatory factor 1, Itga6 - integrin alpha 6, Ddrgk1 - ddrgk domain containing 1, Vtn - vitronectin, Itgb3 - integrin beta 3, Flna - filamin, alpha, Sema4a - sema domain, immunoglobulin domain (ig), transmembrane domain (tm) and short cytoplasmic domain, (semaphorin) 4a, Mcam - melanoma cell adhesion molecule, Lgmn - legumain, Fga - fibrinogen alpha chain, Irs2 - insulin receptor substrate 2, Stat3 - signal transducer and activator of transcription 3, Cd151 - cd151 antigen] |
| GO:0007229 | integrin-mediated signaling pathway | 5.61E-5 | 4.63E-3 | 6,75 | 4935 | 32 | 160 | 7 | [Itgb1 - integrin beta 1 (fibronectin receptor beta), Fn1 - fibronectin 1, Itga6 - integrin alpha 6, Fcer1g - fc receptor, ige, high affinity i, gamma polypeptide, Apoa1 - apolipoprotein a-i, Itgb3 - integrin beta 3, Itgb2 - integrin beta 2] |
| GO:0051239 | regulation of multicellular organismal process | 6.3E-5 | 5.15E-3 | 1,64 | 4935 | 995 | 160 | 53 | [Ahsg - alpha-2-hs-glycoprotein, Fn1 - fibronectin 1, Clic1 - chloride intracellular channel 1, Aif1 - allograft inflammatory factor 1, Flna - filamin, alpha, Apoh - apolipoprotein h, Apod - apolipoprotein d, Alb - albumin, Fcer1g - fc receptor, ige, high affinity i, gamma polypeptide, Cpe - carboxypeptidase e, Kng1 - kininogen 1, Glipr2 - gli pathogenesis-related 2, Itgb1 - integrin beta 1 (fibronectin receptor beta), Vim - vimentin, Itgam - integrin alpha m, Itga6 - integrin alpha 6, F2 - coagulation factor ii, Fabp5 - fatty acid binding protein 5, epidermal, Itgb3 - integrin beta 3, Tpm1 - tropomyosin 1, alpha, Rbp1 - retinol binding protein 1, cellular, Itgb2 - integrin beta 2, Gba - glucosidase, beta, acid, Scarb2 - scavenger receptor class b, member 2, Serpinb1a - serine (or cysteine) peptidase inhibitor, clade b, member 1a, Arhgdib - rho, gdp dissociation inhibitor (gdi) beta, Stat3 - signal transducer and activator of transcription 3, Nptxr - neuronal pentraxin receptor, Stat1 - signal transducer and activator of transcription 1, Anxa2 - annexin a2, Scn3b - sodium channel, voltage-gated, type iii, beta, Plg - plasminogen, Anxa3 - annexin a3, Anpep - alanyl (membrane) aminopeptidase, Lgmn - legumain, Serpinc1 - serine (or cysteine) peptidase inhibitor, clade c (antithrombin), member 1, Fgg - fibrinogen gamma chain, Gaa - glucosidase, alpha, acid, Sod1 - superoxide dismutase 1, soluble, Fgb - fibrinogen beta chain, C3 - complement component 3, H2-D1 - histocompatibility 2, d region locus 1, Lgals1 - lectin, galactose binding, soluble 1, C1qc - complement component 1, q subcomponent, c chain, Lingo2 - leucine rich repeat and ig domain containing 2, Hrg - histidine-rich glycoprotein, Kcnj10 - potassium inwardly-rectifying channel, subfamily j, member 10, Sema4a - sema domain, immunoglobulin domain (ig), transmembrane domain (tm) and short cytoplasmic domain, (semaphorin) 4a, Gfap - glial fibrillary acidic protein, Fga - fibrinogen alpha chain, Apoa1 - apolipoprotein a-i, Clu - clusterin, Nptx2 - neuronal pentraxin 2] |
| GO:0050865 | regulation of cell activation | 6.71E-5 | 5.46E-3 | 3,32 | 4935 | 130 | 160 | 14 | [Igkc - immunoglobulin kappa constant, Lgals1 - lectin, galactose binding, soluble 1, Itgam - integrin alpha m, Aif1 - allograft inflammatory factor 1, Gsn - gelsolin, Hrg - histidine-rich glycoprotein, Lamp1 - lysosomal-associated membrane protein 1, Itgb2 - integrin beta 2, Ighg2b - immunoglobulin heavy constant gamma 2b, Fgg - fibrinogen gamma chain, Ighg1 - immunoglobulin heavy constant gamma 1 (g1m marker), Fcer1g - fc receptor, ige, high affinity i, gamma polypeptide, Ighm - immunoglobulin heavy constant mu, Irs2 - insulin receptor substrate 2] |
| GO:0002269 | leukocyte activation involved in inflammatory response | 6.73E-5 | 5.43E-3 | 15,42 | 4935 | 8 | 160 | 4 | [Itgam - integrin alpha m, Aif1 - allograft inflammatory factor 1, Clu - clusterin, C1qa - complement component 1, q subcomponent, alpha polypeptide] |
| GO:0001774 | microglial cell activation | 6.73E-5 | 5.39E-3 | 15,42 | 4935 | 8 | 160 | 4 | [Itgam - integrin alpha m, Aif1 - allograft inflammatory factor 1, Clu - clusterin, C1qa - complement component 1, q subcomponent, alpha polypeptide] |
| GO:0043062 | extracellular structure organization | 7.18E-5 | 5.71E-3 | 4,41 | 4935 | 70 | 160 | 10 | [Plg - plasminogen, Itgb1 - integrin beta 1 (fibronectin receptor beta), Fn1 - fibronectin 1, Gfap - glial fibrillary acidic protein, Serpinh1 - serine (or cysteine) peptidase inhibitor, clade h, member 1, Itih1 - inter-alpha trypsin inhibitor, heavy chain 1, Apoa1 - apolipoprotein a-i, Lcp1 - lymphocyte cytosolic protein 1, Vtn - vitronectin, Anxa2 - annexin a2] |
| GO:0030198 | extracellular matrix organization | 8.61E-5 | 6.8E-3 | 4,79 | 4935 | 58 | 160 | 9 | [Plg - plasminogen, Itgb1 - integrin beta 1 (fibronectin receptor beta), Fn1 - fibronectin 1, Gfap - glial fibrillary acidic protein, Serpinh1 - serine (or cysteine) peptidase inhibitor, clade h, member 1, Itih1 - inter-alpha trypsin inhibitor, heavy chain 1, Lcp1 - lymphocyte cytosolic protein 1, Vtn - vitronectin, Anxa2 - annexin a2] |

| | | | | | | | | | |
|---|---|---|---|---|---|---|---|---|---|
| GO:0050794 | regulation of cellular process | 9,00E-05 | 7.06E-3 | 1,25 | 4935 | 2835 | 160 | 115 | [illegible] apolipoprotein d, Psmg2 - proteasome (prosome, macropain) assembly chaperone 2, Fcer1g - fc receptor, ige, high affinity i, gamma polypeptide, Kng1 - kininogen 1, F2 - coagulation factor ii, Arhgap22 - rho gtpase activating protein 22, Fabp5 - fatty acid binding protein 5, epidermal, Psme1 - proteasome (prosome, macropain) activator subunit 1 (pa28 alpha), Sh3bgrl3 - sh3 domain binding glutamic acid-rich protein-like 3, Gba - glucosidase, beta, acid, Arhgdib - rho, gdp dissociation inhibitor (gdi) beta, Irs2 - insulin receptor substrate 2, Stat3 - signal transducer and activator of transcription 3, Nptxr - neuronal pentraxin receptor, Arpc1b - actin related protein 2/3 complex, subunit 1b, Stat1 - signal transducer and activator of transcription 1, Anxa2 - annexin a2, Plg - plasminogen, Serpina6 - serine (or cysteine) peptidase inhibitor, clade a, member 6, Serpinh1 - serine (or cysteine) peptidase inhibitor, clade h, member 1, Ctsb - cathepsin b, Lcp1 - lymphocyte cytosolic protein 1, Cst3 - cystatin c, Serpinc1 - serine (or cysteine) peptidase inhibitor, clade c (antithrombin), member 1, Irgm1 - immunity-related gtpase family m member 1, Fgg - fibrinogen gamma chain, Ctsd - cathepsin d, Fgb - fibrinogen beta chain, Ide - insulin degrading enzyme, S100a13 - s100 calcium binding protein a13, Hrg - histidine-rich glycoprotein, Hexb - hexosaminidase b, Ncstn - nicastrin, Clu - clusterin, Serpina1b - serine (or cysteine) preptidase inhibitor, clade a, member 1b, Wfs1 - wolfram syndrome 1 homolog (human), Scg2 - secretogranin ii, Serpina1a - serine (or cysteine) peptidase inhibitor, clade a, member 1a, Ahsg - alpha-2-hs-glycoprotein, Fn1 - fibronectin 1, Clic1 - chloride intracellular channel 1, Aif1 - allograft inflammatory factor 1, Acot1 - acyl-coa thioesterase 1, Nes - nestin, Serpina1d - serine (or cysteine) peptidase inhibitor, clade a, member 1d, Serpina1e - serine (or cysteine) peptidase inhibitor, clade a, member 1e, Mcam - melanoma cell adhesion molecule, P4hb - prolyl 4-hydroxylase, beta polypeptide, Ppp1r1b - protein phosphatase 1, regulatory (inhibitor) subunit 1b, Serpina3k - serine (or cysteine) peptidase inhibitor, clade a, member 3k, Crip2 - cysteine rich protein 2, Alb - albumin, Ctsa - cathepsin a, Marcksl1 - marcks-like 1, Cpe - carboxypeptidase e, Glipr2 - gli pathogenesis-related 2, Igkc - immunoglobulin kappa constant, Itgb1 - integrin beta 1 (fibronectin receptor beta), Vim - vimentin, Itgam - integrin alpha m, Itga6 - integrin alpha 6, Vtn - vitronectin, Itgb3 - integrin beta 3, Tpm1 - tropomyosin 1, alpha, Rbp1 - retinol binding protein 1, cellular, Itgb2 - integrin beta 2, Scarb2 - scavenger receptor class b, member 2, Serpinb1a - serine (or cysteine) peptidase inhibitor, clade b, member 1a, Ighg2b - immunoglobulin heavy constant gamma 2b, Itih3 - inter-alpha trypsin inhibitor, heavy chain 3, Ighg1 - immunoglobulin heavy constant gamma 1 (g1m marker), Itih2 - inter-alpha trypsin inhibitor, heavy chain 2, Itih1 - inter-alpha trypsin inhibitor, heavy chain 1, Ighm - immunoglobulin heavy constant mu, Cd151 - cd151 antigen, Scn3b - sodium channel, voltage-gated, type iii, beta, Anxa3 - annexin a3, Ier3ip1 - immediate early response 3 interacting protein 1, Lamp2 - lysosomal-associated membrane protein 2, Lamp1 - lysosomal-associated membrane protein 1, Anpep - alanyl (membrane) aminopeptidase, Gjb6 - gap junction protein, beta 6, Lgmn - legumain, Fhit - fragile histidine triad gene, Sod1 - superoxide dismutase 1, soluble, C3 - complement component 3, H2-D1 - histocompatibility 2, d region locus 1, Lgals1 - lectin, galactose binding, soluble 1, Ppp1r1a - protein phosphatase 1, regulatory (inhibitor) subunit 1a, Hp - haptoglobin, C1qc - complement component 1, q subcomponent, c chain, Lingo2 - leucine rich repeat and ig domain containing 2, C1qa - complement component 1, q subcomponent, alpha polypeptide, Kcnj10 - potassium inwardly-rectifying channel, subfamily j, member 10, Ddrgk1 - ddrgk domain containing 1, Tgm1 - transglutaminase 1, k polypeptide, Ybx1 - y box protein 1, Hpx - hemopexin, Sema4a - sema domain, immunoglobulin domain (ig), transmembrane domain (tm) and short cytoplasmic domain, (semaphorin) 4a, Tgm2 - transglutaminase 2, c polypeptide, Fga - fibrinogen alpha chain, Gfap - glial fibrillary acidic protein, Apoa1 - apolipoprotein a-i, Pdcd5 - programmed cell death 5, Nptx2 - neuronal pentraxin 2, Txndc5 - thioredoxin domain containing 5, Commd1 - comm domain containing 1, Mug1 - murinoglobulin 1, Tbl2 - transducin (beta)-like 2] |
| GO:0030334 | regulation of cell migration | 9.32E-5 | 7.26E-3 | 2,29 | 4935 | 323 | 160 | 24 | [Glipr2 - gli pathogenesis-related 2, Itgb1 - integrin beta 1 (fibronectin receptor beta), Plg - plasminogen, Anxa3 - annexin a3, Vim - vimentin, Fn1 - fibronectin 1, Aif1 - allograft inflammatory factor 1, Itga6 - integrin alpha 6, Hrg - histidine-rich glycoprotein, Ddrgk1 - ddrgk domain containing 1, Vtn - vitronectin, Itgb3 - integrin beta 3, Tpm1 - tropomyosin 1, alpha, Flna - filamin, alpha, Sema4a - sema domain, immunoglobulin domain (ig), transmembrane domain (tm) and short cytoplasmic domain, (semaphorin) 4a, Apoh - apolipoprotein h, Mcam - melanoma cell adhesion molecule, Apod - apolipoprotein d, Lgmn - legumain, Fga - fibrinogen alpha chain, Arhgdib - rho, gdp dissociation inhibitor (gdi) beta, Irs2 - insulin receptor substrate 2, Stat3 - signal transducer and activator of transcription 3, Cd151 - cd151 antigen] |
| GO:0051272 | positive regulation of cellular component movement | 9.44E-5 | 7.31E-3 | 2,71 | 4935 | 205 | 160 | 18 | [Glipr2 - gli pathogenesis-related 2, Itgb1 - integrin beta 1 (fibronectin receptor beta), Anxa3 - annexin a3, Plg - plasminogen, Fn1 - fibronectin 1, Aif1 - allograft inflammatory factor 1, Itga6 - integrin alpha 6, Ddrgk1 - ddrgk domain containing 1, Vtn - vitronectin, Itgb3 - integrin beta 3, Flna - filamin, alpha, Sema4a - sema domain, immunoglobulin domain (ig), transmembrane domain (tm) and short cytoplasmic domain, (semaphorin) 4a, Mcam - melanoma cell adhesion molecule, Lgmn - legumain, Fga - fibrinogen alpha chain, Irs2 - insulin receptor substrate 2, Stat3 - signal transducer and activator of transcription 3, Cd151 - cd151 antigen] |
| GO:0001525 | angiogenesis | 1.03E-4 | 7.95E-3 | 4,23 | 4935 | 73 | 160 | 10 | [Flna - filamin, alpha, Sema4a - sema domain, immunoglobulin domain (ig), transmembrane domain (tm) and short cytoplasmic domain, (semaphorin) 4a, Scg2 - secretogranin ii, Mcam - melanoma cell adhesion molecule, Fn1 - fibronectin 1, Cfh - complement component factor h, Arhgap22 - rho gtpase activating protein 22, Hrg - histidine-rich glycoprotein, Anpep - alanyl (membrane) aminopeptidase, Anxa2 - annexin a2] |
| GO:1901342 | regulation of vasculature development | 1.09E-4 | 8.32E-3 | 3,86 | 4935 | 88 | 160 | 11 | [C3 - complement component 3, Sema4a - sema domain, immunoglobulin domain (ig), transmembrane domain (tm) and short cytoplasmic domain, (semaphorin) 4a, Anxa3 - annexin a3, Plg - plasminogen, Itgb1 - integrin beta 1 (fibronectin receptor beta), Apoh - apolipoprotein h, Hrg - histidine-rich glycoprotein, Stat3 - signal transducer and activator of transcription 3, Itgb3 - integrin beta 3, Stat1 - signal transducer and activator of transcription 1, Itgb2 - integrin beta 2] |
| GO:0002673 | regulation of acute inflammatory response | 1.1E-4 | 8.35E-3 | 9,64 | 4935 | 16 | 160 | 5 | [C3 - complement component 3, Ighg2b - immunoglobulin heavy constant gamma 2b, Ighg1 - immunoglobulin heavy constant gamma 1 (g1m marker), Fcer1g - fc receptor, ige, high affinity i, gamma polypeptide, Cfh - complement component factor h] |
| GO:0050871 | positive regulation of B cell activation | 1.1E-4 | 8.29E-3 | 9,64 | 4935 | 16 | 160 | 5 | [Igkc - immunoglobulin kappa constant, Ighg2b - immunoglobulin heavy constant gamma 2b, Ighg1 - immunoglobulin heavy constant gamma 1 (g1m marker), Ighm - immunoglobulin heavy constant mu, Irs2 - insulin receptor substrate 2] |

| | | | | | | | | | |
|---|---|---|---|---|---|---|---|---|---|
| GO:0034109 | homotypic cell-cell adhesion | 1.12E-4 | 8.4E-3 | 7,40 | 4935 | 25 | 160 | 6 | [Fn1 - fibronectin 1, Fga - fibrinogen alpha chain, Fgg - fibrinogen gamma chain, Cfh - complement component factor h, Itgb3 - integrin beta 3, Fgb - fibrinogen beta chain] |
| GO:0002822 | regulation of adaptive immune response based on somatic recombination of immune receptors built from immunoglobulin superfamily domains | 1.12E-4 | 8.35E-3 | 7,40 | 4935 | 25 | 160 | 6 | [Hpx - hemopexin, C3 - complement component 3, H2-D1 - histocompatibility 2, d region locus 1, Ighg2b - immunoglobulin heavy constant gamma 2b, Ighg1 - immunoglobulin heavy constant gamma 1 (g1m marker), Fcer1g - fc receptor, ige, high affinity i, gamma polypeptide] |
| GO:0002455 | humoral immune response mediated by circulating immunoglobulin | 1.31E-4 | 9.65E-3 | 23,13 | 4935 | 4 | 160 | 3 | [Ighg2b - immunoglobulin heavy constant gamma 2b, Ighg1 - immunoglobulin heavy constant gamma 1 (g1m marker), Ighm - immunoglobulin heavy constant mu] |
| GO:0030155 | regulation of cell adhesion | 1.36E-4 | 9.98E-3 | 2,55 | 4935 | 230 | 160 | 19 | [Itgb1 - integrin beta 1 (fibronectin receptor beta), Plg - plasminogen, Lgals1 - lectin, galactose binding, soluble 1, Fn1 - fibronectin 1, Aif1 - allograft inflammatory factor 1, Itga6 - integrin alpha 6, Gsn - gelsolin, Vtn - vitronectin, Itgb3 - integrin beta 3, Tpm1 - tropomyosin 1, alpha, Itgb2 - integrin beta 2, Flna - filamin, alpha, Tgm2 - transglutaminase 2, c polypeptide, Apod - apolipoprotein d, P4hb - prolyl 4-hydroxylase, beta polypeptide, Fgg - fibrinogen gamma chain, Fga - fibrinogen alpha chain, Apoa1 - apolipoprotein a-i, Fgb - fibrinogen beta chain] |
| GO:0043066 | negative regulation of apoptotic process | 1.42E-4 | 1.04E-2 | 2,28 | 4935 | 311 | 160 | 23 | [Wfs1 - wolfram syndrome 1 homolog (human), Itgb1 - integrin beta 1 (fibronectin receptor beta), Scg2 - secretogranin ii, Fn1 - fibronectin 1, Aif1 - allograft inflammatory factor 1, Itga6 - integrin alpha 6, Acot1 - acyl-coa thioesterase 1, Nes - nestin, Ddrgk1 - ddrgk domain containing 1, Gba - glucosidase, beta, acid, Ybx1 - y box protein 1, Flna - filamin, alpha, Apoh - apolipoprotein h, Lgmn - legumain, Psmg2 - proteasome (prosome, macropain) assembly chaperone 2, Fgg - fibrinogen gamma chain, Fga - fibrinogen alpha chain, Fcer1g - fc receptor, ige, high affinity i, gamma polypeptide, Alb - albumin, Clu - clusterin, Irs2 - insulin receptor substrate 2, Sod1 - superoxide dismutase 1, soluble, Fgb - fibrinogen beta chain] |
| GO:0032269 | negative regulation of cellular protein metabolic process | 1.47E-4 | 1.06E-2 | 2,14 | 4935 | 375 | 160 | 26 | [Pzp - pregnancy zone protein, Serpina1b - serine (or cysteine) preptidase inhibitor, clade a, member 1b, Serpina6 - serine (or cysteine) peptidase inhibitor, clade a, member 6, Ahsg - alpha-2-hs-glycoprotein, Serpina1a - serine (or cysteine) peptidase inhibitor, clade a, member 1a, Serpinh1 - serine (or cysteine) peptidase inhibitor, clade h, member 1, Cst3 - cystatin c, Serpina1d - serine (or cysteine) peptidase inhibitor, clade a, member 1d, Serpina1e - serine (or cysteine) peptidase inhibitor, clade a, member 1e, Serpinc1 - serine (or cysteine) peptidase inhibitor, clade c (antithrombin), member 1, Serpina3k - serine (or cysteine) peptidase inhibitor, clade a, member 3k, Ppp1r1b - protein phosphatase 1, regulatory (inhibitor) subunit 1b, Fhit - fragile histidine triad gene, Kng1 - kininogen 1, Ide - insulin degrading enzyme, F2 - coagulation factor ii, Hrg - histidine-rich glycoprotein, Ddrgk1 - ddrgk domain containing 1, Vtn - vitronectin, Gba - glucosidase, beta, acid, Serpinb1a - serine (or cysteine) peptidase inhibitor, clade b, member 1a, Itih3 - inter-alpha trypsin inhibitor, heavy chain 3, Itih2 - inter-alpha trypsin inhibitor, heavy chain 2, Itih1 - inter-alpha trypsin inhibitor, heavy chain 1, Mug1 - murinoglobulin 1, Anxa2 - annexin a2] |
| GO:0031589 | cell-substrate adhesion | 1.47E-4 | 1.06E-2 | 4,48 | 4935 | 62 | 160 | 9 | [Itgb1 - integrin beta 1 (fibronectin receptor beta), Fn1 - fibronectin 1, Fga - fibrinogen alpha chain, Fgg - fibrinogen gamma chain, Itga6 - integrin alpha 6, Vtn - vitronectin, Itgb3 - integrin beta 3, Fgb - fibrinogen beta chain, Itgb2 - integrin beta 2] |
| GO:0006909 | phagocytosis | 1.5E-4 | 1.07E-2 | 5,84 | 4935 | 37 | 160 | 7 | [Tgm2 - transglutaminase 2, c polypeptide, Itgb1 - integrin beta 1 (fibronectin receptor beta), Anxa3 - annexin a3, Itgam - integrin alpha m, Txndc5 - thioredoxin domain containing 5, Itgb3 - integrin beta 3, Itgb2 - integrin beta 2] |
| GO:2000145 | regulation of cell motility | 1.5E-4 | 1.07E-2 | 2,22 | 4935 | 333 | 160 | 24 | [Glipr2 - gli pathogenesis-related 2, Itgb1 - integrin beta 1 (fibronectin receptor beta), Plg - plasminogen, Anxa3 - annexin a3, Vim - vimentin, Fn1 - fibronectin 1, Aif1 - allograft inflammatory factor 1, Itga6 - integrin alpha 6, Hrg - histidine-rich glycoprotein, Ddrgk1 - ddrgk domain containing 1, Vtn - vitronectin, Itgb3 - integrin beta 3, Tpm1 - tropomyosin 1, alpha, Flna - filamin, alpha, Sema4a - sema domain, immunoglobulin domain (ig), transmembrane domain (tm) and short cytoplasmic domain, (semaphorin) 4a, Apoh - apolipoprotein h, Mcam - melanoma cell adhesion molecule, Apod - apolipoprotein d, Lgmn - legumain, Fga - fibrinogen alpha chain, Arhgdib - rho, gdp dissociation inhibitor (gdi) beta, Irs2 - insulin receptor substrate 2, Stat3 - signal transducer and activator of transcription 3, Cd151 - cd151 antigen] |
| GO:0030168 | platelet activation | 1.52E-4 | 1.07E-2 | 9,07 | 4935 | 17 | 160 | 5 | [Fga - fibrinogen alpha chain, Fgg - fibrinogen gamma chain, F2 - coagulation factor ii, Itgb3 - integrin beta 3, Fgb - fibrinogen beta chain] |

| | | | | | | | | | |
|---|---|---|---|---|---|---|---|---|---|
| GO:0048518 | positive regulation of biological process | 1.77E-4 | 1.24E-2 | 1,38 | 4935 | 1811 | 160 | 81 | [Wfs1 - wolfram syndrome 1 homolog (human), Scg2 - secretogranin ii, Mvd - mevalonate (diphospho) decarboxylase, Ahsg - alpha-2-hs-glycoprotein, Fn1 - fibronectin 1, Clic1 - chloride intracellular channel 1, Aif1 - allograft inflammatory factor 1, Gsn - gelsolin, Nes - nestin, Flna - filamin, alpha, Apoh - apolipoprotein h, Mcam - melanoma cell adhesion molecule, P4hb - prolyl 4-hydroxylase, beta polypeptide, Alb - albumin, Crip2 - cysteine rich protein 2, Fcer1g - fc receptor, ige, high affinity i, gamma polypeptide, Marcksl1 - marcks-like 1, Igkc - immunoglobulin kappa constant, Glipr2 - gli pathogenesis-related 2, Itgb1 - integrin beta 1 (fibronectin receptor beta), Itgam - integrin alpha m, Vim - vimentin, Itga6 - integrin alpha 6, F2 - coagulation factor ii, Fabp5 - fatty acid binding protein 5, epidermal, Psme1 - proteasome (prosome, macropain) activator subunit 1 (pa28 alpha), Vtn - vitronectin, Itgb3 - integrin beta 3, Tpm1 - tropomyosin 1, alpha, Itgb2 - integrin beta 2, Gba - glucosidase, beta, acid, Scarb2 - scavenger receptor class b, member 2, Ighg2b - immunoglobulin heavy constant gamma 2b, Ighg1 - immunoglobulin heavy constant gamma 1 (g1m marker), Ighm - immunoglobulin heavy constant mu, Irs2 - insulin receptor substrate 2, Stat3 - signal transducer and activator of transcription 3, Cd151 - cd151 antigen, Arpc1b - actin related protein 2/3 complex, subunit 1b, Stat1 - signal transducer and activator of transcription 1, Anxa2 - annexin a2, Scn3b - sodium channel, voltage-gated, type iii, beta, Anxa3 - annexin a3, Plg - plasminogen, Lcp1 - lymphocyte cytosolic protein 1, Cst3 - cystatin c, Lamp1 - lysosomal-associated membrane protein 1, Irgm1 - immunity-related gtpase family m member 1, Lgmn - legumain, Fgg - fibrinogen gamma chain, Cfh - complement component factor h, Ctsd - cathepsin d, Sod1 - superoxide dismutase 1, soluble, Fgb - fibrinogen beta chain, Krt1 - keratin 1, C3 - complement component 3, H2-D1 - histocompatibility 2, d region locus 1, Lgals1 - lectin, galactose binding, soluble 1, C4b - complement component 4b (chido blood group), Hp - haptoglobin, Cfb - complement factor b, C1qc - complement component 1, q subcomponent, c chain, Lingo2 - leucine rich repeat and ig domain containing 2, S100a13 - s100 calcium binding protein a13, C1qa - complement component 1, q subcomponent, alpha polypeptide, C1qb - complement component 1, q subcomponent, beta polypeptide, Hrg - histidine-rich glycoprotein, Ddrgk1 - ddrgk domain containing 1, Hexb - hexosaminidase b, Hpx - hemopexin, Ybx1 - y box protein 1, Tgm1 - transglutaminase 1, k polypeptide, Sema4a - sema domain, immunoglobulin domain (ig), transmembrane domain (tm) and short cytoplasmic domain, (semaphorin) 4a, Tgm2 - transglutaminase 2, c polypeptide, Fga - fibrinogen alpha chain, Gfap - glial fibrillary acidic protein, Apoa1 - apolipoprotein a-i, Clu - clusterin, Ncstn - nicastrin, Pdcd5 - programmed cell death 5, Commd1 - comm domain containing 1] |
| GO:0043067 | regulation of programmed cell death | 1.8E-4 | 1.26E-2 | 1,92 | 4935 | 514 | 160 | 32 | [Wfs1 - wolfram syndrome 1 homolog (human), Scg2 - secretogranin ii, Fn1 - fibronectin 1, Aif1 - allograft inflammatory factor 1, Gsn - gelsolin, Acot1 - acyl-coa thioesterase 1, Cst3 - cystatin c, Nes - nestin, Ier3ip1 - immediate early response 3 interacting protein 1, Flna - filamin, alpha, Apoh - apolipoprotein h, Lgmn - legumain, Psmg2 - proteasome (prosome, macropain) assembly chaperone 2, P4hb - prolyl 4-hydroxylase, beta polypeptide, Fgg - fibrinogen gamma chain, Alb - albumin, Fcer1g - fc receptor, ige, high affinity i, gamma polypeptide, Ctsd - cathepsin d, Sod1 - superoxide dismutase 1, soluble, Fgb - fibrinogen beta chain, Itgb1 - integrin beta 1 (fibronectin receptor beta), Itgam - integrin alpha m, Itga6 - integrin alpha 6, Ddrgk1 - ddrgk domain containing 1, Gba - glucosidase, beta, acid, Ybx1 - y box protein 1, Tgm2 - transglutaminase 2, c polypeptide, Fga - fibrinogen alpha chain, Clu - clusterin, Irs2 - insulin receptor substrate 2, Pdcd5 - programmed cell death 5, Stat1 - signal transducer and activator of transcription 1] |
| GO:0018149 | peptide cross-linking | 1.92E-4 | 1.33E-2 | 12,34 | 4935 | 10 | 160 | 4 | [Tgm1 - transglutaminase 1, k polypeptide, Krt1 - keratin 1, Tgm2 - transglutaminase 2, c polypeptide, Fn1 - fibronectin 1] |
| GO:1900048 | positive regulation of hemostasis | 1.92E-4 | 1.32E-2 | 12,34 | 4935 | 10 | 160 | 4 | [Apoh - apolipoprotein h, Plg - plasminogen, F2 - coagulation factor ii, Hrg - histidine-rich glycoprotein] |
| GO:0034114 | regulation of heterotypic cell-cell adhesion | 1.92E-4 | 1.32E-2 | 12,34 | 4935 | 10 | 160 | 4 | [Fgg - fibrinogen gamma chain, Fga - fibrinogen alpha chain, Apoa1 - apolipoprotein a-i, Fgb - fibrinogen beta chain] |
| GO:0030194 | positive regulation of blood coagulation | 1.92E-4 | 1.31E-2 | 12,34 | 4935 | 10 | 160 | 4 | [Plg - plasminogen, Apoh - apolipoprotein h, F2 - coagulation factor ii, Hrg - histidine-rich glycoprotein] |
| GO:2000351 | regulation of endothelial cell apoptotic process | 1.92E-4 | 1.3E-2 | 12,34 | 4935 | 10 | 160 | 4 | [Scg2 - secretogranin ii, Fgg - fibrinogen gamma chain, Fga - fibrinogen alpha chain, Fgb - fibrinogen beta chain] |
| GO:0002861 | regulation of inflammatory response to antigenic stimulus | 1.92E-4 | 1.29E-2 | 12,34 | 4935 | 10 | 160 | 4 | [C3 - complement component 3, Ighg2b - immunoglobulin heavy constant gamma 2b, Ighg1 - immunoglobulin heavy constant gamma 1 (g1m marker), Fcer1g - fc receptor, ige, high affinity i, gamma polypeptide] |
| GO:0046716 | muscle cell cellular homeostasis | 1.92E-4 | 1.28E-2 | 12,34 | 4935 | 10 | 160 | 4 | [Plg - plasminogen, Gaa - glucosidase, alpha, acid, Lamp2 - lysosomal-associated membrane protein 2, Sod1 - superoxide dismutase 1, soluble] |

| | | | | | | | | | |
|---|---|---|---|---|---|---|---|---|---|
| GO:0010810 | regulation of cell-substrate adhesion | 1.99E-4 | 1.32E-2 | 3,61 | 4935 | 94 | 160 | 11 | [Flna - filamin, alpha, Plg - plasminogen, Itgb1 - integrin beta 1 (fibronectin receptor beta), Lgals1 - lectin, galactose binding, soluble 1, Apod - apolipoprotein d, Fn1 - fibronectin 1, P4hb - prolyl 4-hydroxylase, beta polypeptide, Itga6 - integrin alpha 6, Apoa1 - apolipoprotein a-i, Vtn - vitronectin, Itgb3 - integrin beta 3] |
| GO:0042445 | hormone metabolic process | 2.12E-4 | 1.4E-2 | 5,54 | 4935 | 39 | 160 | 7 | [Serpina6 - serine (or cysteine) peptidase inhibitor, clade a, member 6, Ctsb - cathepsin b, Apoa1 - apolipoprotein a-i, Ttr - transthyretin, Cpe - carboxypeptidase e, Rbp1 - retinol binding protein 1, cellular, Ide - insulin degrading enzyme] |
| GO:0002694 | regulation of leukocyte activation | 2.18E-4 | 1.43E-2 | 3,33 | 4935 | 111 | 160 | 12 | [Igkc - immunoglobulin kappa constant, Lgals1 - lectin, galactose binding, soluble 1, Itgam - integrin alpha m, Ighg2b - immunoglobulin heavy constant gamma 2b, Ighg1 - immunoglobulin heavy constant gamma 1 (g1m marker), Aif1 - allograft inflammatory factor 1, Ighm - immunoglobulin heavy constant mu, Fcer1g - fc receptor, ige, high affinity i, gamma polypeptide, Irs2 - insulin receptor substrate 2, Gsn - gelsolin, Lamp1 - lysosomal-associated membrane protein 1, Itgb2 - integrin beta 2] |
| GO:0002274 | myeloid leukocyte activation | 2.2E-4 | 1.44E-2 | 6,61 | 4935 | 28 | 160 | 6 | [Anxa3 - annexin a3, Itgam - integrin alpha m, Aif1 - allograft inflammatory factor 1, Fcer1g - fc receptor, ige, high affinity i, gamma polypeptide, Clu - clusterin, C1qa - complement component 1, q subcomponent, alpha polypeptide] |
| GO:0019882 | antigen processing and presentation | 2.2E-4 | 1.43E-2 | 6,61 | 4935 | 28 | 160 | 6 | [H2-D1 - histocompatibility 2, d region locus 1, Ighm - immunoglobulin heavy constant mu, Fcer1g - fc receptor, ige, high affinity i, gamma polypeptide, Psme1 - proteasome (prosome, macropain) activator subunit 1 (pa28 alpha), Gba - glucosidase, beta, acid, Ide - insulin degrading enzyme] |
| GO:0043069 | negative regulation of programmed cell death | 2.39E-4 | 1.55E-2 | 2,20 | 4935 | 322 | 160 | 23 | [Wfs1 - wolfram syndrome 1 homolog (human), Itgb1 - integrin beta 1 (fibronectin receptor beta), Scg2 - secretogranin ii, Fn1 - fibronectin 1, Aif1 - allograft inflammatory factor 1, Itga6 - integrin alpha 6, Acot1 - acyl-coa thioesterase 1, Nes - nestin, Ddrgk1 - ddrgk domain containing 1, Gba - glucosidase, beta, acid, Ybx1 - y box protein 1, Flna - filamin, alpha, Apoh - apolipoprotein h, Lgmn - legumain, Psmg2 - proteasome (prosome, macropain) assembly chaperone 2, Fgg - fibrinogen gamma chain, Fga - fibrinogen alpha chain, Fcer1g - fc receptor, ige, high affinity i, gamma polypeptide, Alb - albumin, Clu - clusterin, Irs2 - insulin receptor substrate 2, Sod1 - superoxide dismutase 1, soluble, Fgb - fibrinogen beta chain] |
| GO:0010952 | positive regulation of peptidase activity | 2.4E-4 | 1.54E-2 | 4,21 | 4935 | 66 | 160 | 9 | [Lgmn - legumain, Ncstn - nicastrin, Pdcd5 - programmed cell death 5, Gsn - gelsolin, Ctsd - cathepsin d, Psme1 - proteasome (prosome, macropain) activator subunit 1 (pa28 alpha), Ddrgk1 - ddrgk domain containing 1, Stat3 - signal transducer and activator of transcription 3, Stat1 - signal transducer and activator of transcription 1] |
| GO:0040012 | regulation of locomotion | 2.42E-4 | 1.55E-2 | 2,11 | 4935 | 365 | 160 | 25 | [Anxa3 - annexin a3, Scg2 - secretogranin ii, Plg - plasminogen, Fn1 - fibronectin 1, Aif1 - allograft inflammatory factor 1, Flna - filamin, alpha, Apoh - apolipoprotein h, Mcam - melanoma cell adhesion molecule, Apod - apolipoprotein d, Lgmn - legumain, Glipr2 - gli pathogenesis-related 2, Itgb1 - integrin beta 1 (fibronectin receptor beta), Vim - vimentin, Itga6 - integrin alpha 6, Hrg - histidine-rich glycoprotein, Vtn - vitronectin, Ddrgk1 - ddrgk domain containing 1, Itgb3 - integrin beta 3, Tpm1 - tropomyosin 1, alpha, Sema4a - sema domain, immunoglobulin domain (ig), transmembrane domain (tm) and short cytoplasmic domain, (semaphorin) 4a, Fga - fibrinogen alpha chain, Arhgdib - rho, gdp dissociation inhibitor (gdi) beta, Irs2 - insulin receptor substrate 2, Stat3 - signal transducer and activator of transcription 3, Cd151 - cd151 antigen] |
| GO:0042981 | regulation of apoptotic process | 2.5E-4 | 1.59E-2 | 1,91 | 4935 | 500 | 160 | 31 | [Wfs1 - wolfram syndrome 1 homolog (human), Scg2 - secretogranin ii, Fn1 - fibronectin 1, Aif1 - allograft inflammatory factor 1, Gsn - gelsolin, Acot1 - acyl-coa thioesterase 1, Nes - nestin, Ier3ip1 - immediate early response 3 interacting protein 1, Flna - filamin, alpha, Apoh - apolipoprotein h, Psmg2 - proteasome (prosome, macropain) assembly chaperone 2, Lgmn - legumain, P4hb - prolyl 4-hydroxylase, beta polypeptide, Fgg - fibrinogen gamma chain, Alb - albumin, Fcer1g - fc receptor, ige, high affinity i, gamma polypeptide, Ctsd - cathepsin d, Sod1 - superoxide dismutase 1, soluble, Fgb - fibrinogen beta chain, Itgb1 - integrin beta 1 (fibronectin receptor beta), Itgam - integrin alpha m, Itga6 - integrin alpha 6, Ddrgk1 - ddrgk domain containing 1, Gba - glucosidase, beta, acid, Ybx1 - y box protein 1, Tgm2 - transglutaminase 2, c polypeptide, Fga - fibrinogen alpha chain, Clu - clusterin, Irs2 - insulin receptor substrate 2, Pdcd5 - programmed cell death 5, Stat1 - signal transducer and activator of transcription 1] |
| GO:0042592 | homeostatic process | 2.59E-4 | 1.64E-2 | 1,91 | 4935 | 501 | 160 | 31 | [Wfs1 - wolfram syndrome 1 homolog (human), Plg - plasminogen, Lamp2 - lysosomal-associated membrane protein 2, P4hb - prolyl 4-hydroxylase, beta polypeptide, Cfh - complement component factor h, Gaa - glucosidase, alpha, acid, Sod1 - superoxide dismutase 1, soluble, Kng1 - kininogen 1, Ide - insulin degrading enzyme, Krt1 - keratin 1, Itgb1 - integrin beta 1 (fibronectin receptor beta), Cp - ceruloplasmin, F2 - coagulation factor ii, Fabp5 - fatty acid binding protein 5, epidermal, Kcnj10 - potassium inwardly-rectifying channel, subfamily j, member 10, Hexb - hexosaminidase b, Sh3bgrl3 - sh3 domain binding glutamic acid-rich protein-like 3, Rbp1 - retinol binding protein 1, cellular, Gba - glucosidase, beta, acid, Hpx - hemopexin, Tgm2 - transglutaminase 2, c polypeptide, Serpinb1a - serine (or cysteine) peptidase inhibitor, clade b, member 1a, Ampd3 - adenosine monophosphate deaminase 3, Ncstn - nicastrin, Apoa1 - apolipoprotein a-i, Car14 - carbonic anhydrase 14, Txndc5 - thioredoxin domain containing 5, Commd1 - comm domain containing 1, Stat3 - signal transducer and activator of transcription 3, Stat1 - signal transducer and activator of transcription 1, Scn3b - sodium channel, voltage-gated, type iii, beta] |
| GO:0016485 | protein processing | 2.62E-4 | 1.65E-2 | 4,66 | 4935 | 53 | 160 | 8 | [Apoh - apolipoprotein h, Hp - haptoglobin, Lgmn - legumain, Fga - fibrinogen alpha chain, Fgg - fibrinogen gamma chain, Ncstn - nicastrin, Cpe - carboxypeptidase e, Fgb - fibrinogen beta chain] |
| GO:0042110 | T cell activation | 2.62E-4 | 1.64E-2 | 4,66 | 4935 | 53 | 160 | 8 | [Sema4a - sema domain, immunoglobulin domain (ig), transmembrane domain (tm) and short cytoplasmic domain, (semaphorin) 4a, Itgam - integrin alpha m, Fcer1g - fc receptor, ige, high affinity i, gamma polypeptide, Ncstn - nicastrin, Lcp1 - lymphocyte cytosolic protein 1, Cd151 - cd151 antigen, Gba - glucosidase, beta, acid, Itgb2 - integrin beta 2] |
| GO:0070661 | leukocyte proliferation | 2.7E-4 | 1.68E-2 | 6,38 | 4935 | 29 | 160 | 6 | [Itgam - integrin alpha m, Ncstn - nicastrin, Clu - clusterin, Cd151 - cd151 antigen, Itgb2 - integrin beta 2, Gba - glucosidase, beta, acid] |

| | | | | | | | | | |
|---|---|---|---|---|---|---|---|---|---|
| GO:0002819 | regulation of adaptive immune response | 2.7E-4 | 1.67E-2 | 6,38 | 4935 | 29 | 160 | 6 | [Hpx - hemopexin, C3 - complement component 3, H2-D1 - histocompatibility 2, d region locus 1, Ighg2b - immunoglobulin heavy constant gamma 2b, Ighg1 - immunoglobulin heavy constant gamma 1 (g1m marker), Fcer1g - fc receptor, ige, high affinity i, gamma polypeptide] |
| GO:1903524 | positive regulation of blood circulation | 2.71E-4 | 1.66E-2 | 8,12 | 4935 | 19 | 160 | 5 | [Fgg - fibrinogen gamma chain, Fga - fibrinogen alpha chain, Fgb - fibrinogen beta chain, Tpm1 - tropomyosin 1, alpha, Scn3b - sodium channel, voltage-gated, type iii, beta] |
| GO:0050764 | regulation of phagocytosis | 2.93E-4 | 1.79E-2 | 5,27 | 4935 | 41 | 160 | 7 | [C3 - complement component 3, Ahsg - alpha-2-hs-glycoprotein, Ighg2b - immunoglobulin heavy constant gamma 2b, Ighg1 - immunoglobulin heavy constant gamma 1 (g1m marker), Fcer1g - fc receptor, ige, high affinity i, gamma polypeptide, Apoa1 - apolipoprotein a-i, Sod1 - superoxide dismutase 1, soluble] |
| GO:0043302 | positive regulation of leukocyte degranulation | 2.94E-4 | 1.79E-2 | 11,22 | 4935 | 11 | 160 | 4 | [Itgam - integrin alpha m, Fcer1g - fc receptor, ige, high affinity i, gamma polypeptide, Lamp1 - lysosomal-associated membrane protein 1, Itgb2 - integrin beta 2] |
| GO:0061900 | glial cell activation | 2.94E-4 | 1.78E-2 | 11,22 | 4935 | 11 | 160 | 4 | [Itgam - integrin alpha m, Aif1 - allograft inflammatory factor 1, Clu - clusterin, C1qa - complement component 1, q subcomponent, alpha polypeptide] |
| GO:0051270 | regulation of cellular component movement | 2.99E-4 | 1.8E-2 | 2,08 | 4935 | 370 | 160 | 25 | [Anxa3 - annexin a3, Plg - plasminogen, Fn1 - fibronectin 1, Aif1 - allograft inflammatory factor 1, Lamp1 - lysosomal-associated membrane protein 1, Flna - filamin, alpha, Apoh - apolipoprotein h, Mcam - melanoma cell adhesion molecule, Apod - apolipoprotein d, Lgmn - legumain, Glipr2 - gli pathogenesis-related 2, Itgb1 - integrin beta 1 (fibronectin receptor beta), Vim - vimentin, Itga6 - integrin alpha 6, Hrg - histidine-rich glycoprotein, Vtn - vitronectin, Ddrgk1 - ddrgk domain containing 1, Itgb3 - integrin beta 3, Tpm1 - tropomyosin 1, alpha, Sema4a - sema domain, immunoglobulin domain (ig), transmembrane domain (tm) and short cytoplasmic domain, (semaphorin) 4a, Fga - fibrinogen alpha chain, Arhgdib - rho, gdp dissociation inhibitor (gdi) beta, Irs2 - insulin receptor substrate 2, Stat3 - signal transducer and activator of transcription 3, Cd151 - cd151 antigen] |
| GO:0051251 | positive regulation of lymphocyte activation | 2.99E-4 | 1.79E-2 | 4,57 | 4935 | 54 | 160 | 8 | [Igkc - immunoglobulin kappa constant, Lgals1 - lectin, galactose binding, soluble 1, Ighg2b - immunoglobulin heavy constant gamma 2b, Ighg1 - immunoglobulin heavy constant gamma 1 (g1m marker), Aif1 - allograft inflammatory factor 1, Ighm - immunoglobulin heavy constant mu, Irs2 - insulin receptor substrate 2, Lamp1 - lysosomal-associated membrane protein 1] |
| GO:0051918 | negative regulation of fibrinolysis | 3.19E-4 | 1.9E-2 | 18,51 | 4935 | 5 | 160 | 3 | [Apoh - apolipoprotein h, Plg - plasminogen, Hrg - histidine-rich glycoprotein] |
| GO:0051917 | regulation of fibrinolysis | 3.19E-4 | 1.89E-2 | 18,51 | 4935 | 5 | 160 | 3 | [Plg - plasminogen, Apoh - apolipoprotein h, Hrg - histidine-rich glycoprotein] |
| GO:0010001 | glial cell differentiation | 3.29E-4 | 1.94E-2 | 6,17 | 4935 | 30 | 160 | 6 | [Vim - vimentin, Gfap - glial fibrillary acidic protein, Stat3 - signal transducer and activator of transcription 3, Tspan2 - tetraspanin 2, Vtn - vitronectin, Gba - glucosidase, beta, acid] |
| GO:0051093 | negative regulation of developmental process | 3.45E-4 | 2.03E-2 | 2,25 | 4935 | 288 | 160 | 21 | [Itgb1 - integrin beta 1 (fibronectin receptor beta), Plg - plasminogen, H2-D1 - histocompatibility 2, d region locus 1, Vim - vimentin, Lgals1 - lectin, galactose binding, soluble 1, Ahsg - alpha-2-hs-glycoprotein, C1qc - complement component 1, q subcomponent, c chain, F2 - coagulation factor ii, Hrg - histidine-rich glycoprotein, Itgb3 - integrin beta 3, Ybx1 - y box protein 1, Flna - filamin, alpha, Sema4a - sema domain, immunoglobulin domain (ig), transmembrane domain (tm) and short cytoplasmic domain, (semaphorin) 4a, Apoh - apolipoprotein h, Lgmn - legumain, Gfap - glial fibrillary acidic protein, Arhgdib - rho, gdp dissociation inhibitor (gdi) beta, Stat3 - signal transducer and activator of transcription 3, Cpe - carboxypeptidase e, Stat1 - signal transducer and activator of transcription 1, Anxa2 - annexin a2] |
| GO:0090303 | positive regulation of wound healing | 3.51E-4 | 2.05E-2 | 7,71 | 4935 | 20 | 160 | 5 | [Apoh - apolipoprotein h, Itgb1 - integrin beta 1 (fibronectin receptor beta), Plg - plasminogen, F2 - coagulation factor ii, Hrg - histidine-rich glycoprotein] |
| GO:0046649 | lymphocyte activation | 3.73E-4 | 2.16E-2 | 3,63 | 4935 | 85 | 160 | 10 | [Igkc - immunoglobulin kappa constant, Sema4a - sema domain, immunoglobulin domain (ig), transmembrane domain (tm) and short cytoplasmic domain, (semaphorin) 4a, Lgals1 - lectin, galactose binding, soluble 1, Itgam - integrin alpha m, Fcer1g - fc receptor, ige, high affinity i, gamma polypeptide, Ncstn - nicastrin, Lcp1 - lymphocyte cytosolic protein 1, Cd151 - cd151 antigen, Gba - glucosidase, beta, acid, Itgb2 - integrin beta 2] |
| GO:0051258 | protein polymerization | 3.97E-4 | 2.29E-2 | 5,97 | 4935 | 31 | 160 | 6 | [Fga - fibrinogen alpha chain, Fgg - fibrinogen gamma chain, Aif1 - allograft inflammatory factor 1, Gsn - gelsolin, Vtn - vitronectin, Fgb - fibrinogen beta chain] |
| GO:1903036 | positive regulation of response to wounding | 3.97E-4 | 2.28E-2 | 5,97 | 4935 | 31 | 160 | 6 | [Flna - filamin, alpha, Itgb1 - integrin beta 1 (fibronectin receptor beta), Apoh - apolipoprotein h, Plg - plasminogen, F2 - coagulation factor ii, Hrg - histidine-rich glycoprotein] |

| | | | | | | | | | |
|---|---|---|---|---|---|---|---|---|---|
| GO:0010712 | regulation of collagen metabolic process | 4.3E-4 | 2.46E-2 | 10,28 | 4935 | 12 | 160 | 4 | [Itgb1 - integrin beta 1 (fibronectin receptor beta), Vim - vimentin, Fn1 - fibronectin 1, F2 - coagulation factor ii] |
| GO:0050820 | positive regulation of coagulation | 4.3E-4 | 2.45E-2 | 10,28 | 4935 | 12 | 160 | 4 | [Apoh - apolipoprotein h, Plg - plasminogen, F2 - coagulation factor ii, Hrg - histidine-rich glycoprotein] |
| GO:0042116 | macrophage activation | 4.3E-4 | 2.43E-2 | 10,28 | 4935 | 12 | 160 | 4 | [Itgam - integrin alpha m, Aif1 - allograft inflammatory factor 1, Clu - clusterin, C1qa - complement component 1, q subcomponent, alpha polypeptide] |
| GO:0010811 | positive regulation of cell-substrate adhesion | 4.38E-4 | 2.47E-2 | 4,33 | 4935 | 57 | 160 | 8 | [Flna - filamin, alpha, Itgb1 - integrin beta 1 (fibronectin receptor beta), Fn1 - fibronectin 1, P4hb - prolyl 4-hydroxylase, beta polypeptide, Itga6 - integrin alpha 6, Apoa1 - apolipoprotein a-i, Vtn - vitronectin, Itgb3 - integrin beta 3] |
| GO:0009892 | negative regulation of metabolic process | 4.6E-4 | 2.58E-2 | 1,64 | 4935 | 811 | 160 | 43 | [Serpina1b - serine (or cysteine) preptidase inhibitor, clade a, member 1b, Wfs1 - wolfram syndrome 1 homolog (human), Pzp - pregnancy zone protein, Serpina6 - serine (or cysteine) peptidase inhibitor, clade a, member 6, Fn1 - fibronectin 1, Serpina1a - serine (or cysteine) peptidase inhibitor, clade a, member 1a, Ahsg - alpha-2-hs-glycoprotein, Aif1 - allograft inflammatory factor 1, Serpinh1 - serine (or cysteine) peptidase inhibitor, clade h, member 1, Cst3 - cystatin c, Serpina1d - serine (or cysteine) peptidase inhibitor, clade a, member 1d, Serpina1e - serine (or cysteine) peptidase inhibitor, clade a, member 1e, Flna - filamin, alpha, Apoh - apolipoprotein h, Apod - apolipoprotein d, Lgmn - legumain, Serpinc1 - serine (or cysteine) peptidase inhibitor, clade c (antithrombin), member 1, Serpina3k - serine (or cysteine) peptidase inhibitor, clade a, member 3k, Ppp1r1b - protein phosphatase 1, regulatory (inhibitor) subunit 1b, Fhit - fragile histidine triad gene, Ctsa - cathepsin a, Sod1 - superoxide dismutase 1, soluble, Kng1 - kininogen 1, Ide - insulin degrading enzyme, Itgam - integrin alpha m, Hp - haptoglobin, F2 - coagulation factor ii, Hrg - histidine-rich glycoprotein, Ddrgk1 - ddrgk domain containing 1, Vtn - vitronectin, Itgb3 - integrin beta 3, Gba - glucosidase, beta, acid, Lsm6 - lsm6 homolog, u6 small nuclear rna associated (s. cerevisiae), Ybx1 - y box protein 1, Serpinb1a - serine (or cysteine) peptidase inhibitor, clade b, member 1a, Itih3 - inter-alpha trypsin inhibitor, heavy chain 3, Itih2 - inter-alpha trypsin inhibitor, heavy chain 2, Itih1 - inter-alpha trypsin inhibitor, heavy chain 1, Irs2 - insulin receptor substrate 2, Clu - clusterin, Mug1 - murinoglobulin 1, Stat3 - signal transducer and activator of transcription 3, Anxa2 - annexin a2] |
| GO:0050729 | positive regulation of inflammatory response | 4.76E-4 | 2.65E-2 | 5,78 | 4935 | 32 | 160 | 6 | [C3 - complement component 3, Tgm2 - transglutaminase 2, c polypeptide, Ighg2b - immunoglobulin heavy constant gamma 2b, Ighg1 - immunoglobulin heavy constant gamma 1 (g1m marker), Fcer1g - fc receptor, ige, high affinity i, gamma polypeptide, Stat3 - signal transducer and activator of transcription 3] |
| GO:0050864 | regulation of B cell activation | 5.66E-4 | 3.14E-2 | 7,01 | 4935 | 22 | 160 | 5 | [Igkc - immunoglobulin kappa constant, Ighg2b - immunoglobulin heavy constant gamma 2b, Ighg1 - immunoglobulin heavy constant gamma 1 (g1m marker), Ighm - immunoglobulin heavy constant mu, Irs2 - insulin receptor substrate 2] |
| GO:0050853 | B cell receptor signaling pathway | 6.05E-4 | 3.34E-2 | 9,49 | 4935 | 13 | 160 | 4 | [Igkc - immunoglobulin kappa constant, Ighg2b - immunoglobulin heavy constant gamma 2b, Ighg1 - immunoglobulin heavy constant gamma 1 (g1m marker), Ighm - immunoglobulin heavy constant mu] |
| GO:0097242 | amyloid-beta clearance | 6.05E-4 | 3.33E-2 | 9,49 | 4935 | 13 | 160 | 4 | [C3 - complement component 3, Itgam - integrin alpha m, Ide - insulin degrading enzyme, Itgb2 - integrin beta 2] |
| GO:0006957 | complement activation, alternative pathway | 6.23E-4 | 3.41E-2 | 15,42 | 4935 | 6 | 160 | 3 | [C3 - complement component 3, Cfb - complement factor b, Cfh - complement component factor h] |
| GO:0034116 | positive regulation of heterotypic cell-cell adhesion | 6.23E-4 | 3.39E-2 | 15,42 | 4935 | 6 | 160 | 3 | [Fga - fibrinogen alpha chain, Fgg - fibrinogen gamma chain, Fgb - fibrinogen beta chain] |
| GO:0030212 | hyaluronan metabolic process | 6.23E-4 | 3.37E-2 | 15,42 | 4935 | 6 | 160 | 3 | [Itih3 - inter-alpha trypsin inhibitor, heavy chain 3, Itih2 - inter-alpha trypsin inhibitor, heavy chain 2, Itih1 - inter-alpha trypsin inhibitor, heavy chain 1] |
| GO:0031349 | positive regulation of defense response | 6.99E-4 | 3.77E-2 | 3,65 | 4935 | 76 | 160 | 9 | [Hpx - hemopexin, C3 - complement component 3, Tgm2 - transglutaminase 2, c polypeptide, Irgm1 - immunity-related gtpase family m member 1, Ighg2b - immunoglobulin heavy constant gamma 2b, Ighg1 - immunoglobulin heavy constant gamma 1 (g1m marker), Fcer1g - fc receptor, ige, high affinity i, gamma polypeptide, Stat3 - signal transducer and activator of transcription 3, Lamp1 - lysosomal-associated membrane protein 1] |

| | | | | | | | | | |
|---|---|---|---|---|---|---|---|---|---|
| GO:0042221 | response to chemical | 7.84E-4 | 4.21E-2 | 1,58 | 4935 | 857 | 160 | 44 | [Serpina1b - serine (or cysteine) preptidase inhibitor, clade a, member 1b, Wfs1 - wolfram syndrome 1 homolog (human), Fn1 - fibronectin 1, Serpina1a - serine (or cysteine) peptidase inhibitor, clade a, member 1a, Aif1 - allograft inflammatory factor 1, S100a16 - s100 calcium binding protein a16, Gsn - gelsolin, Serpina1d - serine (or cysteine) peptidase inhibitor, clade a, member 1d, Serpina1e - serine (or cysteine) peptidase inhibitor, clade a, member 1e, Apod - apolipoprotein d, Lgmn - legumain, Irgm1 - immunity-related gtpase family m member 1, Serpinc1 - serine (or cysteine) peptidase inhibitor, clade c (antithrombin), member 1, P4hb - prolyl 4-hydroxylase, beta polypeptide, Serpina3k - serine (or cysteine) peptidase inhibitor, clade a, member 3k, Ppp1r1b - protein phosphatase 1, regulatory (inhibitor) subunit 1b, Fgg - fibrinogen gamma chain, Fcer1g - fc receptor, ige, high affinity i, gamma polypeptide, Cfh - complement component factor h, Sod1 - superoxide dismutase 1, soluble, Fgb - fibrinogen beta chain, Itgb1 - integrin beta 1 (fibronectin receptor beta), Vim - vimentin, Crot - carnitine o-octanoyltransferase, Cfb - complement factor b, Hp - haptoglobin, Cp - ceruloplasmin, Mgst1 - microsomal glutathione s-transferase 1, Hrg - histidine-rich glycoprotein, Itgb3 - integrin beta 3, Gba - glucosidase, beta, acid, Gss - glutathione synthetase, Itgb2 - integrin beta 2, Rbp1 - retinol binding protein 1, cellular, Ybx1 - y box protein 1, Fga - fibrinogen alpha chain, Pdcd5 - programmed cell death 5, Ncstn - nicastrin, Irs2 - insulin receptor substrate 2, Clu - clusterin, Stat3 - signal transducer and activator of transcription 3, Stat1 - signal transducer and activator of transcription 1, Tbl2 - transducin (beta)-like 2, Anxa2 - annexin a2] |
| GO:0034097 | response to cytokine | 8.19E-4 | 4.37E-2 | 2,51 | 4935 | 184 | 160 | 15 | [Serpina1b - serine (or cysteine) preptidase inhibitor, clade a, member 1b, Vim - vimentin, Serpina1a - serine (or cysteine) peptidase inhibitor, clade a, member 1a, Fn1 - fibronectin 1, Gsn - gelsolin, Serpina1d - serine (or cysteine) peptidase inhibitor, clade a, member 1d, Gba - glucosidase, beta, acid, Serpina1e - serine (or cysteine) peptidase inhibitor, clade a, member 1e, Ybx1 - y box protein 1, Irgm1 - immunity-related gtpase family m member 1, Serpina3k - serine (or cysteine) peptidase inhibitor, clade a, member 3k, P4hb - prolyl 4-hydroxylase, beta polypeptide, Cfh - complement component factor h, Stat3 - signal transducer and activator of transcription 3, Stat1 - signal transducer and activator of transcription 1] |
| GO:0051172 | negative regulation of nitrogen compound metabolic process | 8.71E-4 | 4.63E-2 | 1,70 | 4935 | 634 | 160 | 35 | [Serpina1b - serine (or cysteine) preptidase inhibitor, clade a, member 1b, Wfs1 - wolfram syndrome 1 homolog (human), Pzp - pregnancy zone protein, Serpina6 - serine (or cysteine) peptidase inhibitor, clade a, member 6, Serpina1a - serine (or cysteine) peptidase inhibitor, clade a, member 1a, Ahsg - alpha-2-hs-glycoprotein, Serpinh1 - serine (or cysteine) peptidase inhibitor, clade h, member 1, Cst3 - cystatin c, Serpina1d - serine (or cysteine) peptidase inhibitor, clade a, member 1d, Serpina1e - serine (or cysteine) peptidase inhibitor, clade a, member 1e, Flna - filamin, alpha, Apod - apolipoprotein d, Serpinc1 - serine (or cysteine) peptidase inhibitor, clade c (antithrombin), member 1, Ppp1r1b - protein phosphatase 1, regulatory (inhibitor) subunit 1b, Serpina3k - serine (or cysteine) peptidase inhibitor, clade a, member 3k, Fhit - fragile histidine triad gene, Ctsa - cathepsin a, Kng1 - kininogen 1, Ide - insulin degrading enzyme, Itgam - integrin alpha m, F2 - coagulation factor ii, Hrg - histidine-rich glycoprotein, Ddrgk1 - ddrgk domain containing 1, Vtn - vitronectin, Itgb3 - integrin beta 3, Gba - glucosidase, beta, acid, Ybx1 - y box protein 1, Itih3 - inter-alpha trypsin inhibitor, heavy chain 3, Serpinb1a - serine (or cysteine) peptidase inhibitor, clade b, member 1a, Itih2 - inter-alpha trypsin inhibitor, heavy chain 2, Itih1 - inter-alpha trypsin inhibitor, heavy chain 1, Clu - clusterin, Stat3 - signal transducer and activator of transcription 3, Mug1 - murinoglobulin 1, Anxa2 - annexin a2] |
| GO:0080090 | regulation of primary metabolic process | 8.8E-4 | 4.66E-2 | 1,40 | 4935 | 1458 | 160 | 66 | [Pzp - pregnancy zone protein, Wfs1 - wolfram syndrome 1 homolog (human), Serpina1b - serine (or cysteine) preptidase inhibitor, clade a, member 1b, Fn1 - fibronectin 1, Ahsg - alpha-2-hs-glycoprotein, Serpina1a - serine (or cysteine) peptidase inhibitor, clade a, member 1a, Aif1 - allograft inflammatory factor 1, Gsn - gelsolin, Serpina1d - serine (or cysteine) peptidase inhibitor, clade a, member 1d, Serpina1e - serine (or cysteine) peptidase inhibitor, clade a, member 1e, Flna - filamin, alpha, Apoh - apolipoprotein h, Apod - apolipoprotein d, Serpina3k - serine (or cysteine) peptidase inhibitor, clade a, member 3k, Ppp1r1b - protein phosphatase 1, regulatory (inhibitor) subunit 1b, Ctsa - cathepsin a, Kng1 - kininogen 1, Glipr2 - gli pathogenesis-related 2, Itgb1 - integrin beta 1 (fibronectin receptor beta), Vim - vimentin, Itgam - integrin alpha m, F2 - coagulation factor ii, Itga6 - integrin alpha 6, Fabp5 - fatty acid binding protein 5, epidermal, Psme1 - proteasome (prosome, macropain) activator subunit 1 (pa28 alpha), Vtn - vitronectin, Itgb3 - integrin beta 3, Gba - glucosidase, beta, acid, Itgb2 - integrin beta 2, Scarb2 - scavenger receptor class b, member 2, Serpinb1a - serine (or cysteine) peptidase inhibitor, clade b, member 1a, Itih3 - inter-alpha trypsin inhibitor, heavy chain 3, Itih2 - inter-alpha trypsin inhibitor, heavy chain 2, Ighm - immunoglobulin heavy constant mu, Itih1 - inter-alpha trypsin inhibitor, heavy chain 1, Irs2 - insulin receptor substrate 2, Stat3 - signal transducer and activator of transcription 3, Stat1 - signal transducer and activator of transcription 1, Anxa2 - annexin a2, Serpina6 - serine (or cysteine) peptidase inhibitor, clade a, member 6, Anxa3 - annexin a3, Serpinh1 - serine (or cysteine) peptidase inhibitor, clade h, member 1, Cst3 - cystatin c, Lgmn - legumain, Serpinc1 - serine (or cysteine) peptidase inhibitor, clade c (antithrombin), member 1, Fgg - fibrinogen gamma chain, Fhit - fragile histidine triad gene, Cfh - complement component factor h, Ctsd - cathepsin d, Sod1 - superoxide dismutase 1, soluble, Fgb - fibrinogen beta chain, Ide - insulin degrading enzyme, C3 - complement component 3, Hrg - histidine-rich glycoprotein, Ddrgk1 - ddrgk domain containing 1, Hexb - hexosaminidase b, Hpx - hemopexin, Ybx1 - y box protein 1, Gfap - glial fibrillary acidic protein, Fga - fibrinogen alpha chain, Pdcd5 - programmed cell death 5, Apoa1 - apolipoprotein a-i, Clu - clusterin, Ncstn - nicastrin, Commd1 - comm domain containing 1, Mug1 - murinoglobulin 1] |
| GO:0030100 | regulation of endocytosis | 9.31E-4 | 4.9E-2 | 2,59 | 4935 | 167 | 160 | 14 | [C3 - complement component 3, Itgb1 - integrin beta 1 (fibronectin receptor beta), Ahsg - alpha-2-hs-glycoprotein, Vtn - vitronectin, Itgb3 - integrin beta 3, Ighg2b - immunoglobulin heavy constant gamma 2b, Ighg1 - immunoglobulin heavy constant gamma 1 (g1m marker), Fcer1g - fc receptor, ige, high affinity i, gamma polypeptide, Ighm - immunoglobulin heavy constant mu, Apoa1 - apolipoprotein a-i, Clu - clusterin, Cd151 - cd151 antigen, Sod1 - superoxide dismutase 1, soluble, Anxa2 - annexin a2] |
| GO:0010950 | positive regulation of endopeptidase activity | 9.67E-4 | 5.07E-2 | 3,86 | 4935 | 64 | 160 | 8 | [Lgmn - legumain, Pdcd5 - programmed cell death 5, Ncstn - nicastrin, Gsn - gelsolin, Ctsd - cathepsin d, Psme1 - proteasome (prosome, macropain) activator subunit 1 (pa28 alpha), Stat3 - signal transducer and activator of transcription 3, Stat1 - signal transducer and activator of transcription 1] |

**enriched GO terms of decreased proteins (biological process)**

| GO Term | Description | P-value | FDR q-value | Enrichment | N | B | n | b | Genes |
|---|---|---|---|---|---|---|---|---|---|
| GO:0048169 | regulation of long-term neuronal synaptic plasticity | 4.39E-6 | 4.99E-2 | 19,35 | 4935 | 25 | 51 | 5 | [Dlg4 - discs, large homolog 4 (drosophila), Shank3 - sh3/ankyrin domain gene 3, Grin1 - glutamate receptor, ionotropic, nmda1 (zeta 1), Syngap1 - synaptic ras gtpase activating protein 1 homolog (rat), Grin2b - glutamate receptor, ionotropic, nmda2b (epsilon 2)] |
| GO:0099601 | regulation of neurotransmitter receptor activity | 7.73E-5 | 4.4E-1 | 11,00 | 4935 | 44 | 51 | 5 | [Dlg4 - discs, large homolog 4 (drosophila), Dlgap2 - discs, large (drosophila) homolog-associated protein 2, Shank3 - sh3/ankyrin domain gene 3, Gsg1l - gsg1-like, Begain - brain-enriched guanylate kinase-associated] |
| GO:0048168 | regulation of neuronal synaptic plasticity | 1.07E-4 | 4.05E-1 | 10,29 | 4935 | 47 | 51 | 5 | [Dlg4 - discs, large homolog 4 (drosophila), Shank3 - sh3/ankyrin domain gene 3, Grin1 - glutamate receptor, ionotropic, nmda1 (zeta 1), Syngap1 - synaptic ras gtpase activating protein 1 homolog (rat), Grin2b - glutamate receptor, ionotropic, nmda2b (epsilon 2)] |
| GO:0035405 | histone-threonine phosphorylation | 3.12E-4 | 8.88E-1 | 64,51 | 4935 | 3 | 51 | 2 | [Prkca - protein kinase c, alpha, Pkn1 - protein kinase n1] |
| GO:0051705 | multi-organism behavior | 3.32E-4 | 7.56E-1 | 11,73 | 4935 | 33 | 51 | 4 | [Dlg4 - discs, large homolog 4 (drosophila), Shank3 - sh3/ankyrin domain gene 3, Grin1 - glutamate receptor, ionotropic, nmda1 (zeta 1), Grin2b - glutamate receptor, ionotropic, nmda2b (epsilon 2)] |
| GO:0050905 | neuromuscular process | 3.71E-4 | 7.04E-1 | 7,93 | 4935 | 61 | 51 | 5 | [Dlg4 - discs, large homolog 4 (drosophila), Shank3 - sh3/ankyrin domain gene 3, Grin1 - glutamate receptor, ionotropic, nmda1 (zeta 1), Grin2b - glutamate receptor, ionotropic, nmda2b (epsilon 2), Rbfox1 - rna binding protein, fox-1 homolog (c. elegans) 1] |
| GO:0010469 | regulation of signaling receptor activity | 4,00E-04 | 6.51E-1 | 7,80 | 4935 | 62 | 51 | 5 | [Dlg4 - discs, large homolog 4 (drosophila), Dlgap2 - discs, large (drosophila) homolog-associated protein 2, Shank3 - sh3/ankyrin domain gene 3, Gsg1l - gsg1-like, Begain - brain-enriched guanylate kinase-associated] |
| GO:0043113 | receptor clustering | 4.18E-4 | 5.95E-1 | 11,06 | 4935 | 35 | 51 | 4 | [Dlg4 - discs, large homolog 4 (drosophila), Shank3 - sh3/ankyrin domain gene 3, Syngap1 - synaptic ras gtpase activating protein 1 homolog (rat), Grin2b - glutamate receptor, ionotropic, nmda2b (epsilon 2)] |
| GO:2000821 | regulation of grooming behavior | 6.2E-4 | 7.84E-1 | 48,38 | 4935 | 4 | 51 | 2 | [Dlg4 - discs, large homolog 4 (drosophila), Shank3 - sh3/ankyrin domain gene 3] |
| GO:0097553 | calcium ion transmembrane import into cytosol | 7.61E-4 | 8.66E-1 | 16,13 | 4935 | 18 | 51 | 3 | [Grin1 - glutamate receptor, ionotropic, nmda1 (zeta 1), Plcg2 - phospholipase c, gamma 2, Grin2b - glutamate receptor, ionotropic, nmda2b (epsilon 2)] |
| GO:0016570 | histone modification | 8.01E-4 | 8.29E-1 | 6,72 | 4935 | 72 | 51 | 5 | [Prkca - protein kinase c, alpha, Chd5 - chromodomain helicase dna binding protein 5, Pkn1 - protein kinase n1, Smarca4 - swi/snf related, matrix associated, actin dependent regulator of chromatin, subfamily a, member 4, Smyd3 - set and mynd domain containing 3] |
| GO:0016569 | covalent chromatin modification | 8.01E-4 | 7.6E-1 | 6,72 | 4935 | 72 | 51 | 5 | [Prkca - protein kinase c, alpha, Chd5 - chromodomain helicase dna binding protein 5, Pkn1 - protein kinase n1, Smarca4 - swi/snf related, matrix associated, actin dependent regulator of chromatin, subfamily a, member 4, Smyd3 - set and mynd domain containing 3] |
| GO:0045471 | response to ethanol | 8.97E-4 | 7.85E-1 | 15,28 | 4935 | 19 | 51 | 3 | [Prkca - protein kinase c, alpha, Grin1 - glutamate receptor, ionotropic, nmda1 (zeta 1), Grin2b - glutamate receptor, ionotropic, nmda2b (epsilon 2)] |
