## Supplementary Data S2c - LPS vs Intact for "Brain injury environment critically influences the connectivity of transplanted neurons"

| Protein FDR Confidence: Combined | Accession | # Unique Peptides | Gene symbol | Description | Abundance Ratio: (LPS) / (intact) | Abundance Ratio P-Value: (LPS) / (intact) |
|---|---|---|---|---|---|---|
| High | A2ALK8 | 3 | **Ptpn3** | Tyrosine-protein phosphatase non-receptor type 3 OS=Mus musculus OX=10090 GN=Ptpn3 PE=1 SV=1 | 16,61 | 0,0000 |
| Medium | Q09200 | 1 | **B4galnt1** | Beta-1,4 N-acetylgalactosaminyltransferase 1 OS=Mus musculus OX=10090 GN=B4galnt1 PE=1 SV=1 | 8,98 | 0,0000 |
| High | P01867 | 3 | **Igh-3** | Ig gamma-2B chain C region OS=Mus musculus OX=10090 GN=Igh-3 PE=1 SV=3 | 5,53 | 0,0000 |
| High | P04186 | 6 | **Cfb** | Complement factor B OS=Mus musculus OX=10090 GN=Cfb PE=1 SV=2 | 5,24 | 0,0000 |
| High | Q9R0H5 | 1 | **Krt71** | Keratin, type II cytoskeletal 71 OS=Mus musculus OX=10090 GN=Krt71 PE=1 SV=1 | 4,40 | 0,0000 |
| Medium | Q8CIH5 | 1 | **Plcg2** | 1-phosphatidylinositol 4,5-bisphosphate phosphodiesterase gamma-2 OS=Mus musculus OX=10090 GN=Plcg2 PE=1 SV=1 | 3,80 | 0,0000 |
| High | Q91X72 | 20 | **Hpx** | Hemopexin OS=Mus musculus OX=10090 GN=Hpx PE=1 SV=2 | 2,92 | 0,0003 |
| High | P01864 | 3 | | Ig gamma-2A chain C region secreted form OS=Mus musculus OX=10090 PE=1 SV=1 | 2,84 | 0,0012 |
| High | Q09143 | 2 | **Slc7a1** | High affinity cationic amino acid transporter 1 OS=Mus musculus OX=10090 GN=Slc7a1 PE=1 SV=1 | 2,65 | 0,0016 |
| High | A2AT37 | 7 | **Upf2** | Regulator of nonsense transcripts 2 OS=Mus musculus OX=10090 GN=Upf2 PE=1 SV=1 | 2,61 | 0,0035 |
| High | P59326 | 2 | **Ythdf1** | YTH domain-containing family protein 1 OS=Mus musculus OX=10090 GN=Ythdf1 PE=1 SV=1 | 2,60 | 0,0027 |
| High | P01837 | 3 | **Igkc** | Immunoglobulin kappa constant OS=Mus musculus OX=10090 GN=Igkc PE=1 SV=2 | 2,49 | 0,0082 |
| High | Q61235 | 6 | **Sntb2** | Beta-2-syntrophin OS=Mus musculus OX=10090 GN=Sntb2 PE=1 SV=2 | 2,46 | 0,0057 |
| High | Q6NXH9 | 1 | **Krt73** | Keratin, type II cytoskeletal 73 OS=Mus musculus OX=10090 GN=Krt73 PE=1 SV=1 | 2,45 | 0,0013 |
| High | P07309 | 2 | **Ttr** | Transthyretin OS=Mus musculus OX=10090 GN=Ttr PE=1 SV=1 | 2,43 | 0,0055 |
| High | O35143 | 2 | **ATP5IF1** | ATPase inhibitor, mitochondrial OS=Mus musculus OX=10090 GN=ATP5IF1 PE=1 SV=2 | 2,43 | 0,0046 |
| High | Q9JJU8 | 4 | **Sh3bgrl** | SH3 domain-binding glutamic acid-rich-like protein OS=Mus musculus OX=10090 GN=Sh3bgrl PE=1 SV=1 | 2,36 | 0,0050 |
| High | P03987 | 5 | | Ig gamma-3 chain C region OS=Mus musculus OX=10090 PE=1 SV=2 | 2,34 | 0,0061 |
| High | Q8R332 | 2 | **Nup58** | Nucleoporin p58/p45 OS=Mus musculus OX=10090 GN=Nup58 PE=1 SV=1 | 2,31 | 0,0048 |
| High | Q60590 | 3 | **Orm1** | Alpha-1-acid glycoprotein 1 OS=Mus musculus OX=10090 GN=Orm1 PE=1 SV=1 | 2,30 | 0,0115 |
| High | P62073 | 2 | **Timm10** | Mitochondrial import inner membrane translocase subunit Tim10 OS=Mus musculus OX=10090 GN=Timm10 PE=1 SV=1 | 2,24 | 0,0107 |
| High | Q91Z38 | 3 | **Ttc1** | Tetratricopeptide repeat protein 1 OS=Mus musculus OX=10090 GN=Ttc1 PE=1 SV=1 | 2,21 | 0,0086 |
| High | O88351 | 4 | **Ikbkb** | Inhibitor of nuclear factor kappa-B kinase subunit beta OS=Mus musculus OX=10090 GN=Ikbkb PE=1 SV=1 | 2,20 | 0,0089 |
| High | Q9WVA2 | 2 | **Timm8a1** | Mitochondrial import inner membrane translocase subunit Tim8 A OS=Mus musculus OX=10090 GN=Timm8a1 PE=1 SV=1 | 2,20 | 0,0093 |
| High | P02769 | 33 | **ALB** | Serum albumin OS=Bos taurus OX=9913 GN=ALB PE=1 SV=4 | 2,18 | 0,0260 |
| High | P54726 | 3 | **Rad23a** | UV excision repair protein RAD23 homolog A OS=Mus musculus OX=10090 GN=Rad23a PE=1 SV=2 | 2,17 | 0,0100 |
| High | Q64436 | 3 | **Atp4a** | Potassium-transporting ATPase alpha chain 1 OS=Mus musculus OX=10090 GN=Atp4a PE=1 SV=3 | 2,15 | 0,0208 |
| High | Q9R1Z7 | 4 | **Pts** | 6-pyruvoyl tetrahydrobiopterin synthase OS=Mus musculus OX=10090 GN=Pts PE=1 SV=2 | 2,09 | 0,0137 |
| High | Q61646 | 8 | **Hp** | Haptoglobin OS=Mus musculus OX=10090 GN=Hp PE=1 SV=1 | 2,08 | 0,0114 |
| High | Q9DB25 | 3 | **Alg5** | Dolichyl-phosphate beta-glucosyltransferase OS=Mus musculus OX=10090 GN=Alg5 PE=1 SV=1 | 2,05 | 0,0203 |
| High | Q8JZV7 | 2 | **Amdhd2** | N-acetylglucosamine-6-phosphate deacetylase OS=Mus musculus OX=10090 GN=Amdhd2 PE=1 SV=1 | 2,03 | 0,0268 |
| Medium | Q3UL36 | 1 | **Arglu1** | Arginine and glutamate-rich protein 1 OS=Mus musculus OX=10090 GN=Arglu1 PE=1 SV=2 | 2,02 | 0,0175 |
| High | Q9Z0S9 | 2 | **Rabac1** | Prenylated Rab acceptor protein 1 OS=Mus musculus OX=10090 GN=Rabac1 PE=1 SV=1 | 2,01 | 0,0245 |
| High | Q91VW3 | 2 | **Sh3bgrl3** | SH3 domain-binding glutamic acid-rich-like protein 3 OS=Mus musculus OX=10090 GN=Sh3bgrl3 PE=1 SV=1 | 2,00 | 0,0223 |
| High | Q8BHL8 | 2 | **Psmf1** | Proteasome inhibitor PI31 subunit OS=Mus musculus OX=10090 GN=Psmf1 PE=1 SV=1 | 1,99 | 0,0229 |
| High | P17225 | 4 | **Ptbp1** | Polypyrimidine tract-binding protein 1 OS=Mus musculus OX=10090 GN=Ptbp1 PE=1 SV=2 | 1,97 | 0,0218 |
| High | P29699 | 7 | **Ahsg** | Alpha-2-HS-glycoprotein OS=Mus musculus OX=10090 GN=Ahsg PE=1 SV=1 | 1,97 | 0,0331 |
| High | Q8K045 | 1 | **Pkn3** | Serine/threonine-protein kinase N3 OS=Mus musculus OX=10090 GN=Pkn3 PE=1 SV=1 | 1,95 | 0,0201 |
| High | P20918 | 18 | **Plg** | Plasminogen OS=Mus musculus OX=10090 GN=Plg PE=1 SV=3 | 1,94 | 0,0304 |
| High | Q9DCU2 | 2 | **Pllp** | Plasmolipin OS=Mus musculus OX=10090 GN=Pllp PE=1 SV=1 | 1,93 | 0,0383 |
| High | P52503 | 5 | **Ndufs6** | NADH dehydrogenase [ubiquinone] iron-sulfur protein 6, mitochondrial OS=Mus musculus OX=10090 GN=Ndufs6 PE=1 SV=2 | 1,92 | 0,0239 |
| High | P31786 | 2 | **Dbi** | Acyl-CoA-binding protein OS=Mus musculus OX=10090 GN=Dbi PE=1 SV=2 | 1,90 | 0,0182 |
| High | Q05816 | 9 | **Fabp5** | Fatty acid-binding protein 5 OS=Mus musculus OX=10090 GN=Fabp5 PE=1 SV=3 | 1,88 | 0,0445 |
| High | Q9ET22 | 4 | **Dpp7** | Dipeptidyl peptidase 2 OS=Mus musculus OX=10090 GN=Dpp7 PE=1 SV=2 | 1,87 | 0,0298 |
| High | Q03958 | 3 | **Pfdn6** | Prefoldin subunit 6 OS=Mus musculus OX=10090 GN=Pfdn6 PE=1 SV=1 | 1,87 | 0,0284 |
| High | P16045 | 5 | **Lgals1** | Galectin-1 OS=Mus musculus OX=10090 GN=Lgals1 PE=1 SV=3 | 1,86 | 0,0288 |
| High | Q8R3Q6 | 4 | **Ccdc58** | Coiled-coil domain-containing protein 58 OS=Mus musculus OX=10090 GN=Ccdc58 PE=1 SV=1 | 1,86 | 0,0476 |
| High | P08228 | 7 | **Sod1** | Superoxide dismutase [Cu-Zn] OS=Mus musculus OX=10090 GN=Sod1 PE=1 SV=2 | 1,86 | 0,0263 |
| High | Q8VCI5 | 5 | **Pex19** | Peroxisomal biogenesis factor 19 OS=Mus musculus OX=10090 GN=Pex19 PE=1 SV=1 | 1,82 | 0,0343 |
| Medium | Q8BL80 | 1 | **Arhgap22** | Rho GTPase-activating protein 22 OS=Mus musculus OX=10090 GN=Arhgap22 PE=1 SV=2 | 1,82 | 0,0480 |
| High | P97929 | 1 | **Brca2** | Breast cancer type 2 susceptibility protein homolog OS=Mus musculus OX=10090 GN=Brca2 PE=1 SV=2 | 1,82 | 0,0492 |
| High | Q62288 | 4 | **Spock1** | Testican-1 OS=Mus musculus OX=10090 GN=Spock1 PE=2 SV=2 | 1,80 | 0,0432 |
| High | P97450 | 4 | **Atp5pf** | ATP synthase-coupling factor 6, mitochondrial OS=Mus musculus OX=10090 GN=Atp5pf PE=1 SV=1 | 1,77 | 0,0389 |
| High | Q80V26 | 8 | **Impad1** | Golgi-resident adenosine 3',5'-bisphosphate 3'-phosphatase OS=Mus musculus OX=10090 GN=Impad1 PE=1 SV=1 | 1,77 | 0,0396 |
| High | P21550 | 1 | **Eno3** | Beta-enolase OS=Mus musculus OX=10090 GN=Eno3 PE=1 SV=3 | 1,77 | 0,0405 |
| High | O08677 | 11 | **Kng1** | Kininogen-1 OS=Mus musculus OX=10090 GN=Kng1 PE=1 SV=1 | 1,76 | 0,0475 |
| High | Q91WK5 | 2 | **Gcsh** | Glycine cleavage system H protein, mitochondrial OS=Mus musculus OX=10090 GN=Gcsh PE=1 SV=2 | 1,75 | 0,0465 |
| High | P62858 | 2 | **Rps28** | 40S ribosomal protein S28 OS=Mus musculus OX=10090 GN=Rps28 PE=1 SV=1 | 1,75 | 0,0428 |
| High | Q9CQW0 | 2 | **Emc6** | ER membrane protein complex subunit 6 OS=Mus musculus OX=10090 GN=Emc6 PE=1 SV=1 | 1,74 | 0,0494 |
| High | O70591 | 4 | **Pfdn2** | Prefoldin subunit 2 OS=Mus musculus OX=10090 GN=Pfdn2 PE=1 SV=2 | 1,72 | 0,0489 |
| High | Q923T9 | 15 | **Camk2g** | Calcium/calmodulin-dependent protein kinase type II subunit gamma OS=Mus musculus OX=10090 GN=Camk2g PE=1 SV=1 | 0,56 | 0,0470 |
| High | Q9QWI6 | 53 | **Srcin1** | SRC kinase signaling inhibitor 1 OS=Mus musculus OX=10090 GN=Srcin1 PE=1 SV=2 | 0,56 | 0,0423 |
| High | Q4ACU6 | 53 | **Shank3** | SH3 and multiple ankyrin repeat domains protein 3 OS=Mus musculus OX=10090 GN=Shank3 PE=1 SV=3 | 0,55 | 0,0408 |
| High | F6SEU4 | 51 | **Syngap1** | Ras/Rap GTPase-activating protein SynGAP OS=Mus musculus OX=10090 GN=Syngap1 PE=1 SV=2 | 0,55 | 0,0367 |
| High | Q60698 | 1 | **Ski** | Ski oncogene OS=Mus musculus OX=10090 GN=Ski PE=1 SV=2 | 0,54 | 0,0330 |
| High | Q68EF6 | 25 | **Begain** | Brain-enriched guanylate kinase-associated protein OS=Mus musculus OX=10090 GN=Begain PE=1 SV=2 | 0,53 | 0,0292 |
| High | Q9CXS4 | 11 | **Cenpv** | Centromere protein V OS=Mus musculus OX=10090 GN=Cenpv PE=1 SV=2 | 0,53 | 0,0468 |
| High | Q80TE7 | 45 | **Lrrc7** | Leucine-rich repeat-containing protein 7 OS=Mus musculus OX=10090 GN=Lrrc7 PE=1 SV=2 | 0,51 | 0,0415 |
| High | P62911 | 6 | **Rpl32** | 60S ribosomal protein L32 OS=Mus musculus OX=10090 GN=Rpl32 PE=1 SV=2 | 0,50 | 0,0248 |
| High | P62892 | 1 | **Rpl39** | 60S ribosomal protein L39 OS=Mus musculus OX=10090 GN=Rpl39 PE=1 SV=2 | 0,50 | 0,0443 |
| High | Q3UXZ6 | 17 | **Fam81a** | Protein FAM81A OS=Mus musculus OX=10090 GN=Fam81a PE=1 SV=2 | 0,50 | 0,0359 |
| High | P83093 | 9 | **Stim2** | Stromal interaction molecule 2 OS=Mus musculus OX=10090 GN=Stim2 PE=1 SV=2 | 0,49 | 0,0363 |

| | | | | | | |
|---|---|---|---|---|---|---|
| High | Q80W54 | 7 | **Zmpste24** | CAAX prenyl protease 1 homolog OS=Mus musculus OX=10090 GN=Zmpste24 PE=1 SV=2 | 0,49 | 0,0448 |
| High | P41105 | 6 | **Rpl28** | 60S ribosomal protein L28 OS=Mus musculus OX=10090 GN=Rpl28 PE=1 SV=2 | 0,48 | 0,0158 |
| Medium | Q9QX98 | 1 | **Ptf1a** | Pancreas transcription factor 1 subunit alpha OS=Mus musculus OX=10090 GN=Ptf1a PE=1 SV=1 | 0,46 | 0,0267 |
| High | Q9D338 | 6 | **Mrpl19** | 39S ribosomal protein L19, mitochondrial OS=Mus musculus OX=10090 GN=Mrpl19 PE=1 SV=1 | 0,46 | 0,0201 |
| High | Q8C015 | 21 | **Pak5** | Serine/threonine-protein kinase PAK 5 OS=Mus musculus OX=10090 GN=Pak5 PE=1 SV=1 | 0,46 | 0,0200 |
| High | B2RWJ3 | 2 | **Tmem240** | Transmembrane protein 240 OS=Mus musculus OX=10090 GN=Tmem240 PE=1 SV=1 | 0,45 | 0,0427 |
| High | Q8K1A5 | 1 | **Tmem41b** | Transmembrane protein 41B OS=Mus musculus OX=10090 GN=Tmem41b PE=1 SV=1 | 0,45 | 0,0294 |
| Medium | Q8BH32 | 1 | **Susd4** | Sushi domain-containing protein 4 OS=Mus musculus OX=10090 GN=Susd4 PE=2 SV=1 | 0,45 | 0,0300 |
| High | Q8R2U6 | 3 | **Nudt4** | Diphosphoinositol polyphosphate phosphohydrolase 2 OS=Mus musculus OX=10090 GN=Nudt4 PE=1 SV=1 | 0,43 | 0,0410 |
| High | P02463 | 5 | **Col4a1** | Collagen alpha-1(IV) chain OS=Mus musculus OX=10090 GN=Col4a1 PE=1 SV=4 | 0,41 | 0,0060 |
| High | Q922U1 | 4 | **Prpf3** | U4/U6 small nuclear ribonucleoprotein Prp3 OS=Mus musculus OX=10090 GN=Prpf3 PE=1 SV=1 | 0,41 | 0,0334 |
| High | Q80XU3 | 2 | **Nucks1** | Nuclear ubiquitous casein and cyclin-dependent kinase substrate 1 OS=Mus musculus OX=10090 GN=Nucks1 PE=1 SV=1 | 0,41 | 0,0108 |
| Medium | Q4FZF3 | 1 | **Ddx49** | Probable ATP-dependent RNA helicase DDX49 OS=Mus musculus OX=10090 GN=Ddx49 PE=2 SV=1 | 0,41 | 0,0073 |
| High | Q91YW3 | 4 | **Dnajc3** | DnaJ homolog subfamily C member 3 OS=Mus musculus OX=10090 GN=Dnajc3 PE=1 SV=1 | 0,39 | 0,0295 |
| High | Q5XJY4 | 2 | **Parl** | Presenilins-associated rhomboid-like protein, mitochondrial OS=Mus musculus OX=10090 GN=Parl PE=1 SV=1 | 0,38 | 0,0345 |
| High | O35316 | 2 | **Slc6a6** | Sodium- and chloride-dependent taurine transporter OS=Mus musculus OX=10090 GN=Slc6a6 PE=1 SV=2 | 0,37 | 0,0137 |
| High | Q9EQJ9 | 9 | **Magi3** | Membrane-associated guanylate kinase, WW and PDZ domain-containing protein 3 OS=Mus musculus OX=10090 GN=Magi3 PE=1 SV=2 | 0,36 | 0,0208 |
| High | Q99KN2 | 2 | **Ciao1** | SV=1 | 0,35 | 0,0264 |
| High | Q99LC8 | 4 | **Eif2b1** | Translation initiation factor eIF-2B subunit alpha OS=Mus musculus OX=10090 GN=Eif2b1 PE=1 SV=1 | 0,35 | 0,0097 |
| High | Q9CQQ0 | 1 | **Smim8** | Small integral membrane protein 8 OS=Mus musculus OX=10090 GN=Smim8 PE=1 SV=1 | 0,33 | 0,0059 |
| High | Q8BVP5 | 1 | **Csnk1g2** | Casein kinase I isoform gamma-2 OS=Mus musculus OX=10090 GN=Csnk1g2 PE=1 SV=1 | 0,33 | 0,0025 |
| High | Q91YR7 | 13 | **Prpf6** | Pre-mRNA-processing factor 6 OS=Mus musculus OX=10090 GN=Prpf6 PE=1 SV=1 | 0,32 | 0,0009 |
| High | Q6PFX7 | 3 | **Nyap1** | Neuronal tyrosine-phosphorylated phosphoinositide-3-kinase adapter 1 OS=Mus musculus OX=10090 GN=Nyap1 PE=1 SV=1 | 0,32 | 0,0222 |
| Medium | Q9QXM1 | 2 | **Jmy** | Junction-mediating and -regulatory protein OS=Mus musculus OX=10090 GN=Jmy PE=1 SV=1 | 0,32 | 0,0217 |
| High | Q91XY4 | 2 | **Pcdhga4** | Protocadherin gamma-A4 OS=Mus musculus OX=10090 GN=Pcdhga4 PE=1 SV=1 | 0,30 | 0,0001 |
| High | Q8K0F1 | 4 | **Tbc1d23** | TBC1 domain family member 23 OS=Mus musculus OX=10090 GN=Tbc1d23 PE=1 SV=1 | 0,29 | 0,0019 |
| High | Q9QYI4 | 2 | **Dnajb12** | DnaJ homolog subfamily B member 12 OS=Mus musculus OX=10090 GN=Dnajb12 PE=1 SV=2 | 0,28 | 0,0001 |
| High | Q8C4G9 | 2 | **Adgra1** | Adhesion G protein-coupled receptor A1 OS=Mus musculus OX=10090 GN=Adgra1 PE=2 SV=1 | 0,28 | 0,0011 |
| Medium | Q8BGE5 | 1 | **Fancm** | Fanconi anemia group M protein homolog OS=Mus musculus OX=10090 GN=Fancm PE=1 SV=3 | 0,27 | 0,0000 |
| High | P97770 | 2 | **Thumpd3** | THUMP domain-containing protein 3 OS=Mus musculus OX=10090 GN=Thumpd3 PE=1 SV=1 | 0,25 | 0,0000 |
| High | Q6GQW0 | 4 | **Btbd11** | SV=2 | 0,24 | 0,0004 |
| Medium | Q9D384 | 1 | **Snrnp35** | U11/U12 small nuclear ribonucleoprotein 35 kDa protein OS=Mus musculus OX=10090 GN=Snrnp35 PE=2 SV=1 | 0,24 | 0,0000 |
| High | Q9Z2E9 | 2 | **Bscl2** | Seipin OS=Mus musculus OX=10090 GN=Bscl2 PE=1 SV=2 | 0,23 | 0,0002 |
| High | Q9R0M0 | 2 | **Celsr2** | Cadherin EGF LAG seven-pass G-type receptor 2 OS=Mus musculus OX=10090 GN=Celsr2 PE=1 SV=3 | 0,23 | 0,0004 |
| High | P60191 | 5 | Rims4 | Regulating synaptic membrane exocytosis protein 4 OS=Mus musculus OX=10090 GN=Rims4 PE=1 SV=1 | 0,22 | 0,0000 |
| High | P63054 | 3 | **Pcp4** | Calmodulin regulator protein PCP4 OS=Mus musculus OX=10090 GN=Pcp4 PE=3 SV=2 | 0,18 | 0,0000 |
| High | P58064 | 2 | **Mrps6** | 28S ribosomal protein S6, mitochondrial OS=Mus musculus OX=10090 GN=Mrps6 PE=1 SV=3 | 0,17 | 0,0000 |
| High | Q8C0Q9 | 1 | **Rapgef5** | Rap guanine nucleotide exchange factor 5 OS=Mus musculus OX=10090 GN=Rapgef5 PE=2 SV=2 | 0,16 | 0,0000 |
| High | Q6ZWQ7 | 3 | **Spcs3** | Signal peptidase complex subunit 3 OS=Mus musculus OX=10090 GN=Spcs3 PE=1 SV=1 | 0,16 | 0,0000 |
| High | O35963 | 4 | **Rab33b** | Ras-related protein Rab-33B OS=Mus musculus OX=10090 GN=Rab33b PE=1 SV=1 | 0,16 | 0,0000 |
| High | Q99N92 | 2 | **Mrpl27** | 39S ribosomal protein L27, mitochondrial OS=Mus musculus OX=10090 GN=Mrpl27 PE=1 SV=1 | 0,15 | 0,0000 |
| High | P62984 | 1 | **Uba52** | Ubiquitin-60S ribosomal protein L40 OS=Mus musculus OX=10090 GN=Uba52 PE=1 SV=2 | 0,15 | 0,0000 |
| High | Q61738 | 2 | **Itga7** | Integrin alpha-7 OS=Mus musculus OX=10090 GN=Itga7 PE=1 SV=3 | 0,11 | 0,0000 |
| High | P56387 | 3 | **Dynlt3** | Dynein light chain Tctex-type 3 OS=Mus musculus OX=10090 GN=Dynlt3 PE=1 SV=1 | 0,05 | 0,0000 |
| Medium | Q9D706 | 1 | **Rpap3** | RNA polymerase II-associated protein 3 OS=Mus musculus OX=10090 GN=Rpap3 PE=1 SV=1 | 0,02 | 0,0000 |
| High | Q80TH2 | 1 | **Erbin** | Erbin OS=Mus musculus OX=10090 GN=Erbin PE=1 SV=3 | 0,01 | 0,0000 |
| High | Q8BUM6 | 1 | **Fam163b** | Protein FAM163B OS=Mus musculus OX=10090 GN=Fam163b PE=1 SV=1 | 0,01 | 0,0000 |
| High | Q61762 | 1 | **Kcna5** | Potassium voltage-gated channel subfamily A member 5 OS=Mus musculus OX=10090 GN=Kcna5 PE=2 SV=2 | 0,01 | 0,0000 |
| High | O70456 | 2 | **Sfn** | 14-3-3 protein sigma OS=Mus musculus OX=10090 GN=Sfn PE=1 SV=2 | 0,01 | 0,0000 |
| Medium | D3Z7Q2 | 1 | **Smim20** | Small integral membrane protein 20 OS=Mus musculus OX=10090 GN=Smim20 PE=1 SV=1 | 0,01 | 0,0000 |
| High | B7ZMP1 | 3 | **Xpnpep3** | Xaa-Pro aminopeptidase 3 OS=Mus musculus OX=10090 GN=Xpnpep3 PE=1 SV=1 | 0,01 | 0,0000 |
