## Supplementary Data S2d - GO terms LPS for "Brain injury environment critically influences the connectivity of transplanted neurons"

### enriched GO terms of increased proteins (biological process)

| GO Term | Description | P-value | FDR q-value | Enrichment | N | B | n | b | Genes |
|---|---|---|---|---|---|---|---|---|---|
| GO:0072321 | chaperone-mediated protein transport | 7.05E-5 | 8.03E-1 | 33,65 | 4935 | 8 | 55 | 3 | [Timm8a1 - translocase of inner mitochondrial membrane 8a1, Timm10 - translocase of inner mitochondrial membrane 10, Pex19 - peroxisomal biogenesis factor 19] |
| GO:0006953 | acute-phase response | 1.49E-4 | 8.46E-1 | 26,92 | 4935 | 10 | 55 | 3 | [Ahsg - alpha-2-hs-glycoprotein, Hp - haptoglobin, Orm1 - orosomucoid 1] |
| GO:0050853 | B cell receptor signaling pathway | 3.46E-4 | 1,00E+00 | 20,71 | 4935 | 13 | 55 | 3 | [Igkc - immunoglobulin kappa constant, Ighg2b - immunoglobulin heavy constant gamma 2b, Plcg2 - phospholipase c, gamma 2] |
| GO:0002526 | acute inflammatory response | 4.37E-4 | 1,00E+00 | 19,23 | 4935 | 14 | 55 | 3 | [Ahsg - alpha-2-hs-glycoprotein, Hp - haptoglobin, Orm1 - orosomucoid 1] |
| GO:0030183 | B cell differentiation | 5.42E-4 | 1,00E+00 | 17,95 | 4935 | 15 | 55 | 3 | [Igkc - immunoglobulin kappa constant, Lgals1 - lectin, galactose binding, soluble 1, Plcg2 - phospholipase c, gamma 2] |
| GO:0002313 | mature B cell differentiation involved in immune response | 7.21E-4 | 1,00E+00 | 44,86 | 4935 | 4 | 55 | 2 | [Lgals1 - lectin, galactose binding, soluble 1, Plcg2 - phospholipase c, gamma 2] |
| GO:0006956 | complement activation | 9.5E-4 | 1,00E+00 | 14,95 | 4935 | 18 | 55 | 3 | [Igkc - immunoglobulin kappa constant, Cfb - complement factor b, Ighg2b - immunoglobulin heavy constant gamma 2b] |

### enriched GO terms of decreased proteins (biological process)

| GO Term | Description | P-value | FDR q-value | Enrichment | N | B | n | b | Genes |
|---|---|---|---|---|---|---|---|---|---|
| GO:0036297 | interstrand cross-link repair | 1.55E-4 | 1,00E+00 | 79,60 | 4935 | 2 | 62 | 2 | [Nucks1 - nuclear casein kinase and cyclin-dependent kinase substrate 1, Fancm - fanconi anemia, complementation group m] |
