## Supplementary Data S2e - SW-specific proteins for "Brain injury environment critically influences the connectivity of transplanted neurons"

| Gene symbol | Description | Abundance Ratio: (ipsi) / (intact) | Abundance Ratio P-Value: (ipsi) / (intact) |
|---|---|---|---|
| **C4b** | Complement C4-B OS=Mus musculus OX=10090 GN=C4b PE=1 SV=3 | 100,00 | 0,0000 |
| **Serpinc1** | Antithrombin-III OS=Mus musculus OX=10090 GN=Serpinc1 PE=1 SV=1 | 12,07 | 0,0000 |
| **Gfap** | Glial fibrillary acidic protein OS=Mus musculus OX=10090 GN=Gfap PE=1 SV=4 | 7,65 | 0,0000 |
| **Serpina1e** | Alpha-1-antitrypsin 1-5 OS=Mus musculus OX=10090 GN=Serpina1e PE=1 SV=1 | 6,96 | 0,0000 |
| **Itih2** | Inter-alpha-trypsin inhibitor heavy chain H2 OS=Mus musculus OX=10090 GN=Itih2 PE=1 SV=1 | 6,92 | 0,0000 |
| **Alb** | Serum albumin OS=Mus musculus OX=10090 GN=Alb PE=1 SV=3 | 5,80 | 0,0000 |
| **Serpina1a** | Alpha-1-antitrypsin 1-1 OS=Mus musculus OX=10090 GN=Serpina1a PE=1 SV=4 | 5,79 | 0,0000 |
| **Ces1c** | Carboxylesterase 1C OS=Mus musculus OX=10090 GN=Ces1c PE=1 SV=4 | 5,28 | 0,0000 |
| **Tgm1** | Protein-glutamine gamma-glutamyltransferase K OS=Mus musculus OX=10090 GN=Tgm1 PE=1 SV=2 | 5,15 | 0,0000 |
| **Serpina6** | Corticosteroid-binding globulin OS=Mus musculus OX=10090 GN=Serpina6 PE=1 SV=1 | 5,09 | 0,0000 |
| **Gc** | Vitamin D-binding protein OS=Mus musculus OX=10090 GN=Gc PE=1 SV=2 | 4,74 | 0,0000 |
| **Serpina1b** | Alpha-1-antitrypsin 1-2 OS=Mus musculus OX=10090 GN=Serpina1b PE=1 SV=2 | 4,59 | 0,0000 |
| **Apoh** | Beta-2-glycoprotein 1 OS=Mus musculus OX=10090 GN=Apoh PE=1 SV=1 | 3,99 | 0,0000 |
| **Vtn** | Vitronectin OS=Mus musculus OX=10090 GN=Vtn PE=1 SV=2 | 3,88 | 0,0000 |
| **C3** | Complement C3 OS=Mus musculus OX=10090 GN=C3 PE=1 SV=3 | 3,58 | 0,0000 |
| **Serpina3k** | Serine protease inhibitor A3K OS=Mus musculus OX=10090 GN=Serpina3k PE=1 SV=2 | 3,44 | 0,0000 |
| **Tf** | Serotransferrin OS=Mus musculus OX=10090 GN=Tf PE=1 SV=1 | 3,41 | 0,0000 |
| **Vim** | Vimentin OS=Mus musculus OX=10090 GN=Vim PE=1 SV=3 | 3,21 | 0,0000 |
| **Itih3** | Inter-alpha-trypsin inhibitor heavy chain H3 OS=Mus musculus OX=10090 GN=Itih3 PE=1 SV=3 | 3,11 | 0,0000 |
| **H2-D1** | H-2 class I histocompatibility antigen, D-B alpha chain OS=Mus musculus OX=10090 GN=H2-D1 PE=1 SV=2 | 3,09 | 0,0000 |
| **Mug1** | Murinoglobulin-1 OS=Mus musculus OX=10090 GN=Mug1 PE=1 SV=3 | 3,02 | 0,0000 |
| **Itih1** | Inter-alpha-trypsin inhibitor heavy chain H1 OS=Mus musculus OX=10090 GN=Itih1 PE=1 SV=2 | 3,00 | 0,0000 |
| **Fgg** | Fibrinogen gamma chain OS=Mus musculus OX=10090 GN=Fgg PE=1 SV=1 | 2,94 | 0,0000 |
| **F2** | Prothrombin OS=Mus musculus OX=10090 GN=F2 PE=1 SV=1 | 2,84 | 0,0000 |
| **Fabp7** | Fatty acid-binding protein, brain OS=Mus musculus OX=10090 GN=Fabp7 PE=1 SV=2 | 2,67 | 0,0000 |
| **Fga** | Fibrinogen alpha chain OS=Mus musculus OX=10090 GN=Fga PE=1 SV=1 | 2,53 | 0,0000 |
| **Pzp** | Pregnancy zone protein OS=Mus musculus OX=10090 GN=Pzp PE=1 SV=3 | 2,51 | 0,0000 |
| **Fgb** | Fibrinogen beta chain OS=Mus musculus OX=10090 GN=Fgb PE=1 SV=1 | 2,48 | 0,0000 |
| **Aif1** | Allograft inflammatory factor 1 OS=Mus musculus OX=10090 GN=Aif1 PE=1 SV=1 | 2,33 | 0,0000 |
| **Cp** | Ceruloplasmin OS=Mus musculus OX=10090 GN=Cp PE=1 SV=2 | 2,29 | 0,0000 |
| **Apoa1** | Apolipoprotein A-I OS=Mus musculus OX=10090 GN=Apoa1 PE=1 SV=2 | 2,29 | 0,0000 |
| **Glipr2** | Golgi-associated plant pathogenesis-related protein 1 OS=Mus musculus OX=10090 GN=Glipr2 PE=1 SV=3 | 2,29 | 0,0000 |
| **Itgam** | Integrin alpha-M OS=Mus musculus OX=10090 GN=Itgam PE=1 SV=2 | 2,21 | 0,0000 |
| **Mgst1** | Microsomal glutathione S-transferase 1 OS=Mus musculus OX=10090 GN=Mgst1 PE=1 SV=3 | 2,17 | 0,0000 |
| **Ighg1** | Ig gamma-1 chain C region, membrane-bound form OS=Mus musculus OX=10090 GN=Ighg1 PE=1 SV=2 | 2,10 | 0,0000 |
| **Anxa2** | Annexin A2 OS=Mus musculus OX=10090 GN=Anxa2 PE=1 SV=2 | 2,10 | 0,0000 |
| **Serpinh1** | Serpin H1 OS=Mus musculus OX=10090 GN=Serpinh1 PE=1 SV=3 | 2,07 | 0,0000 |
| **Hrg** | Histidine-rich glycoprotein OS=Mus musculus OX=10090 GN=Hrg PE=1 SV=2 | 2,06 | 0,0011 |
| **C1qb** | Complement C1q subcomponent subunit B OS=Mus musculus OX=10090 GN=C1qb PE=1 SV=2 | 2,02 | 0,0000 |
| **Tagln2** | Transgelin-2 OS=Mus musculus OX=10090 GN=Tagln2 PE=1 SV=4 | 2,01 | 0,0000 |
| **Kcnj10** | ATP-sensitive inward rectifier potassium channel 10 OS=Mus musculus OX=10090 GN=Kcnj10 PE=1 SV=1 | 1,99 | 0,0000 |
| **Slc14a1** | Urea transporter 1 OS=Mus musculus OX=10090 GN=Slc14a1 PE=1 SV=2 | 1,97 | 0,0001 |
| **Itgb2** | Integrin beta-2 OS=Mus musculus OX=10090 GN=Itgb2 PE=1 SV=2 | 1,91 | 0,0001 |
| **C1qa** | Complement C1q subcomponent subunit A OS=Mus musculus OX=10090 GN=C1qa PE=1 SV=2 | 1,88 | 0,0000 |
| **Wfs1** | Wolframin OS=Mus musculus OX=10090 GN=Wfs1 PE=1 SV=1 | 1,86 | 0,0000 |
| **Cd151** | CD151 antigen OS=Mus musculus OX=10090 GN=Cd151 PE=1 SV=2 | 1,85 | 0,0007 |
| **Stat1** | Signal transducer and activator of transcription 1 OS=Mus musculus OX=10090 GN=Stat1 PE=1 SV=1 | 1,85 | 0,0005 |
| **Lcp1** | Plastin-2 OS=Mus musculus OX=10090 GN=Lcp1 PE=1 SV=4 | 1,81 | 0,0003 |
| **Serpina1d** | Alpha-1-antitrypsin 1-4 OS=Mus musculus OX=10090 GN=Serpina1d PE=1 SV=1 | 1,81 | 0,0001 |
| **Ide** | Insulin-degrading enzyme OS=Mus musculus OX=10090 GN=Ide PE=1 SV=1 | 1,78 | 0,0004 |
| **Mcam** | Cell surface glycoprotein MUC18 OS=Mus musculus OX=10090 GN=Mcam PE=1 SV=1 | 1,77 | 0,0044 |
| **Nap1l5** | Nucleosome assembly protein 1-like 5 OS=Mus musculus OX=10090 GN=Nap1l5 PE=1 SV=1 | 1,76 | 0,0001 |
| **Arhgdib** | Rho GDP-dissociation inhibitor 2 OS=Mus musculus OX=10090 GN=Arhgdib PE=1 SV=3 | 1,76 | 0,0020 |
| **Clic1** | Chloride intracellular channel protein 1 OS=Mus musculus OX=10090 GN=Clic1 PE=1 SV=3 | 1,76 | 0,0003 |
| **C1qc** | Complement C1q subcomponent subunit C OS=Mus musculus OX=10090 GN=C1qc PE=1 SV=2 | 1,72 | 0,0002 |
| **Cfh** | Complement factor H OS=Mus musculus OX=10090 GN=Cfh PE=1 SV=2 | 1,66 | 0,0061 |
| **Itga6** | Integrin alpha-6 OS=Mus musculus OX=10090 GN=Itga6 PE=1 SV=3 | 1,65 | 0,0022 |
| **Ybx1** | Y-box-binding protein 1 OS=Mus musculus OX=10090 GN=Ybx1 PE=1 SV=3 | 1,64 | 0,0006 |
| **Fn1** | Fibronectin OS=Mus musculus OX=10090 GN=Fn1 PE=1 SV=4 | 1,61 | 0,0053 |
| **Npl** | N-acetylneuraminate lyase OS=Mus musculus OX=10090 GN=Npl PE=1 SV=1 | 1,60 | 0,0092 |
| **Apod** | Apolipoprotein D OS=Mus musculus OX=10090 GN=Apod PE=1 SV=1 | 1,60 | 0,0091 |
| **Lactb2** | Endoribonuclease LACTB2 OS=Mus musculus OX=10090 GN=Lactb2 PE=1 SV=1 | 1,59 | 0,0093 |
| **Itgb3** | Integrin beta-3 OS=Mus musculus OX=10090 GN=Itgb3 PE=1 SV=2 | 1,59 | 0,0020 |
| **Stat3** | Signal transducer and activator of transcription 3 OS=Mus musculus OX=10090 GN=Stat3 PE=1 SV=2 | 1,57 | 0,0235 |
| **Clu** | Clusterin OS=Mus musculus OX=10090 GN=Clu PE=1 SV=1 | 1,56 | 0,0031 |
| **Nptx2** | Neuronal pentraxin-2 OS=Mus musculus OX=10090 GN=Nptx2 PE=2 SV=1 | 1,56 | 0,0021 |
| **Flna** | Filamin-A OS=Mus musculus OX=10090 GN=Flna PE=1 SV=5 | 1,56 | 0,0073 |
| **Arpc1b** | Actin-related protein 2/3 complex subunit 1B OS=Mus musculus OX=10090 GN=Arpc1b PE=1 SV=4 | 1,55 | 0,0078 |
| **Krt77** | Keratin, type II cytoskeletal 1b OS=Mus musculus OX=10090 GN=Krt77 PE=1 SV=1 | 1,54 | 0,0020 |
| **Fcer1g** | High affinity immunoglobulin epsilon receptor subunit gamma OS=Mus musculus OX=10090 GN=Fcer1g PE=1 SV=1 | 1,54 | 0,0106 |
| **Crip2** | Cysteine-rich protein 2 OS=Mus musculus OX=10090 GN=Crip2 PE=1 SV=1 | 1,54 | 0,0048 |
| **Ttc38** | Tetratricopeptide repeat protein 38 OS=Mus musculus OX=10090 GN=Ttc38 PE=1 SV=2 | 1,53 | 0,0165 |

| | | | |
|---|---|---|---|
| **Ddrgk1** | DDRGK domain-containing protein 1 OS=Mus musculus OX=10090 GN=Ddrgk1 PE=1 SV=2 | 1,53 | 0,0182 |
| **Psmg2** | Proteasome assembly chaperone 2 OS=Mus musculus OX=10090 GN=Psmg2 PE=1 SV=1 | 1,52 | 0,0031 |
| **Fhit** | Bis(5'-adenosyl)-triphosphatase OS=Mus musculus OX=10090 GN=Fhit PE=1 SV=3 | 1,51 | 0,0213 |
| **Ighm** | Immunoglobulin heavy constant mu OS=Mus musculus OX=10090 GN=Ighm PE=1 SV=2 | 1,51 | 0,0066 |
| **Gjb6** | Gap junction beta-6 protein OS=Mus musculus OX=10090 GN=Gjb6 PE=1 SV=1 | 1,51 | 0,0154 |
| **Acot1** | Acyl-coenzyme A thioesterase 1 OS=Mus musculus OX=10090 GN=Acot1 PE=1 SV=1 | 1,50 | 0,0111 |
| **Scg3** | Secretogranin-3 OS=Mus musculus OX=10090 GN=Scg3 PE=1 SV=1 | 1,50 | 0,0058 |
| **Crot** | Peroxisomal carnitine O-octanoyltransferase OS=Mus musculus OX=10090 GN=Crot PE=1 SV=1 | 1,49 | 0,0203 |
| **S100a13** | Protein S100-A13 OS=Mus musculus OX=10090 GN=S100a13 PE=1 SV=1 | 1,49 | 0,0055 |
| **Sema4a** | Semaphorin-4A OS=Mus musculus OX=10090 GN=Sema4a PE=1 SV=2 | 1,48 | 0,0073 |
| **Plxdc2** | Plexin domain-containing protein 2 OS=Mus musculus OX=10090 GN=Plxdc2 PE=1 SV=1 | 1,48 | 0,0267 |
| **Gltp** | Glycolipid transfer protein OS=Mus musculus OX=10090 GN=Gltp PE=1 SV=3 | 1,47 | 0,0112 |
| **Lsm6** | U6 snRNA-associated Sm-like protein LSm6 OS=Mus musculus OX=10090 GN=Lsm6 PE=1 SV=1 | 1,47 | 0,0154 |
| **Anpep** | Aminopeptidase N OS=Mus musculus OX=10090 GN=Anpep PE=1 SV=4 | 1,47 | 0,0217 |
| **Tgm2** | Protein-glutamine gamma-glutamyltransferase 2 OS=Mus musculus OX=10090 GN=Tgm2 PE=1 SV=4 | 1,47 | 0,0099 |
| **Lgmn** | Legumain OS=Mus musculus OX=10090 GN=Lgmn PE=1 SV=1 | 1,47 | 0,0076 |
| **Ppp1r1b** | Protein phosphatase 1 regulatory subunit 1B OS=Mus musculus OX=10090 GN=Ppp1r1b PE=1 SV=2 | 1,46 | 0,0053 |
| **Ca14** | Carbonic anhydrase 14 OS=Mus musculus OX=10090 GN=Ca14 PE=1 SV=1 | 1,46 | 0,0239 |
| **Ppp1r1a** | Protein phosphatase 1 regulatory subunit 1A OS=Mus musculus OX=10090 GN=Ppp1r1a PE=1 SV=1 | 1,45 | 0,0294 |
| **Krt76** | Keratin, type II cytoskeletal 2 oral OS=Mus musculus OX=10090 GN=Krt76 PE=1 SV=1 | 1,45 | 0,0137 |
| **Anxa3** | Annexin A3 OS=Mus musculus OX=10090 GN=Anxa3 PE=1 SV=4 | 1,44 | 0,0095 |
| **Ampd3** | AMP deaminase 3 OS=Mus musculus OX=10090 GN=Ampd3 PE=1 SV=2 | 1,44 | 0,0309 |
| **Commd1** | COMM domain-containing protein 1 OS=Mus musculus OX=10090 GN=Commd1 PE=1 SV=2 | 1,44 | 0,0212 |
| **Lamp1** | Lysosome-associated membrane glycoprotein 1 OS=Mus musculus OX=10090 GN=Lamp1 PE=1 SV=2 | 1,44 | 0,0229 |
| **Plp2** | Proteolipid protein 2 OS=Mus musculus OX=10090 GN=Plp2 PE=1 SV=1 | 1,44 | 0,0308 |
| **S100a16** | Protein S100-A16 OS=Mus musculus OX=10090 GN=S100a16 PE=1 SV=1 | 1,43 | 0,0131 |
| **Lasp1** | LIM and SH3 domain protein 1 OS=Mus musculus OX=10090 GN=Lasp1 PE=1 SV=1 | 1,43 | 0,0104 |
| **Nucb1** | Nucleobindin-1 OS=Mus musculus OX=10090 GN=Nucb1 PE=1 SV=2 | 1,43 | 0,0379 |
| **Scn3b** | Sodium channel subunit beta-3 OS=Mus musculus OX=10090 GN=Scn3b PE=1 SV=1 | 1,42 | 0,0348 |
| **Ccdc90b** | Coiled-coil domain-containing protein 90B, mitochondrial OS=Mus musculus OX=10090 GN=Ccdc90b PE=1 SV=1 | 1,42 | 0,0344 |
| **Ncstn** | Nicastrin OS=Mus musculus OX=10090 GN=Ncstn PE=1 SV=3 | 1,42 | 0,0200 |
| **Txndc5** | Thioredoxin domain-containing protein 5 OS=Mus musculus OX=10090 GN=Txndc5 PE=1 SV=2 | 1,42 | 0,0375 |
| **Selenow** | Selenoprotein W OS=Mus musculus OX=10090 GN=Selenow PE=1 SV=3 | 1,41 | 0,0425 |
| **Ier3ip1** | Immediate early response 3-interacting protein 1 OS=Mus musculus OX=10090 GN=Ier3ip1 PE=3 SV=1 | 1,41 | 0,0382 |
| **Cst3** | Cystatin-C OS=Mus musculus OX=10090 GN=Cst3 PE=1 SV=2 | 1,41 | 0,0262 |
| **Hba** | Hemoglobin subunit alpha OS=Mus musculus OX=10090 GN=Hba PE=1 SV=2 | 1,41 | 0,0217 |
| **Mvd** | Diphosphomevalonate decarboxylase OS=Mus musculus OX=10090 GN=Mvd PE=1 SV=2 | 1,41 | 0,0494 |
| **Scg2** | Secretogranin-2 OS=Mus musculus OX=10090 GN=Scg2 PE=1 SV=1 | 1,41 | 0,0455 |
| **Lamp2** | Lysosome-associated membrane glycoprotein 2 OS=Mus musculus OX=10090 GN=Lamp2 PE=1 SV=2 | 1,41 | 0,0271 |
| **Ctsb** | Cathepsin B OS=Mus musculus OX=10090 GN=Ctsb PE=1 SV=2 | 1,41 | 0,0264 |
| **Itgb1** | Integrin beta-1 OS=Mus musculus OX=10090 GN=Itgb1 PE=1 SV=1 | 1,41 | 0,0142 |
| **Rbp1** | Retinol-binding protein 1 OS=Mus musculus OX=10090 GN=Rbp1 PE=1 SV=2 | 1,41 | 0,0283 |
| **Gsn** | Gelsolin OS=Mus musculus OX=10090 GN=Gsn PE=1 SV=3 | 1,41 | 0,0276 |
| **Ctsd** | Cathepsin D OS=Mus musculus OX=10090 GN=Ctsd PE=1 SV=1 | 1,41 | 0,0237 |
| **Ctsa** | Lysosomal protective protein OS=Mus musculus OX=10090 GN=Ctsa PE=1 SV=1 | 1,40 | 0,0154 |
| **Gba** | Lysosomal acid glucosylceramidase OS=Mus musculus OX=10090 GN=Gba PE=1 SV=1 | 1,40 | 0,0472 |
| **Hexb** | Beta-hexosaminidase subunit beta OS=Mus musculus OX=10090 GN=Hexb PE=1 SV=2 | 1,40 | 0,0275 |
| **Cpe** | Carboxypeptidase E OS=Mus musculus OX=10090 GN=Cpe PE=1 SV=2 | 1,40 | 0,0188 |
| **Scarb2** | Lysosome membrane protein 2 OS=Mus musculus OX=10090 GN=Scarb2 PE=1 SV=3 | 1,40 | 0,0107 |
| **Stx12** | Syntaxin-12 OS=Mus musculus OX=10090 GN=Stx12 PE=1 SV=1 | 1,39 | 0,0428 |
| **Nptxr** | Neuronal pentraxin receptor OS=Mus musculus OX=10090 GN=Nptxr PE=1 SV=1 | 1,39 | 0,0447 |
| **Psme1** | Proteasome activator complex subunit 1 OS=Mus musculus OX=10090 GN=Psme1 PE=1 SV=2 | 1,39 | 0,0303 |
| **Tspan2** | Tetraspanin-2 OS=Mus musculus OX=10090 GN=Tspan2 PE=1 SV=1 | 1,39 | 0,0275 |
| **Nes** | Nestin OS=Mus musculus OX=10090 GN=Nes PE=1 SV=1 | 1,38 | 0,0332 |
| **Krt1** | Keratin, type II cytoskeletal 1 OS=Mus musculus OX=10090 GN=Krt1 PE=1 SV=4 | 1,38 | 0,0331 |
| **Pdcd5** | Programmed cell death protein 5 OS=Mus musculus OX=10090 GN=Pdcd5 PE=1 SV=3 | 1,38 | 0,0383 |
| **P4hb** | Protein disulfide-isomerase OS=Mus musculus OX=10090 GN=P4hb PE=1 SV=2 | 1,38 | 0,0390 |
| **Irs2** | Insulin receptor substrate 2 OS=Mus musculus OX=10090 GN=Irs2 PE=1 SV=2 | 1,37 | 0,0366 |
| **Irgm1** | Immunity-related GTPase family M protein 1 OS=Mus musculus OX=10090 GN=Irgm1 PE=1 SV=1 | 1,37 | 0,0451 |
| **Krt72** | Keratin, type II cytoskeletal 72 OS=Mus musculus OX=10090 GN=Krt72 PE=3 SV=1 | 1,37 | 0,0424 |
| **Tmem43** | Transmembrane protein 43 OS=Mus musculus OX=10090 GN=Tmem43 PE=1 SV=1 | 1,36 | 0,0471 |
| **Tpm4** | Tropomyosin alpha-4 chain OS=Mus musculus OX=10090 GN=Tpm4 PE=1 SV=3 | 1,36 | 0,0421 |
| **Gaa** | Lysosomal alpha-glucosidase OS=Mus musculus OX=10090 GN=Gaa PE=1 SV=2 | 1,35 | 0,0353 |
| **Tbl2** | Transducin beta-like protein 2 OS=Mus musculus OX=10090 GN=Tbl2 PE=1 SV=2 | 1,35 | 0,0414 |
| **Marcksl1** | MARCKS-related protein OS=Mus musculus OX=10090 GN=Marcksl1 PE=1 SV=2 | 1,34 | 0,0344 |
| **Tpm1** | Tropomyosin alpha-1 chain OS=Mus musculus OX=10090 GN=Tpm1 PE=1 SV=1 | 1,34 | 0,0429 |
| **Gss** | Glutathione synthetase OS=Mus musculus OX=10090 GN=Gss PE=1 SV=1 | 1,33 | 0,0354 |
| **Mlec** | Malectin OS=Mus musculus OX=10090 GN=Mlec PE=1 SV=2 | 1,33 | 0,0429 |
| **Serpinb1a** | Leukocyte elastase inhibitor A OS=Mus musculus OX=10090 GN=Serpinb1a PE=1 SV=1 | 1,32 | 0,0465 |
| **Lingo2** | Leucine-rich repeat and immunoglobulin-like domain-containing nogo receptor-interacting protein 2 OS=Mus musculus OX=10090 GN=Lingo2 PE=2 SV=1 | 1,31 | 0,0482 |
| **Prkca** | Protein kinase C alpha type OS=Mus musculus OX=10090 GN=Prkca PE=1 SV=3 | 0,78 | 0,0409 |
| **Fbxo41** | F-box only protein 41 OS=Mus musculus OX=10090 GN=Fbxo41 PE=1 SV=3 | 0,77 | 0,0467 |
| **Crocc** | Rootletin OS=Mus musculus OX=10090 GN=Crocc PE=1 SV=2 | 0,77 | 0,0489 |
| **Grin1** | Glutamate receptor ionotropic, NMDA 1 OS=Mus musculus OX=10090 GN=Grin1 PE=1 SV=1 | 0,77 | 0,0421 |

| | | | |
|---|---|---|---|
| **Erc2** | ERC protein 2 OS=Mus musculus OX=10090 GN=Erc2 PE=1 SV=2 | 0,77 | 0,0436 |
| **Grin2b** | Glutamate receptor ionotropic, NMDA 2B OS=Mus musculus OX=10090 GN=Grin2b PE=1 SV=3 | 0,76 | 0,0335 |
| **Dlgap2** | Disks large-associated protein 2 OS=Mus musculus OX=10090 GN=Dlgap2 PE=1 SV=2 | 0,76 | 0,0323 |
| **Kcnj9** | G protein-activated inward rectifier potassium channel 3 OS=Mus musculus OX=10090 GN=Kcnj9 PE=1 SV=2 | 0,75 | 0,0334 |
| **Iqsec2** | IQ motif and SEC7 domain-containing protein 2 OS=Mus musculus OX=10090 GN=Iqsec2 PE=1 SV=3 | 0,74 | 0,0345 |
| **Dlg4** | Disks large homolog 4 OS=Mus musculus OX=10090 GN=Dlg4 PE=1 SV=1 | 0,74 | 0,0171 |
| **Anks1b** | Ankyrin repeat and sterile alpha motif domain-containing protein 1B OS=Mus musculus OX=10090 GN=Anks1b PE=1 SV=3 | 0,73 | 0,0148 |
| **Slc25a31** | ADP/ATP translocase 4 OS=Mus musculus OX=10090 GN=Slc25a31 PE=1 SV=1 | 0,73 | 0,0423 |
| **Osbpl6** | Oxysterol-binding protein-related protein 6 OS=Mus musculus OX=10090 GN=Osbpl6 PE=1 SV=1 | 0,72 | 0,0463 |
| **Clcn4** | H(+)/Cl(-) exchange transporter 4 OS=Mus musculus OX=10090 GN=Clcn4 PE=2 SV=2 | 0,72 | 0,0423 |
| **Acot11** | Acyl-coenzyme A thioesterase 11 OS=Mus musculus OX=10090 GN=Acot11 PE=1 SV=1 | 0,72 | 0,0330 |
| **Kcnd2** | Potassium voltage-gated channel subfamily D member 2 OS=Mus musculus OX=10090 GN=Kcnd2 PE=1 SV=1 | 0,72 | 0,0448 |
| **F13a1** | Coagulation factor XIII A chain OS=Mus musculus OX=10090 GN=F13a1 PE=1 SV=3 | 0,71 | 0,0282 |
| **Gsg1l** | Germ cell-specific gene 1-like protein OS=Mus musculus OX=10090 GN=Gsg1l PE=1 SV=2 | 0,71 | 0,0360 |
| **Mlf2** | Myeloid leukemia factor 2 OS=Mus musculus OX=10090 GN=Mlf2 PE=1 SV=1 | 0,70 | 0,0092 |
| **Lysmd2** | LysM and putative peptidoglycan-binding domain-containing protein 2 OS=Mus musculus OX=10090 GN=Lysmd2 PE=1 SV=2 | 0,68 | 0,0156 |
| **Smarca4** | Transcription activator BRG1 OS=Mus musculus OX=10090 GN=Smarca4 PE=1 SV=1 | 0,68 | 0,0183 |
| **Prodh** | Proline dehydrogenase 1, mitochondrial OS=Mus musculus OX=10090 GN=Prodh PE=1 SV=2 | 0,65 | 0,0093 |
| **Cbln2** | Cerebellin-2 OS=Mus musculus OX=10090 GN=Cbln2 PE=1 SV=1 | 0,64 | 0,0477 |
| **Chd5** | Chromodomain-helicase-DNA-binding protein 5 OS=Mus musculus OX=10090 GN=Chd5 PE=1 SV=1 | 0,63 | 0,0322 |
| **Gatad2b** | Transcriptional repressor p66-beta OS=Mus musculus OX=10090 GN=Gatad2b PE=1 SV=1 | 0,63 | 0,0322 |
| **Pkn1** | Serine/threonine-protein kinase N1 OS=Mus musculus OX=10090 GN=Pkn1 PE=1 SV=3 | 0,62 | 0,0264 |
| **Rbfox1** | RNA binding protein fox-1 homolog 1 OS=Mus musculus OX=10090 GN=Rbfox1 PE=1 SV=3 | 0,61 | 0,0020 |
| **Tmem186** | Transmembrane protein 186 OS=Mus musculus OX=10090 GN=Tmem186 PE=1 SV=2 | 0,59 | 0,0196 |
| **Ube2h** | Ubiquitin-conjugating enzyme E2 H OS=Mus musculus OX=10090 GN=Ube2h PE=1 SV=1 | 0,56 | 0,0017 |
| **Slc25a44** | Solute carrier family 25 member 44 OS=Mus musculus OX=10090 GN=Slc25a44 PE=1 SV=1 | 0,56 | 0,0021 |
| **Cldn10** | Claudin-10 OS=Mus musculus OX=10090 GN=Cldn10 PE=1 SV=2 | 0,52 | 0,0005 |
| **Spata2l** | Spermatogenesis-associated protein 2-like protein OS=Mus musculus OX=10090 GN=Spata2l PE=1 SV=1 | 0,50 | 0,0001 |
| **Dnajb3** | DnaJ homolog subfamily B member 3 OS=Mus musculus OX=10090 GN=Dnajb3 PE=2 SV=1 | 0,34 | 0,0000 |
| **Nr2f1** | COUP transcription factor 1 OS=Mus musculus OX=10090 GN=Nr2f1 PE=2 SV=2 | 0,10 | 0,0000 |
| **Polr2c** | DNA-directed RNA polymerase II subunit RPB3 OS=Mus musculus OX=10090 GN=Polr2c PE=1 SV=2 | 0,01 | 0,0000 |
| **Smyd3** | Histone-lysine N-methyltransferase SMYD3 OS=Mus musculus OX=10090 GN=Smyd3 PE=2 SV=1 | 0,01 | 0,0000 |

178 elements included exclusively in injury vs. Control

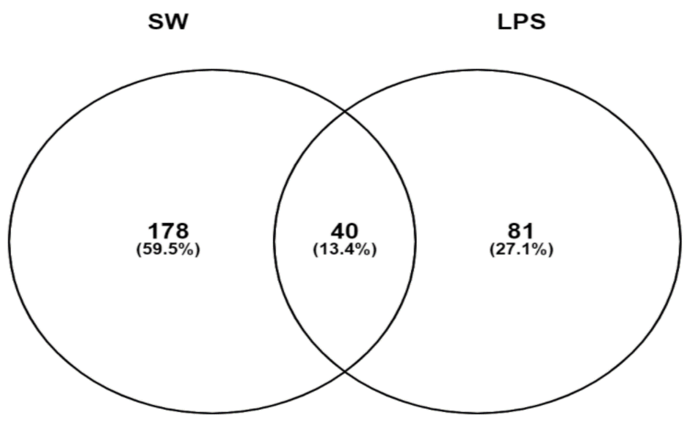
